# Dynamic shifts between pattern separation and reinstatement in human CA3 and dentate gyrus

**DOI:** 10.64898/2026.04.24.720552

**Authors:** Helena M. Gellersen, Loris Naspi, Xenia Przygodda, Emrah Düzel, David Berron

## Abstract

Distinguishing similar memories is thought to require hippocampal pattern separation, the orthogonalization of overlapping neural representations. Here, high-resolution 7-Tesla fMRI during one-shot encoding of similar target-lure pairs demonstrated that pattern separation and reinstatement are modulated by stimulus domain, similarity, and hippocampal long-axis position. Univariate novelty responses during successful object and scene discrimination were found across medial temporal lobe cortex and hippocampal subfields. Multivariate neural pattern similarity was reduced for correctly rejected low-similarity lures across domains throughout all medial temporal regions, consistent with pattern separation. Increasing target-lure similarity resulted in more hippocampal than cortical pattern separation. An exception were high-similarity scene lures which elicited greater hippocampal pattern similarity, indicating reinstatement. In dentate gyrus and CA3, reduced pattern similarity predicted better object, increased similarity better scene memory, depending on hippocampal long-axis position. Pattern separation and reinstatement are therefore not fixed computational properties of specific subregions but shift dynamically during fast episodic learning.

## Introduction

A fundamental challenge of our memory system is to create unique representations of similar experiences after a single exposure ^1–3^. Early theories such as the Complementary Learning Systems account proposed that the medial temporal lobe (MTL) cortex and the hippocampus contribute to this process via different neural computations ^4–9^. Extra-hippocampal regions are thought to operate with graded strength-based signals that carry information about familiarity of a stimulus ^4,5,10,11^. Conversely, the dentate gyrus (DG) and proximal aspect of cornu ammonis 3 (CA3) of the hippocampus engage in *pattern separation*, a process that orthogonalizes similar inputs into dissimilar neural representations ^10,12–14^. Distal CA3 is thought to support memory reinstatement via *pattern completion* upon presentation of a partial cue ^13,15–18^. Electrophysiological and functional neuroimaging studies have provided empirical support for this account in humans ^19–25^. However, more recent work has argued for additional cortical contributions to pattern separation ^26,27^. Previous methodological limitations have left important questions unanswered.

Electrophysiology cannot simultaneously sample the many heterogeneous MTL subregions and is restricted to non-human animals or human clinical populations ^22,28^. Functional magnetic resonance imaging (fMRI) overcomes these limitations, but cannot measure individual neuronal responses ^29^. How can we bridge these micro- and macroscale views of the MTL? A promising approach is the combination of ultra-high resolution fMRI with multivariate analysis methods that examine voxel-wise activity patterns to determine whether a given brain region reduces neural similarity for two stimuli. 7-Tesla imaging can provide a fine-grained view of hippocampal processes where a single voxel at 1mm resolution contains roughly 60,000 granule cells rather than the 200,000 neurons in standard 3T scanners ^19,30^. To our knowledge, no prior work has leveraged the combination of these methods to study MTL processes during one-shot learning as a function of stimulus similarity at scanner resolutions capable of distinguishing CA3 and DG.

Prior univariate fMRI studies during mnemonic discrimination tasks of similar target-lure pairs have found that the hippocampus, and particularly the combined CA3/DG subregion, treat lures similar to novel stimuli. This is interpreted as an indicator of pattern separation ^31–37^. Previous multivariate mnemonic discrimination studies do indeed point to orthogonalized hippocampal patterns but were unable to resolve CA3 and DG signals ^38,39^. The link between CA3/DG novelty signals and pattern separation during one-shot learning thus remains elusive (but see ^19^ for repeated learning). Here, we combine the complementary advantages of univariate and multivariate methods to test whether strength-based novelty responses and decorrelated neural representations converge within the same neural circuits. Without multivariate approaches that characterize voxel-wise activity patterns throughout MTL regions it would not be possible to ascertain whether neural representations of similar stimuli do indeed become more orthogonalized in the DG during one-shot learning - just as they do over repeated episodes^19,20,23,40–43^. Conversely, univariate approaches that measure response magnitude can identify novelty signals that are not expressed through decorrelated neural representations and to which multivariate methods would be blind ^44,45^. This powerful combination of analytical methods and cutting-edge fMRI allowed us to test the core predictions of the most influential theories on how intricate MTL circuits resolve feature interference during learning after a single exposure^5,8,11,26,27^.

We test whether univariate novelty signals and representational measures of pattern separation converge in DG and CA3, and how stimulus similarity and content shape computations across the hippocampal long axis and MTL cortex. We first tested the prediction that DG in particular is a key region for pattern separation and explored whether uni- and multivariate signatures of successful mnemonic discrimination converge or diverge. This is crucial to confirm whether univariate mnemonic discrimination studies are correct in interpreting their findings as indicative of pattern separation. To confirm the direct behavioral relevance of CA3 and DG signals, we also tested whether stronger lure-related novelty responses and lower pattern similarity predict successful lure discrimination.

A second goal was to determine how neural representations change as target-lure similarity increases. Models of MTL computations suggest that cortical regions can distinguish stimuli with limited feature overlap. Increasing interference is thought to require rapid hippocampal learning through pattern separation in the DG, and possibly proximal CA3 ^5–7,9,46^. Studies of repeated learning do indeed show that CA3/DG neural patterns for similar events become progressively more decorrelated over time and that this process is modulated by feature overlap ^19,20,23,40–43^. Equivalent evidence for single-episode mnemonic discrimination is lacking. We hypothesized that hippocampal regions, particularly DG and CA3, should maintain stable activation and distinct neural representations even at higher levels of feature overlap compared to cortical regions.

Third, we tested whether hippocampal pattern separation is domain general. It has been argued that hippocampal pattern separation and completion operate in the same manner across object-level and spatio-contextual information, while cortical regions have domain preferences ^5–7,10,11,27,47,48^. Yet, limitations of univariate studies ^32,36^ and the use of different task conditions and/or insufficient scanner resolution in multivariate studies ^19,20,40–43^ ^21,38,39,49^ made it difficult to assess whether this view holds. To remedy this, we used mnemonic discrimination tasks for objects and scenes under the same encoding and retrieval conditions ^32^. This allowed us to identify domain-general and -specific contributions of strength-based and pattern separation signals in hippocampal subfields and beyond.

Lastly, we tested accounts of hippocampal long-axis heterogeneity ^50–55^. It is currently unclear how this long-axis organization shapes pattern separation because no study has systematically investigated response magnitude and representational content of anterior vs. posterior subfields ^19,31,32,34,36,56,57^. 7-Tesla imaging allows us to reliably separate signals of all major subfields along the anterior-posterior axis. Based on prior work ^50,58,59^, we expected that posterior subfields maintain higher univariate activity and form more distinct neural patterns for lures.

The unique combination of ultra-high resolution imaging alongside univariate and multivariate fMRI analysis methods enable us to gain mechanistic insight into human hippocampal pattern separation and reinstatement during one-shot learning. We offer an in-depth investigation into how feature overlap and informational content shape MTL representations. Thus, we present the most fine-grained view of the contributions of hippocampal subfields and extra-hippocampal MTL regions during one-shot learning currently available *in vivo* in healthy humans. These insights have important methodological and translational implications for how we approach the study of pattern separation with an indirect method such as fMRI and for our understanding of how damage to different MTL regions may result in memory impairment under specific task conditions, aging, and Alzheimer’s disease ^32,60–62^.

## Results

Participants completed mnemonic discrimination tasks of objects and scenes (Figure 1A-C; see Methods for details) in a Siemens Magnetom 7T Plus MRI scanner. Stimulus sequences began with two novel targets followed by either repeats or similar lures, which had to be judged as old or new. Scrambled images served as baseline. Performance was measured as corrected hit rate (hits - false alarms; Supplementary Figure 1 and Supplementary Tables 1-2).

**Figure 1.**
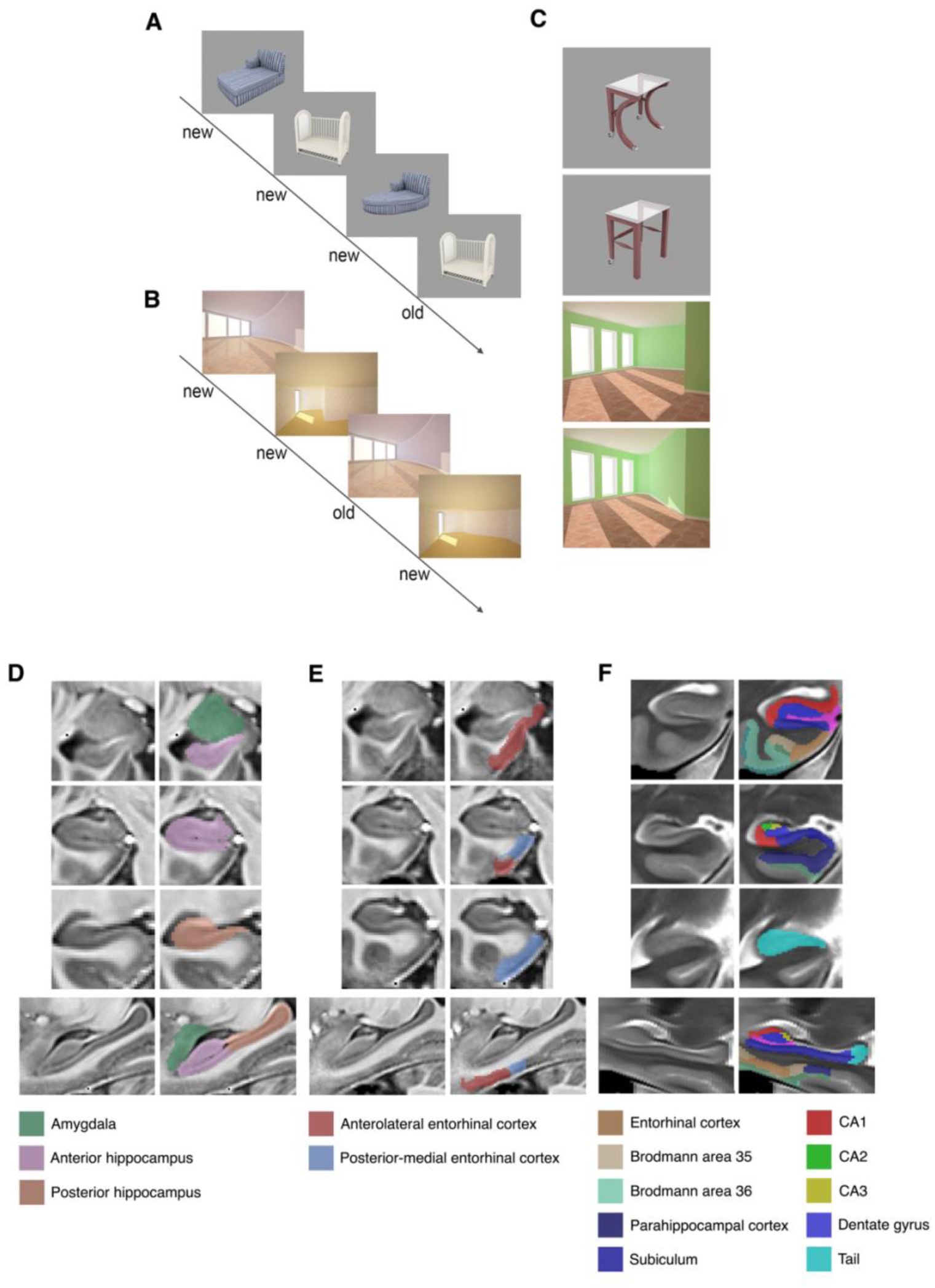
Memory tasks and medial temporal lobe subregions of interest. A. Examples of an object and B. a scene mnemonic discrimination sequence. C. Target-lure pair examples (adapted from Berron et al. with permission ^32^). D. Amygdala, anterior and posterior hippocampal masks (according to protocols in ^67,68^). E. Anterior-lateral and posterior-medial subregions of the entorhinal cortex (according to ^52,63^). F. Automatically segmented regions of interest based on a multimodal study template using T1- and T2-weighted images. The same regions were manually delineated for each participant (according to ^64^). Note that instead of the entorhinal mask shown in F., we used the entorhinal subregions from E.

Images were analysed using a standard preprocessing pipeline in SPM12. We modeled scrambled images and first presentations, repetitions, correct-lure, and incorrect-lure trials separately for objects and scenes in a first-level GLM. Anterior-lateral entorhinal cortex (alERC), posterior-medial ERC (pmERC) ^63^, Brodmann Area 35 (BA35, i.e. perirhinal cortex), BA36 (ectorhinal cortex, often labelled perirhinal elsewhere), PHC, subiculum (SUB), cornu ammonis 1 (CA1), CA3, dentate gyrus (DG), and the hippocampal tail (TAIL) were manually delineated ^64^. We split hippocampal subfields into anterior and posterior regions (prefixes ‘a’ and ‘p’, respectively) and individually analyzed left and right homologues given prior findings on hemispheric differences in category selectivity ^65,66^.

### CA3 and DG contribute to successful lure discrimination via novelty signals, pattern separation and pattern reinstatement

We first employed methods used by prior studies with univariate analysis approaches to determine whether we could replicate common findings on MTL involvement during mnemonic discrimination. ^3,11,31,32,36,61,62,69–71^ These studies argued that the following profile indicates pattern separation-like responses profiles: 1) higher activity for first than repeated targets (repetition suppression), 2) higher activity for correct lures compared to repeats (lure-related novelty), 3) no significant difference between first targets and lure stimuli (no suppression to feature overlap, the lure is treated as a novel stimulus), and 4) greater activity during correct vs. incorrect lures. This approach is essential because a single contrast cannot distinguish whether lures are represented as truly novel events ^19,72^. To test for such response profiles, we extracted *β*-estimates for each trial type defined in the first-level general linear model in native space ^31,32,36^. We fitted a linear mixed model with fixed effects of trial type and region of interest (ROI) and a random intercept per subject to the trial-level *β*-estimates. Unless otherwise specified, all analyses presented are FDR-corrected for multiple comparisons at the contrast and stimulus type level.

The linear mixed model revealed a significant ROI by trial type interaction (*F*(232,7993.1) = 8.80, *p* < .001, *η²* = .20). Figure 2A shows the results of all pairwise comparisons of interest described above (statistics in Table S3). For objects, only right pCA3, right pDG, and left pDG fulfilled all criteria for pattern separation-like responses (*p_FDR_*<.003, *d*>.9). Exploratory analysis across bilateral ROIs also found this pattern for alERC, amygdala, and posterior subfields (Figure S2). For scenes, right pDG met all criteria except higher activity for correct than incorrect lures (*p_FDR_* < .03, *d* < .85).

**Figure 2.**
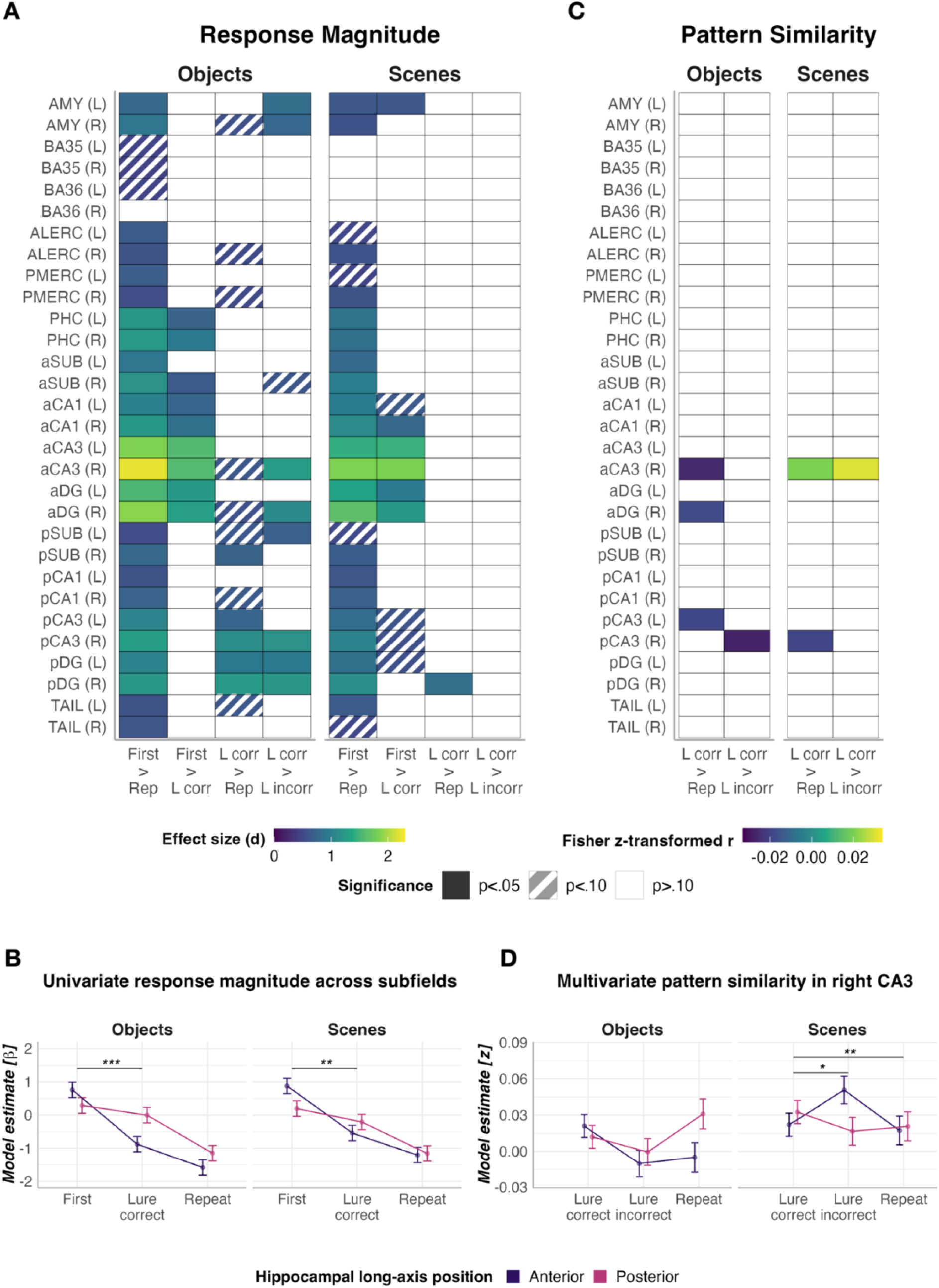
Characterizing response patterns across trial types provides insights into the nature of information processing in medial temporal lobe regions. A. Univariate *β-*estimates per trial type. Colored tiles show statistically significant pairwise comparisons between individual trial types. Comparisons were derived from estimated marginal means ^31,32^. B. Follow-up analysis to A. assessing effects of hippocampal long axis. Asterisks refer to post-hoc tests for an interaction comparing anterior and posterior hippocampus in terms of the difference in activity for first vs. correct lures and repeats vs. correct lures, therefore denoting differences in slopes. C. Multivariate analysis on neural pattern similarity estimated for each trial type relative to the first target presentation. D. Follow-up analysis to C. assessing the effects of hippocampal long axis position on right CA3 pattern similarity. The asterisks denote the differences in slope for the two-way interaction of trial type and hippocampal long axis. Notes. Prefix ‘*a’*: anterior hippocampus; ‘*p’*, posterior hippocampus. ALERC: anterior-lateral entorhinal cortex; AMY: amygdala; BA: Brodmann area; CA: cornu ammonis; DG: dentate gyrus; L: left; L corr: correct lure rejection; L incorr: lure false alarm; PHC: parahippocampal cortex; PMERC: posterior-medial entorhinal cortex; R: right; Rep: repeated target; SUB: subiculum. **p<.01, ***p<.001.

The univariate results suggested differences in anterior and posterior subfield response profiles: anterior subfields do not treat lures like novel stimuli, despite exhibiting lure-related novelty responses (Figure 2A). A follow-up analysis confirmed a significant trial type by ROI interaction (*F*(2, 318.01) = 11.31, *p* <. 001, *η²* = .07; Figure 2B), pointing to steeper decline in activity from first presentations to correct lures in anterior compared to posterior hippocampus for objects (*t*(318) = 3.73, *p_FDR_* = .001) and scenes (*t*(318) = 2.83, *p_FDR_* = .015). Lure-related novelty responses were matched (*p_FDR_* > .2). Prior univariate fMRI studies would argue that this posterior hippocampal response profile is in line with pattern separation, whereas that in the anterior hippocampus may suggest that lures are not treated like a novel stimulus ^31,32,34^.

So far, these univariate analyses suggest that hippocampal subfields and extra-hippocampal MTL can exhibit similar strength-based novelty signals during correct mnemonic discrimination. To test whether the underlying mechanism is indeed representational orthogonalization, we turn to multivoxel pattern analysis ^44,45^. We computed voxelwise similarity between the first target presentation of a given stimulus and its subsequent repetition or corresponding lure. Pattern similarity values are therefore relative to the first target presentation. We tested whether correctly rejected lures are less similar to the target than exact repetitions and incorrect lures. Reduced neural pattern similarity for correct lures suggests that successful mnemonic discrimination is associated with more distinct neural representations - a prerequisite for pattern separation.

We used a linear-mixed effects model with trial and stimulus type for all MTL ROIs (Figure 2C; statistics in Table S4). For objects, correct lures had significantly lower pattern similarity than repeated targets in right aCA3 (*z* = –4.23, *p_FDR_* = .001), right aDG (*z* = –2.84, *p_FDR_* = .045), and left pCA3 (*z* = –3.06, *p*_FDR_ = .034). Correct lures showed reduced similarity compared to incorrect lures in right pCA3 (*z* = –3.81, *p*_FDR_ = .004).

For scenes, right pCA3 showed lower pattern similarity during correct lures compared to repeats (*z* = –2.99, *p_FDR_* = .041). Surprisingly, right aCA3 showed the opposite effect to that observed in the object task and had greater pattern similarity for correct lures than repeats (*z* = 3.12, *p_FDR_* = .041) and incorrect lures (*z* = 4.13, *p_FDR_* = .001). Rather than signaling pattern separation, this could indicate that successful lure discrimination in aCA3 engages pattern reinstatement of the original target trace.

In an exploratory analysis, we followed up these CA3 findings with a linear mixed model and found a significant three-way interaction between trial type, stimulus domain and hippocampal long-axis (*F*(2,7508.4) = 5.02, *p* = .007). As expected from the patterns in Figure 2D, there was a significant difference in the lure correct vs. repeat contrast between anterior and posterior CA3 for scenes (*z* = -2.82, *p_FDR_* = .005). This difference was significantly larger compared to that in the object task (*z* = -2.88, *p_FDR_* = .004). Similarly, the difference in neural pattern similarity between incorrect and correct scene lures in aCA3 was also significantly greater than in pCA3 (*z* = 2.01, *p_FDR_* = .044). These results were driven by the increase in neural pattern similarity in aCA3 during correct lures, providing further support that correct mnemonic discrimination in the scene task may be associated with pattern reinstatement in anterior and pattern separation in posterior CA3.

The results thus far confirm the involvement of both hippocampal and extra-hippocampal MTL subregions in successful mnemonic discrimination. They also suggest mechanistic differences in how these regions overcome feature interference, pointing to a potential special role for CA3 and DG for the encoding of similar representations into more differentiated neural patterns. However, subsequent analyses suggest that this view is too simplistic. Specifically, we followed up on the ROI-based analysis with a voxel-wise approach which allows us to directly examine the spatial correspondence between uni- and multivariate effects. ROI analyses summarized neural pattern similarity and novelty signals into a single value per region, identifying areas most strongly involved in minimizing interference. Thus, they may miss weaker, localized effects. To identify such contributions, we conducted univariate cluster-level and multivariate voxel-wise searchlight analyses, focusing on the contrasts correct lures vs. repeats and correct vs. incorrect lures. For completeness, we also estimated repetition suppression effects for the univariate contrasts, which were prominent throughout the MTL (Figure S3).

For both objects and scenes, the searchlight found that voxels in all MTL regions had significantly reduced neural similarity for correct lures compared to repeats (Figure 3A-B). This contrast to our ROI-based analysis is likely due to the searchlight prioritizing spatial extent and not voxelwise intensity, which renders it more sensitive to widespread, subtle effects.

**Figure 3.**
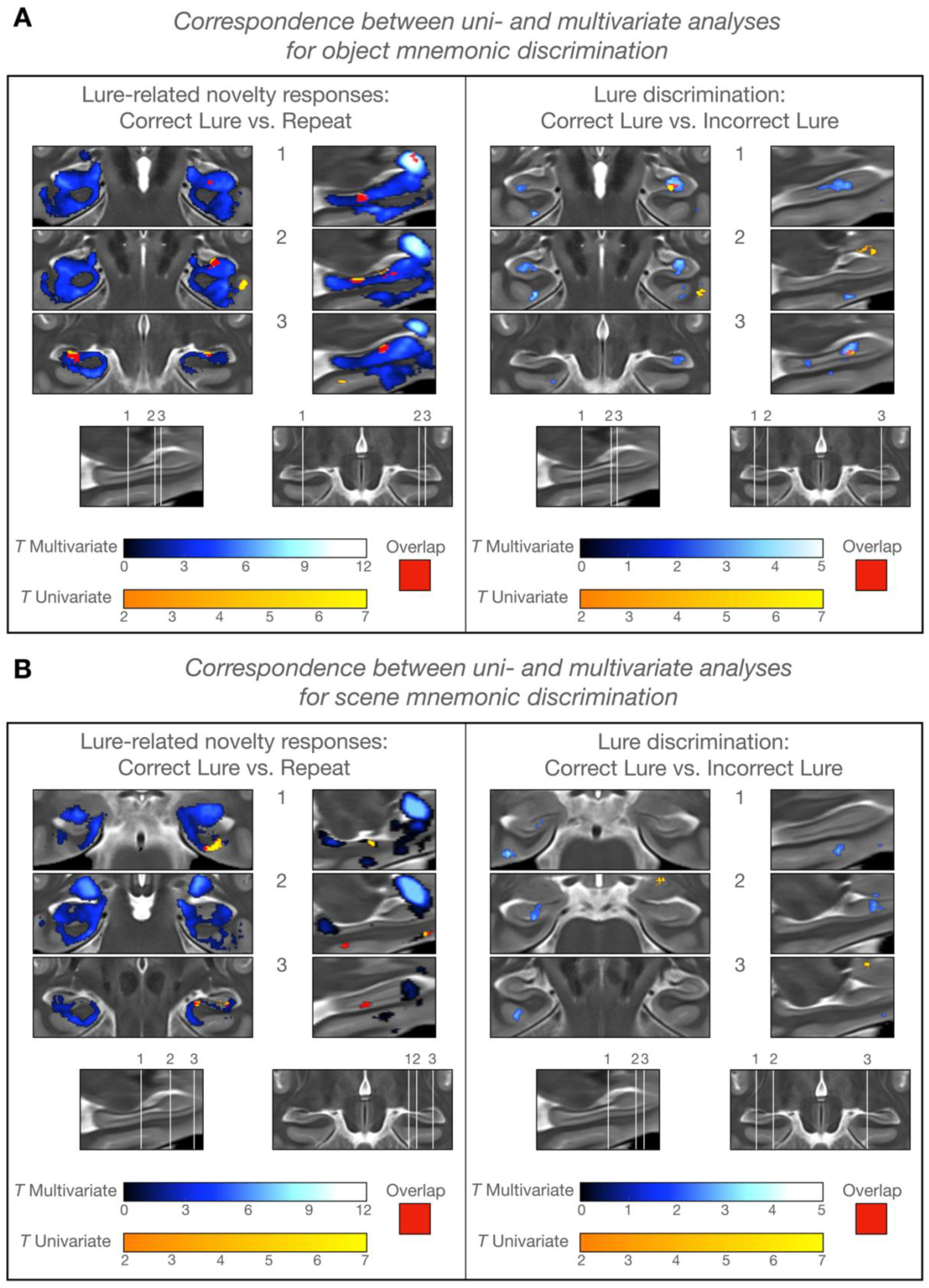
Voxelwise analyses to compare brain activity during correct lures vs. repeats and correct vs. incorrect lures. Univariate effects reflect differences in overall response magnitude between two conditions of interest. Multivariate effects reflect differences in pattern similarity between conditions. A. Effects for object mnemonic discrimination. B. Effects for scene mnemonic discrimination.

For objects, the univariate analysis found smaller clusters for the correct lures vs. repeats contrast in bilateral amygdala, pDG and pCA3, right BA36, PHC, aDG, and aCA3, and left BA35, pSUB, and TAIL (activation profiles per trial type in Figures S4-5). In the correct versus incorrect lures contrast, univariate clusters were found in bilateral amygdala, right BA36, aDG, and the ERC-subiculum junction (Figure 3A-B). The searchlight revealed that reduced pattern similarity for correct compared to incorrect object lures was most prominent in CA3 and DG throughout the hippocampus and the right TAIL. Smaller clusters were located in lateral ERC neighboring BA35. Importantly, the only spatial overlap between uni- and multivariate effects was in right aDG, specifically the same region that also showed an overlap of uni- and multivariate effects for the correct lures vs. repeats condition. This suggests a direct relationship between novelty signals and pattern separation.

For scenes, univariate clusters of lure-related novelty responses were exclusively right lateralized in BA35, PHC, pSUB, pCA1, and pDG. For the contrast of correct and incorrect scene lures, the univariate analysis only found clusters in right amygdala and BA36, while lower pattern similarity for correct lures was found in left amygdala, BA36, PHC, aSUB, aCA1, and aDG, and right alERC. Univariate and multivariate effects showed less spatial overlap than in the object task.

Lastly, earlier mnemonic discrimination work collapsed across all lure stimuli regardless of behavioral output ^31,34^. Exploratory analyses that compared repeats to all lures found substantially smaller clusters of lower pattern similarity and novelty responses (Figures S6-7). This suggests that successful lure recognition, not just perceptual overlap, is the driver behind these widespread decreases in pattern similarity.

These results implicate all hippocampal subfields, the amygdala, and MTL cortex in successful mnemonic discrimination and point to a partial divergence between uni- and multivariate effects. Notably, the most consistent spatial overlap between these methods was found in the DG, highlighting this region as a likely locus of pattern separation during object mnemonic discrimination.

### Target-lure similarity differently modulates response profiles across medial temporal lobe and hippocampal subregions

In everyday life, events range vastly in terms of the degree of similarity. Yet, so far we treated all lures equally, which has the advantage of greater statistical power but may obscure important inter-regional differences with regards to the ability to disambiguate different degrees of representational overlap ^73^. Regions capable of pattern separation may maintain similar activity levels between first presentations and similar lures with low to moderate, and potentially high, feature overlap. We therefore classified lures into three similarity bins based on difficulty levels obtained from an independent lifespan sample ^74^ and conducted a model fitting analysis based on univariate trial-level estimates for first targets, low, moderate, and high similarity lures, and repeats.

Across stimulus domains, activity in the amygdala, pmERC, and most anterior hippocampal subfields was best characterised by models that included terms for strong activity suppression to all lures and repeats (Figure 4 for select examples; Figure S8, Table S5 for model selection details; Figure S9 for voxelwise analysis). No posterior hippocampal subfield was best fit by these models.

**Figure 4.**
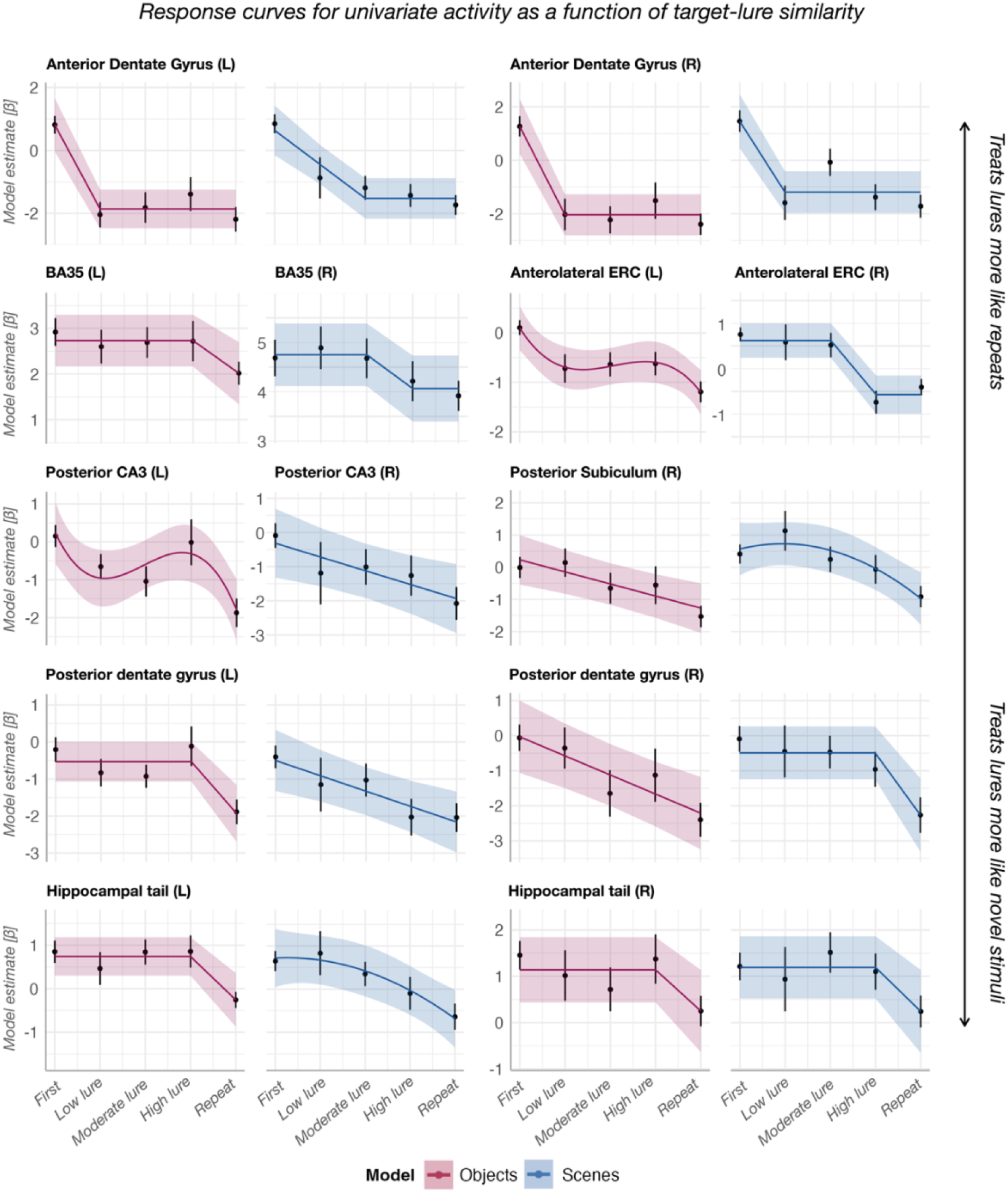
Results from model fitting of ROI-wise univariate activity across all trial types when considering three levels of lure similarity based on memory performance from an independent lifespan sample. From top to bottom row, regions are progressively less prone to activity suppression in the face of stimulus similarity. Regions with strong activity suppression in response to stimulus similarity are characterized by either a cubic fit or a model in which all lures and repeats are coded as belonging to the same condition, thus eliciting a sharp drop in response even to small degrees of feature overlap (e.g. aDG). Regions without activity suppression in response to feature overlap are characterized by similar activity from first to high similarity lures followed by activity suppression for the repeat. Point estimates and error bars represent the raw trial-level *β*-values averaged across participants and their 95% confidence interval. The fitted line represents the predicted values and their 95% confidence interval for the model that best describes brain activity patterns for a given region. Models with the lowest Akaike Information Criterion were chosen.

For the object task, left BA35, left pDG, and TAIL (disproportionately occupied by DG ^75^) were best fit by a model that assumed stable response magnitude for first, low, moderate, and high similarity lures followed by activity suppression during repeats. Left pCA3 showed slight activity suppression during low and moderate similarity object lures, followed by an increase in activity during the high similarity lure. This may reflect a repulsion effect whereby highly similar stimuli are encoded as more distinct to the corresponding target than a low similarity lure ^23,42,76^. However, caution is warranted here with regards to this interpretation given the risk of potential overfitting for cubic trends.

For scenes, right BA35, right BA36, right alERC, and left pSUB were best fit by models where low to moderate similarity lures had similar activity levels to first targets, while right pDG and right TAIL treated high similarity lures like novel stimuli. Posterior CA3 was best fit by a linear term tracking similarity.

These results further strengthen the evidence for different response profiles in the two hemispheres and along the longitudinal hippocampal axis, with anterior hippocampal activity exhibiting substantial activity suppression to any degree of feature overlap. The right hemisphere contained more regions resistant to spatial, the left hemisphere for object-level feature interference. In line with our predictions, pDG and TAIL treated even highly similar lures more like a novel than a repeated stimulus, which is consistent with pattern separation accounts. The trends in BA35, BA36, and alERC suggest that this behavior is not entirely unique to the hippocampus.

Following from the univariate analysis, we aimed to determine whether target-lure similarity differently modulates pattern similarity in hippocampal and extrahippocampal regions. According to the pattern separation account, DG is necessary for disambiguating stimuli with substantial feature overlap when cortical representations are insufficiently distinct ^7,77^. Our earlier searchlight results of cortical pattern separation may thus reflect effects from lower similarity stimuli. To test this possibility, we conducted the same searchlight analysis above separately for each lure bin. Given our *a priori* hypothesis, we tested for lower pattern similarity during correct compared to incorrect lures using a one-tailed test (Figure 5A).

**Figure 5.**
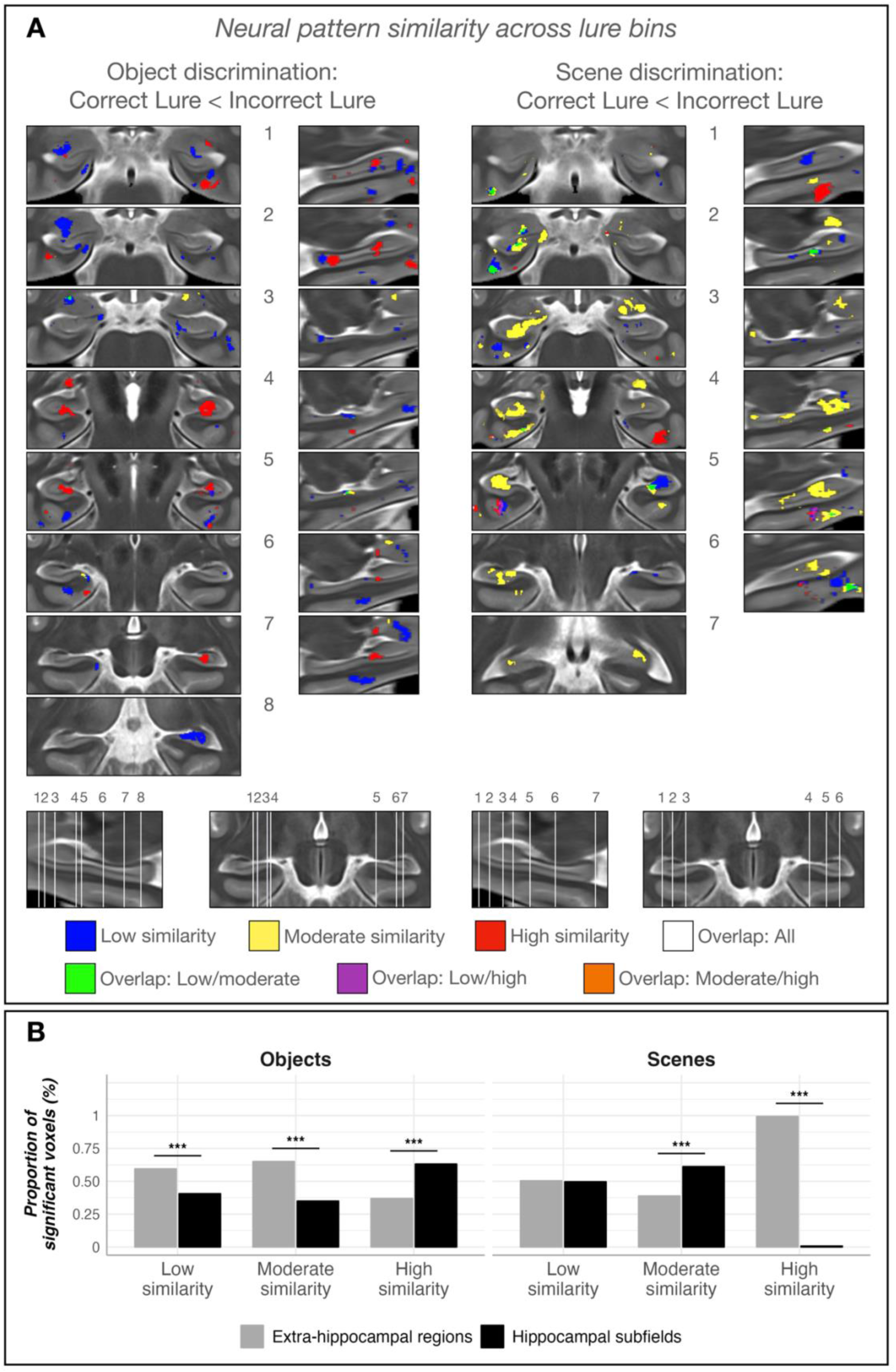
A. Differences in neural similarity between correctly and incorrectly identified low, moderate, and high ambiguity object and scene lures, respectively. Clusters indicate regions with significantly lower neural similarity between first target presentations and correct lures than between first targets and incorrect lures. B. Results of Chi-square tests comparing the proportion of hippocampal and extra-hippocampal voxels for each lure bin identified in the searchlight analysis. Voxel counts for each division were adjusted for the total number of voxels across all hippocampal and extra-hippocampal regions, respectively. \*\*\**p*<.001

For objects, multiple cortical and hippocampal regions exhibited reduced pattern similarity during correct compared to incorrect low similarity lures, including aDG and pDG. Effects for moderate similarity lures were largely absent. Lower pattern similarity for high similarity lures was also present in several cortical clusters and right pDG. The strongest effects were found in bilateral aDG and right BA35.

For scenes, lower pattern similarity during correct low similarity lures was prominent throughout cortical and hippocampal regions. The largest hippocampal clusters were found for moderately similar lures in the uncus, aDG, pDG, pCA3, pSUB and TAIL. Several clusters were also found in cortical regions. For high similarity lures, the searchlight only revealed lower pattern similarity in BA35, BA36 and ERC.

This lack of hippocampal effects for high similarity scenes was contrary to our hypothesis. However, our findings above pointed to increased lure-related neural pattern similarity as beneficial for scene memory. In our pre-planned one-tailed analysis we were unable to examine potential clusters of increased pattern similarity. We therefore conducted an exploratory two-tailed analysis (Figures S10-11). We only found higher pattern similarity during high, but not low or moderate similarity lures in left pDG and right aDG, in line with the interpretation that correct lure discrimination for high similarity scenes is more strongly driven by pattern reinstatement than separation.

Finally, we compared the proportion of above-threshold voxels located in the hippocampus versus extra-hippocampal regions for each similarity bin (Figure 5B). For objects, relatively more extra-hippocampal regions carried distinct lure representations for low (*χ²*(1) = 97.80, *p* < .001) and moderate similarity bins (*χ²*(1) = 15.06, *p* < .001). Conversely, successful discrimination of high similarity lures involved the hippocampus more than other MTL regions (*χ²*(1) = 74.54, *p* < .001), supporting the hypothesis that increasing feature interference modulates the degree to which hippocampal and extra-hippocampal regions can form distinct neural representations.

For scenes, there was no significant difference in voxel distribution for the low similarity bin (*χ²*(1) = .10, *p* = .749). The moderate similarity bin involved significantly more hippocampal voxels (*χ²*(1) = 241.29, *p* < .001), while high similarity scenes involved more cortical voxels (*χ²*(1) = 688.14, *p* < .001). This pattern is consistent with the idea that successful discrimination of highly similar scene lures may rely on greater target-lure neural similarity, possibly reflecting associative reinstatement that could support a recall-to-reject strategy ^78,79^. Such an effortful strategy may prolong reaction times. Indeed, there was an interaction between stimulus and response type (*F*(1,1921.81) = 5.93, *p* = .015), showing that correctly rejecting scene lures took significantly longer than object correct rejections (*t*(187) = -3.43, *p* =.004), whereas false alarms for both stimulus types had similar response times (*t*(230) = -.143, *p* = .889). Object and scene memory may therefore be supported by different neural processes as a function of interference.

### CA3 and DG activity and pattern similarity correlate with mnemonic discrimination performance

Our results thus far provide strong evidence that CA3 and DG are consistently involved in resolving feature interference in memory. We further strengthened these findings by examining trialwise and inter-individual brain-behavior associations.

We previously demonstrated changes in neural similarity in CA3 and DG during correct lures when averaged across all trials. Next, we sought to determine whether neural similarity in these regions reliably tracked trialwise memory performance. For each ROI, we fit a generalized linear mixed-effects model predicting memory outcome (correct or incorrect lure), using neural similarity (*z*-transformed Fisher correlation) and stimulus type as fixed effects (Figure 6A). For objects, lower target-lure neural similarity in right pCA3 marginally predicted an increased likelihood of correct rejections, indicating successful pattern separation as beneficial (*z*-ratio = - 2.43, *z-*trend=-.92, *p_FDR_*=.061, *p_uncorrected_*=.015). A similar trend was observed in right aDG (*z-*ratio= -1.65, *z-*trend=-.87, *p_FDR_*=.200, *p_uncorrected_*=.100). Conversely, for scenes, higher neural similarity was marginally associated with correct rejections, suggesting possible pattern reinstatement as beneficial (*z-*ratio= 2.50, *z-*trend=.79, *p_FDR_*=.050, *p_uncorrected_*=.013). No other region exhibited associations of trial-by-trial pattern similarity and performance (Figure S12).

**Figure 6.**
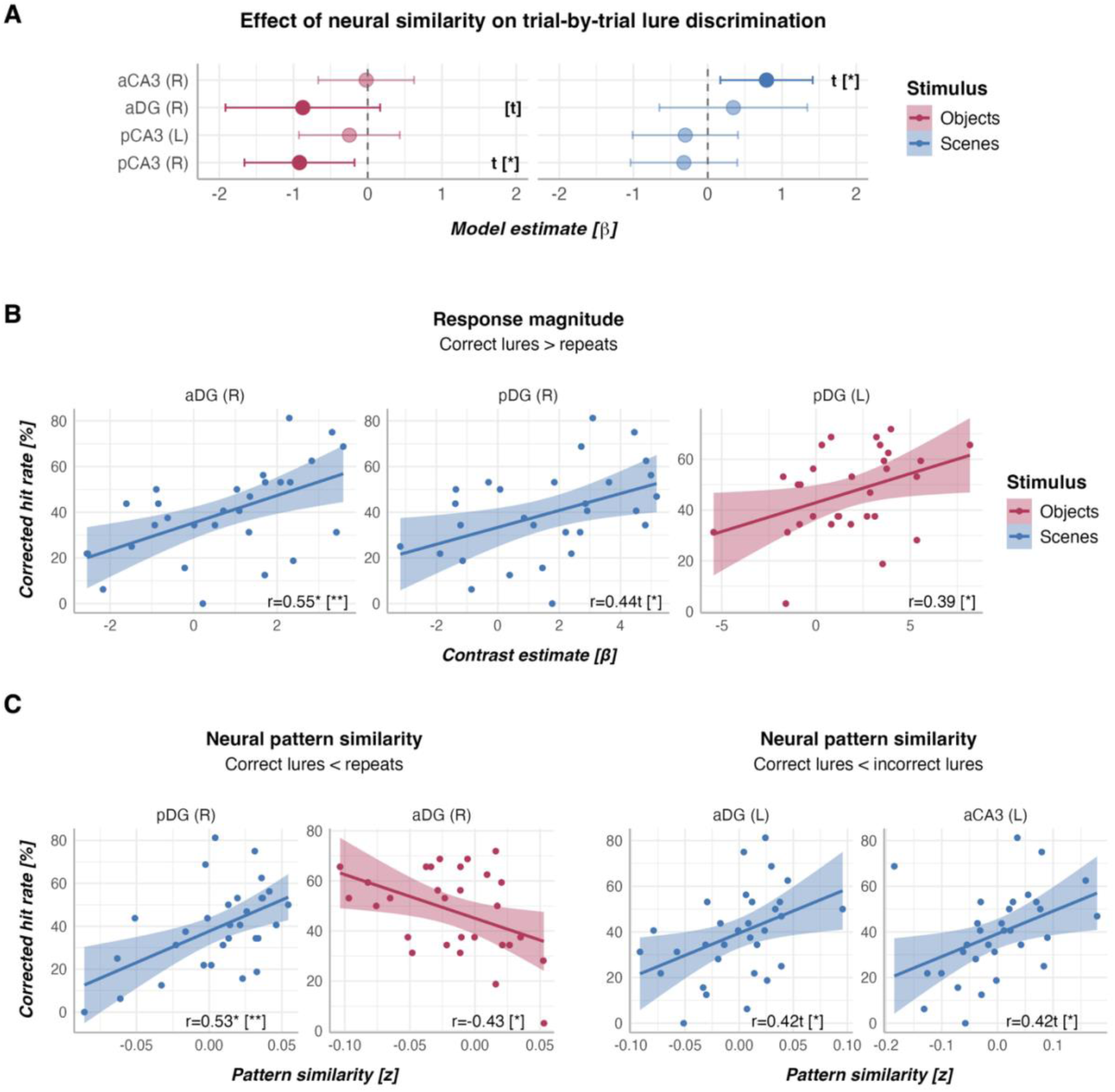
Correlations between brain responses and mnemonic discrimination performance. A. Trial-by-trial associations of target-lure neural pattern similarity and lure discrimination. Positive *β*-values suggest that increased pattern similarity between target and lure are associated with a higher likelihood of correctly accepting the lure as old. Negative *β*-coefficients suggest that lower pattern similarity between the target and its corresponding lure is associated with correct memory responses. Bars with solid colors represent effects with *p*-values at the p<.10 level. Error bars represent 95% confidence intervals. B. Correlations between univariate estimates of brain responses and inter-individual variability in mnemonic discrimination. C. Correlations between neural pattern similarity and inter-individual variability in mnemonic discrimination. The *x*-axis represents the difference in pattern similarity between the conditions. In all analyses, asterisks inside the square brackets indicate uncorrected *p*-values, asterisks outside the brackets FDR-corrected *p*-values. Brain-behavior correlations for all MTL regions are shown in the Supplementary Material. Notes. aCA3: anterior cornu ammonis-3; pCA3: posterior cornu ammonis-3; aDG: anterior dentate gyrus, pDG: posterior dentate gyrus. *^t^p*<.1, \**p*<.05, \*\**p*<.01.

We next asked whether CA3 and DG activity can explain variability in memory performance between participants (Figure 6B-C). Univariate lure-related novelty responses in right aDG correlated positively with scene mnemonic discrimination performance (*r*(27) = .55, *p_FDR_* = .017). The same correlation did not survive FDR correction for the right (*r*(27) = .44, *p_FDR_* = .065, *p_uncorrected_* = .016) or left pDG for object performance (*r*(28) = .39, *p_FDR_* = .246, *p_uncorrected_* = .034). There was no association between memory performance and univariate responses in the lure correct vs. incorrect contrast. An exploratory analysis showed significant correlations between greater repetition suppression and better memory performance in various subfields and extra-hippocampal regions (Figure S13, Table S6).

Contrary to our hypothesis of distinct hippocampal neural patterns supporting memory performance, *greater* pattern similarity in right pDG for correct scene lures than repeats correlated with better performance (*r*(28) = .53, *p_FDR_* = .021; statistics for all regions in Table S7). That is, the more similar neural activity patterns during the presentation of correct lures were to neural activity patterns during first scene target presentations, the better the mnemonic discrimination. The same was true for participants with greater pattern similarity for correct than incorrect lures in left aCA3 (*r*(29) = .42, *p_FDR_* = .080, *p_uncorrected_* = .019), and aDG, although this association was only at trend after FDR correction (*r*(29) = .42, *p_FDR_* = .080, *p_uncorrected_* = .020). These seemingly counterintuitive results echo our findings above linking better scene memory to pattern reinstatement. Interestingly, the opposite was true for the object task in right aDG. Although the correlation did not survive FDR correction, the coefficient suggests that individuals who exhibit lower pattern similarity between first and correct lure presentations compared to firsts and repeats tend to discriminate similar objects better (*r*(27) = -.43, *p_FDR_* = .169, *p_uncorrected_* = .021). This is indicative of pattern separation as the likely mechanism behind better object memory.

In an exploratory analysis, we used a model selection procedure to identify the best combination of predictors to explain mnemonic discrimination performance. Candidate predictors were uni- and multivariate measures across all MTL ROIs. Among the best predictors for object and scene models were greater univariate DG lure-related novelty responses and higher or lower SUB pattern similarity, for objects and scenes, respectively (details in Table S8).

These findings point to a direct behavioral relevance for hippocampal activation strength and pattern similarity for inter-individual variability in memory fidelity across visual domains. Interestingly, stronger univariate responses and lower neural pattern similarity for correct lures are not always better. These results caution against the interpretation of pattern separation as an exclusive mechanism for the resolution of interference in memory.

### Associations between univariate and multivariate measures of mnemonic discrimination are not consistent across task demands

Our findings revealed a divergence between univariate and multivariate signatures of mnemonic discrimination: stronger univariate lure-related responses consistently tracked successful performance, whereas both lower and higher pattern similarity were beneficial depending on stimulus domain. This suggests a complex relationship between activation strength and neural similarity. To test this directly, we correlated subject-level contrast estimates from the univariate and multivariate analyses. For each participant, we computed condition differences (correct lure − repetitions and correct lures − incorrect lures) separately for activation and neural similarity. Because this analysis tests associations between contrasts, a positive correlation would suggest that participants who show stronger activation differences between conditions also show larger neural similarity differences for the same contrast.

We focused our planned analyses on DG and CA3. For objects, we observed a significant *positive* correlation between univariate and multivariate estimates in left aDG for the correct lure versus repetition contrast (r(28) = .50, *p_FDR_*= .043). Thus, participants who exhibited larger activation differences between correct lure rejections and repetitions also showed larger differences in neural similarity between those same conditions. This suggests that univariate and multivariate measures converge in reflecting separation between memory states. In the scene condition, we observed the opposite relationship, namely a significant *negative* correlation in right pCA3 (r(28) = –.52, *p_FDR_* = .028). Here, participants with larger activation differences between conditions showed smaller neural similarity differences. The relationship between response magnitude and representational structure therefore differs depending on stimulus domain. In line with our findings above, this could hint at stronger pCA3 overall engagement possibly being linked to pattern reinstatement for scenes.

Thus, consistent with the above findings, we again observe a functional dissociation between object and scene memory in the form of different relationships between response magnitude and neural similarity. Interestingly, this trend is echoed across many other ROIs (Figure S14, Table S9). This highlights the importance of caution when interpreting lure-related univariate novelty responses as indicative of pattern separation ^11,31,32,36,62^, given that the relationship between activation strength and neural similarity depends on task demands.

## Discussion

Our findings show that pattern separation and reinstatement are not fixed properties of specific subregions but dynamic computations that shift with stimulus content and similarity, a distinction that univariate methods alone cannot resolve. Using 7T fMRI with manual hippocampal subfield segmentation, we combined univariate and multivariate analyses during one-shot mnemonic discrimination to show that CA3 and DG are central to differentiating similar events, yet engage opposing representational strategies depending on stimulus domain and stimulus similarity. Increased similarity drove pattern separation for objects but favored reinstatement for scenes. Extra-hippocampal MTL regions also carried distinct representations, though hippocampal involvement became progressively more prominent as similarity increased. Our results support the central tenets of the pattern separation account ^5,8^ but also point to extra-hippocampal contributions ^26,27^. We further highlight important methodological considerations for researchers seeking to understand MTL computations.

We first followed established methodology from univariate studies and confirmed that all MTL regions carried lure-related novelty responses for either objects and/or scenes on the ROI or the voxel level ^3,11,19,31,33–37,56,69,71,80–82^. Posterior hippocampal subfields, amygdala and alERC met all criteria for pattern separation-like response profiles as put forward by univariate fMRI studies ^31,32,34,36,62,80^ : repetition suppression, lure-related novelty responses, and similar activity for novel stimuli and lures. Posterior DG, BA35 and alERC signalled novelty even during moderate to high similarity lures. Crucially, our multivoxel analyses confirmed that some of these univariate signals overlapped with distinct neural patterns and may indeed reflect orthogonalization of similar representations. This important mechanistic link between greater activity during correct lures and more distinct representations could not be established by previous studies on one-shot episodic discrimination.

Our evidence for orthogonalized posterior CA3 scene representations and anterior DG and anterior/posterior CA3 object representations dovetail with animal studies in which DG and CA3 neurons proximal to DG remap in similar environments ^13,14,83–85^. These findings are consistent with predictions of the Complementary Learning Systems account regarding hippocampal computations ^5,8^. The finding that more hippocampal than cortical voxels exhibited distinct neural patterns for high similarity objects and moderate similarity scenes also aligns with this view ^4–6,8,9,11^: hippocampal computations become increasingly more important for mnemonic discrimination as feature overlap increases. Our results thus extend prior human work by showing that computational principles at work during repeated learning also apply to rapid memory formation ^20,23,42^. All our analyses converge to provide strong evidence that pattern separation responses are most pronounced in CA3 and DG, become progressively more crucial as stimulus similarity increases, occur across stimulus types, track trial-by-trial lure discrimination success, and explain inter-individual variability in memory performance. Interestingly, the posterior subiculum also met many of these criteria. As hippocampal-neocortical gateway and amplifier of hippocampal output signals, subicular representational fidelity, likely driven by CA3 and DG, therefore seems to be an important additional determinant for successful memory across domains ^81,86,87^ .

A surprising finding was *increased* anterior CA3 and to a lesser extent DG pattern similarity for correct lures in the high interference condition for scenes, providing further evidence that content and context dictate which hippocampal computational mode is engaged ^23,57^. Increased neural similarity suggests pattern reinstatement, a process typically considered detrimental to mnemonic discrimination due to increased false alarms ^11^. Opposing computational mechanisms across stimulus domains may reflect difficulty and/or differences in object and scene processing. High feature overlap may induce a pattern completion bias ^11,46^. Although scenes were moderately more difficult than objects (Figure S1, Table S2), confounding of stimulus domain and difficulty is too simplistic an explanation, because neural similarity correlated with better scene memory. Unlike objects ^36,87,88^, scenes involve more complex, global, less verbalizable features. This may trigger holistic hippocampal recall through the recurrent CA3 architecture ^79,86^ and support a strategy where reinstated scene features are successively compared to the presented item to detect a mismatch ^79,89^. The longer reaction times for scenes compared to objects during correct rejections hint at such an effortful strategy. Rather than reflecting failed pattern separation, increased neural similarity may thus index flexible, context-dependent shifts towards a reinstatement-based recall-to-reject approach.

The comparison of stimulus domain and similarity provides novel insights into hippocampal long axis involvement during feature resolution. Anterior CA3 and DG showed repetition suppression during low similarity lures, while posterior subfields maintained high activity across similarity bins. Nonetheless, all of these regions orthogonalized neural representations, while only anterior subfields supported reinstatement. This may be explained by the anterior-posterior hippocampal gradient in representational granularity. Posterior regions retain fine-grained detail due to smaller receptive fields ^90–92^, thus signalling novelty despite feature overlap. The anterior hippocampus encodes memories at a schematic or global level, supporting both integration and differentiation ^54,59,91^. This may explain its representational flexibility (pattern separation vs. reinstatement) and could result in activity suppression even when faced with low feature similarity. Interestingly, anterior and posterior DG response magnitude positively correlated with both object and scene memory. This suggests that strength-based responses in both regions successfully signal the presence of the lure despite differences in the underlying representations exemplified by higher pattern similarity in the anterior and lower similarity in the posterior subfield. Prior studies with lower field-strength and resolution as well as analyses collapsing subfields across the hippocampal long-axis would have missed such nuance ^19,34,61,62,93,94^.

It is important to note that our findings also revealed lower pattern similarity for correct lures vs. repeats in the amygdala and across MTL cortex in both stimulus domains. Moreover, for the correct vs. incorrect lure contrast, more extra-hippocampal than hippocampal voxels carried distinct neural representations for low similarity lures. Extra-hippocampal regions therefore do carry distributed, modestly pattern separated signals. Whether these extra-hippocampal MTL effects primarily reflect hippocampal input, pattern separated hippocampal output, and/or both is unclear given the limited temporal resolution of fMRI. Here, response magnitude rarely correlated with pattern similarity in extra-hippocampal regions, but often did so in hippocampal subfields (Figure S8), which could hint at cortical pattern similarity being shaped by hippocampal pattern similarity. This cannot be the sole explanation though because several cortical regions showed reduced pattern similarity during high similarity scene lures while the hippocampus supported pattern reinstatement (Figure S13). Prior studies argue that MTL cortex can partially separate representations through cellular mechanisms distinct from the hippocampus ^2,26,27,38,95,96^. This may explain how opposing processes can be present simultaneously in hippocampal and extra-hippocampal regions.

We made important methodological observations that expand on prior work on the relationship between response magnitude and neural representational similarity ^45,97–105^. Specifically, our findings show that uni- and multivariate proxies for pattern separation do not always align: the regional activation magnitude can reflect both pattern separation *and* reinstatement depending on task demands. Thus, both processes can lead to greater local field potentials underpinning the BOLD response ^106^. Consequently, we caution against the use of univariate approaches alone to infer pattern separation/completion or familiarity signals and suggest that combining both methods together can generate knowledge that cannot be gleaned from the use of either alone ^31,32,61^: pattern similarity did not uncover the strict anterior-vs-posterior distinction in resistance to feature overlap found here, whereas univariate approaches could not detect the simultaneous presence of pattern separation and reinstatement responses within the same subfield along the hippocampal axis.

Lastly, our similarity-bin results carry a direct implication for how mnemonic discrimination tasks could be used clinically ^3,61,107–114^. Because cortical MTL regions support pattern separation for low-similarity lures while hippocampal involvement becomes necessary as stimulus similarity increases, the gradient we describe maps onto the known progression of tau pathology which targets BA35 and alERC before the hippocampus in Alzheimer’s disease ^115,116^. Individuals with early cortical pathology should therefore already be impaired on low- to moderate-similarity discriminations. Systematically varying lure similarity in clinical paradigms could thus help to better track where along the MTL hierarchy dysfunction has spread.

Taken together, we provide strong evidence for pattern separation for memory after a single exposure. Previous fMRI studies tested overtrained stimuli and/or did not examine pattern similarity for individual stimulus pairs ^19–21,23,38,40,43,52,57,117–123^. Our study therefore extends prior cross-species findings of pattern separation ^4,5,9,11,13,18,92^ to one-shot learning and highlights the value of combining fMRI analysis methods to disentangle novelty signals, pattern separation and pattern reinstatement. CA3 and DG response magnitude and neural distinctiveness explain memory performance. Both regions flexibly engage in pattern separation and reinstatement as a function of stimulus similarity, stimulus type, and hippocampal long axis position. In addition to novelty signals, cortical MTL regions also carry distinct representations, although to a lesser extent with increasing feature overlap between stimuli. These findings provide a fine-grained understanding of MTL circuitry underpinning memory fidelity in humans and caution against reliance on univariate methods alone to infer computational mechanisms.

## STAR Methods

### EXPERIMENTAL MODEL AND STUDY PARTICIPANT DETAILS

Participants were right-handed, healthy younger adults without a history of neuropsychiatric conditions and no contraindications for 7-Tesla MRI. They gave informed consent and were compensated for their participation. The study was approved by the ethics committee of the Otto-von-Guericke University, Magdeburg, Germany (reference 128/14). Forty-one participants were recruited. The final sample with available fMRI data for analysis consisted of 34 participants (Table 1). Two were excluded due to missing imaging data and five due to excessive motion (13% or more voxels were identified as outliers). Data from participants reported here were also used in another study ^52^.

**Table 1.** Sample demographics.

|  |  |
| --- | --- |
|  | N=34<br>younger adults |
| Sex |  |
| Male | 13 (38.2%) |
| Female | 21 (61.8%) |
| Age (years) |  |
| Mean (SD) | 25.6 (4.12) |
| Median [Min, Max] | 25.0 [19.0, 35.0] |
| Corrected hit rate objects (%) |  |
| Mean (SD) | 47.8 (16.0) |
| Median [Min, Max] | 51.6 [3.13, 71.9] |
| Corrected hit rate scenes (%) |  |
| Mean (SD) | 39.2 (18.5) |
| Median [Min, Max] | 40.6 [0, 81.3] |

### METHOD DETAILS

The data presented in the current study have previously been published by Grande and colleagues in a study mapping entorhinal-hippocampal connectivity pathways ^52^.

#### Object and scene mnemonic discrimination task

The memory tasks used in the present study were originally described by Berron et al. ^32^ and are shown in Figure 1A-C. Stimuli for the mnemonic discrimination tasks were computer-generated (3ds Max, Autodesk Inc., San Rafael, USA) images of everyday indoor objects and scenes depicting empty rooms that were devoid of objects, all on gray backgrounds. Targets and lures in each pair were designed to share most features and differed only in the spatial configuration or shape of features, rather than color or patterns to avoid pop-out effects and emphasize the contribution of MTL regions ^32,124,125^. As a proxy for lure-target similarity and to match difficulty of the object and scene tasks memory performance metrics were obtained from an independent lifespan cohort of 1554 adults aged between 18 and 77 years ^74^. Based on these data, lures were classified as low (corrected hit rate, *CHR*, > .35), moderate (.15 > *CHR* <= .35), and high difficulty (*CHR* <= .15) to obtain an approximately even split (31% low, 34% moderate, 35% high similarity). Object and scene stimuli were matched by difficulty.

The task itself presented sequences of four stimuli of either objects or scenes (Figure 1A). The first two were novel, while the following could either be a repeat or a lure. For each stimulus category, 64 new targets, 32 target repeats, and 32 lures were presented for 3 seconds each, separated by a white fixation star. After the first target presentation, a noise mask was briefly shown. The following two stimuli could either be a lure-lure, repeat-repeat, repeat-lure, or lure-repeat pair. The frequency of these pairs across the task was counterbalanced. The interstimulus interval ranged from 0.6 to 4.2 seconds (mean 1.63 seconds) for an event-related design. Object and scene sequences were intermixed and their order randomized. Subjects used their right index and middle fingers to judge stimuli as “Old” or “New”. We decided not to use an Old/Similar/New format with an additional condition of novel, dissimilar foil stimuli (as in ^31^) to allow for more lure trials and maximise statistical power to detect differences between lures and repeats as well as correct vs. incorrect lures. Also, this paradigm was originally developed to test age-related decline in mnemonic discrimination and keeping to two response options was important to reduce task difficulty. Given that one of our interests is to accurately map the neural correlates of these tasks for translational, early detection work in at-risk populations, we did not alter this design in this study focused on younger adults. At the beginning and end of each of the two runs, ten scrambled images were presented to serve as a baseline. Prior to scanning, participants received task instructions and completed a 5-minute training session outside the scanner. Stimuli were displayed using the Presentation software (Neurobehavioral Systems, https://nbs.neuro-bs.com). Whether the scan would begin with object or scene stimuli was counterbalanced across participants. Stimuli were presented on an MR-compatible LCD display.

#### MRI sequences

Imaging data were acquired with a 7-Tesla Siemens MR scanner (Erlangen, Germany) with a 32-channel head coil. A whole-brain MPRAGE image was acquired first (voxel size 0.6mm isotropic, TR=2500ms, TE=2.8ms, 288 slices, interleaved acquisition, field of view 384x384x288). A T2*-weighted, partial-view image was acquired orthogonally to the long axis of the hippocampus (0.4x0.4x1mm resolution, TR=8000ms, TE=76ms, interleaved, 55 slices, FOV 512x512x55). The memory task was separated into two runs during which echo-planar images (EPI) where acquired orthogonal to the hippocampal long axis (1mm isotropic, 332 volumes per run, TR=2400ms, TE=22ms, FOV 216x216x40, interleaved).

#### Structural MRI preprocessing

The amygdala and anterior and posterior hippocampal masks were obtained from Automated Segmentation of Hippocampal Subfields (ASHS) ^68^ of the T1-weighted image using a protocol developed by Wuestefeld and colleagues ^67^ (Figure 1D). The entorhinal cortex was split into anterolateral and posterior-medial subregions using template-based masks derived from fMRI intrinsic functional connectivity (Figure 1E) ^52,63^. Other medial temporal lobe (MTL) regions comprised of BA35 (perirhinal cortex), BA36 (ectorhinal cortex), and the entorhinal (ERC) and parahippocampal cortices (PHC) as well as subiculum, CA1, CA3, the dentate gyrus (DG), and the tail of the hippocampus were manually segmented according to the Berron et al. protocol ^64^ by two experienced raters with a Dice Similarity Coefficient of .78 indicating high inter-rater reliability (Figure 1F). All masks obtained from automated segmentation were closely inspected and manually edited if necessary. In case of overlapping voxels between different segmentation methods, manually segmented hippocampal and BA35 voxels won over template-based ERC subregions and ERC subregions won over the amygdala. Hippocampal subfields were split into anterior and posterior based on the disappearance of the uncus.

All structural masks were coregistered to the EPI mean functional image using *antsRegistrationSyNQuick* and *antsApplyTransforms* ^126,127^. After inspecting image quality, segmentations, and coregistrations, 33 of the 34 participants with fMRI data passed quality control for the pmERC, alERC, amygdala, and anterior and posterior hippocampus, and 31 had MTL subregions available based on manual segmentation.

We used Advanced Normalization Tools (ANTs) to build a multivariate study-specific template from the T1- and T2-weighted images using the *antsMultivariateTemplateConstruction2.sh* command ^127^. T1- and T2-based ASHS-derived MTL masks of the template were obtained using the Wuestefeld and colleagues ^67^ and Berron et al. ^64^ protocols, respectively.

For each participant, a temporal signal-to-noise (tSNR) image was calculated from all functional volumes across the two runs. The tSNR image was masked to retain only grey matter voxels defined in the SPM T1-image segmentation step. TSNR values were *z*-scored. We used this image to correct our region of interest masks. All voxels with a tSNR-based *z*-score below -2 were removed from the mask of a given region of interest.

#### Functional MRI preprocessing

EPIs were distortion corrected based on a point-spread function and motion corrected during the online reconstruction stage as detailed in ^128^. All other preprocessing steps were conducted with SPM12 (Wellcome Department of Cognitive Neuroscience, University College, London UK) ^129^. EPIs were slice time corrected with realignment to the middle slice and smoothed with a Gaussian kernel of 2mm. The artifact detection toolbox ART ^130^ was used on the slice time corrected images to mark outliers with respect to mean image intensity and motion (threshold for global intensity: *z*>5; movement threshold: 0.3 mm, rotation threshold: 0.1 mm in accordance with Grande et al. ^52^).

### QUANTIFICATION AND STATISTICAL ANALYSIS

We set our threshold for statistical significance at a *p*-value of .05. Where multiple comparisons are made, we correct *p*-values based on the False Discovery Rate (FDR) at the stimulus (objects, scenes) and contrast level (first vs. repeats, repeats vs. correct lures, correct vs. incorrect lures) unless otherwise specified.

#### Behavioral analysis

We first tested for potential differences in memory performance between the object and scene tasks in a repeated-measures ANCOVA with stimulus category as within-subjects factors, age and sex as covariates, and corrected hit rates as outcome. In a second ANOVA with category and response type, we tested for differences in the proportion of hits and correct rejections.

### Univariate Analysis

First-level analyses were carried out in each subject’s native space. In a general linear model (GLM), we coded a total of eight regressors for object and scene stimuli: first and repeated presentations of targets, respectively, lures that were later correctly identified (correct rejections, CR), and lures that were incorrectly endorsed as targets (false alarms, FA). A ninth task-relevant regressor was added for scrambled images. We did not model target misses due to an insufficient number of trials. Six motion parameters estimated during the realignment stage and one regressor accounting for the experimental run were modelled as covariates. Additionally, each outlier volume flagged by ART was included as a separate regressor. We used the default hemodynamic response function (HRF) available in SPM12 ^129^.

Our two main contrasts of interest were “successful lure discrimination” as indexed by higher activity during correct vs. incorrect lures and “lure-related novelty responses” as indexed by higher activation during the correct rejection of a lure compared to the repeated target (see also ^32^). Note that these contrasts both indicate successful lure discrimination but provide complementary information. The former entails novelty-based discrimination, the latter holds stimulus status constant while isolating behavioral differences.

In a sensitivity analysis, we also defined a behavior-agnostic contrast which compared all lures to all repeat presentations as this approach makes use of all stimuli, meaning that the power to detect meaningful effects is independent of performance. It has also been used by prior research on the role of MTL regions in pattern separation allowing a more direct comparison with previous findings ^31^. In addition, we conducted exploratory analyses on a repetition suppression contrast that identified regions with reduced activation during the second compared to the first presentation of the target.

#### Analyses in native space

Using the *fslstats* command, we obtained trial-level *β*-estimates from the first-level GLM for each behavioral regressor (targets, repeats, correct lures, incorrect lures, and scrambled images) in each ROI in native space. This allowed us to characterize each voxel’s response across all trial types while preserving inter-individual anatomical variability between subjects. We also extracted mean *β*-estimates from all contrasts described above.

#### Analyses in template space

For our group-level analysis, we coregistered all contrast images to our group template. To do so, we used *antsApplyTransforms* with the warp field and affine transforms obtained during the ANTs template building process and the affine transform obtained from co-registering the mean functional with the T1-weighted image ^127^. Visual inspection ensured that only correctly warped images were included in the analysis. We conducted one-sample *t*-tests for the aforementioned contrasts in a second-level GLM. We used an uncorrected threshold of *p*<.001 at the voxel-level with a cluster-forming threshold set to FDR across all grey matter voxels in the EPI field of view. Active clusters were saved as binary images for further analysis and visualization with MRIcroGL ^131^. By multiplying these thresholded masks with each of the anatomically delineated regions of interest, we obtained a mask of all above-threshold voxels within each anatomically delineated ROI. This meant that continuous active clusters spanning different subregions in the hippocampus were split into separate subregions according to the anatomical definition of the ROI. We used this approach to extract trial-level *β*-estimates from each of these ROIs with *fslstats* to show descriptive statistics of response profiles of voxels identified in these contrasts of interest.

#### Modelling lure similarity

In addition to the previously described main GLM, we also conducted another analysis in which we did not split lures based on behavioral outcome for a given participant, but rather based on target-lure similarity as defined above. This analysis was motivated by prior studies which argued that researchers seeking evidence for pattern separation-like signals in fMRI studies should ideally utilize target-lure pairs spanning different degrees of feature overlap ^3,61^. In a separate first-level analysis, we therefore coded a regressor for each of low, moderate, and high similarity lures. We then extracted *b*-estimates for each trial type with *fslstats*. For exploratory purposes, we also conducted a second-level analysis in which we identified clusters with significant lure-related novelty responses to low, moderate, and high similarity lures. Note that we did not have sufficient correct and incorrect lure trials per bin in all participants, we did not compute any contrasts of correct vs. incorrect lures as a function of lure similarity.

### Multivariate Analysis

#### fMRI single-trial β-estimates

For the neural pattern similarity analyses, we re-estimated single-trial *β* maps because recent GLM toolboxes such as GLMsingle ^132^ (available at https://github.com/cvnlab/GLMsingle) offer several advantages over conventional single-trial estimation. In particular, GLMsingle combines voxelwise HRF optimization, automated noise regression via GLMdenoise ^133^ and cross-validated ridge regularization to improve the reliability of trial-level estimates. Voxel-specific HRFs account for spatial variability in hemodynamic responses, noise regressors derived from low-task-variance voxels capture structured nuisance signals, and ridge regression mitigates instability from collinearity between trial regressors. For each subject, minimally preprocessed functional data (motion-corrected, spatially aligned, and high-pass filtered) were modeled using a design matrix constructed from trial onset times, durations, and condition labels. GLMsingle first fits an initial model using a canonical HRF to identify a noise pool of voxels with minimal task-related variance. Time series from these voxels are summarized using principal component analysis, producing nuisance regressors that capture structured noise unrelated to the experimental design. The method then estimates voxel-specific HRFs by selecting, for each voxel, the candidate HRF that maximizes cross-validated prediction accuracy, thereby accounting for spatial variability in hemodynamic timing and shape. Noise regressors derived from the noise pool are optimized iteratively using leave-one-run-out cross-validation, with the number of regressors retained determined by the configuration that maximizes predictive accuracy on held-out data. With the optimal HRFs and noise regressors fixed, GLMsingle then applies ridge regression to estimate a separate *β* weight for each trial, selecting the regularization strength for each voxel via cross-validation to balance noise suppression and signal fidelity. The output of this procedure consists of single-trial *β* coefficient maps alongside diagnostic measures, including voxelwise R², HRF parameters, and regularization strengths. All modeling was performed in each subject’s native space to preserve fine-grained spatial patterns, and the resulting *β* maps were only transformed into standard space for group-level statistical analysis.

#### fMRI pattern similarity analysis

To quantify pattern similarity, we used single-trial *β* maps estimated with the GLMsingle toolbox (fMRI single-trial *β*-estimate)^132^. For each subject and each ROI, we extracted the activity pattern corresponding to every trial from the single-trial *β* maps. Pattern similarity was computed as the Pearson correlation (*r*) between the activity vector from the first presentation of a target and the activity vector from its corresponding test trial, which could be either an identical repetition (“old” item) or a perceptually similar lure (“lure item”). To control for shared activity structure unrelated to the specific studied item, we calculated a baseline pattern similarity value for each trial. This baseline was defined as the mean correlation between the activity vector of the first presentation and the activity vectors of all non-matching test trials from the same run, condition, and stimulus category (objects or scenes). Restricting the baseline to the same run, condition, and category controls for temporal proximity effects, low-level visual similarities, and category-level activation patterns that can inflate pattern similarity estimates. We then subtracted this baseline from the matching-trial correlation, yielding a *similarity score* that reflects the degree of item-specific neural pattern similarity above and beyond general similarity structure. This similarity score was Fisher *z*-transformed and served as the dependent measure in all subsequent multivariate statistical analyses. This approach follows prior work showing that baseline correction is critical to isolate event-specific pattern similarity in fMRI analyses ^120,134^.

#### fMRI searchlight analysis in template space

We conducted multivariate searchlight analyses using CoSMoMVPA ^135^. For each participant, first-level searchlight maps were computed in native space within a subject-specific MTL mask. A spherical searchlight (radius = 4 mm) was iteratively moved throughout the mask, and similarity values were assigned to the central voxel, yielding subject-level searchlight maps. Separate maps were generated for objects and scenes, and within each stimulus category for three conditions of interest for our hypotheses (correct lures, incorrect lures, and target repetitions) and one for an exploratory analysis (all lures combined). This resulted in eight subject-specific searchlight maps (main comparisons of interest: object-lure correct, object-lure incorrect, object-repetition, scene-lure correct, scene-lure incorrect, scene-repetition; exploratory analysis: object-lure all, scene-lure all). The same procedure was carried out for the analyses in which lures were split into similarity bins, comparing each contrast (correct lures vs. repeats, correct vs. incorrect lures) within each lure bin.

Individual subject maps were nonlinearly transformed into a common group template space using ANTs ^127^. Group-level inference was performed with SnPM13 ^136^ using cluster-extent permutation testing (10,000 permutations). Clusters were defined using the default cluster-forming threshold of *p*<.001 (uncorrected), and significance was assessed using family-wise error (FWE) correction based on the permutation distribution. We tested paired contrasts between conditions, focusing on correct lures versus repeats and correct lures versus incorrect lures, separately for objects and scenes. Only clusters significant at *p*<.05, FWE-corrected, are reported based on a one-tailed test motivated by our hypothesis of lower pattern similarity in CA3 and DG.

##### Visualisation of group-level fMRI results

All group-level thresholded fMRI results were visualised with MRIcroGL ^131^. In addition to single modality thresholded images, we also computed overlap contrast maps between the univariate and the multivariate searchlight second-level analyses for the correct lures vs. repeats and the correct vs. incorrect lures comparisons. The overlap map was computed using *fslmaths* by multiplying the binarised, thresholded statistical maps from both analyses.

##### Statistical modelling

All analyses conducted on *β*-values extracted from fMRI statistical maps were carried out in *R*. Where possible, we conducted equivalent statistical tests on uni- and multivariate trial- and contrast-level *β*-estimates to allow for a direct comparison between these two methods. For univariate estimates, we use the terms “response magnitude” or “activation strength” given that all estimates reflect either differences in overall activity between two trial types (contrast *β*) or the level of activity during a given trial type (trial-level *β*) averaged across all voxels. For multivariate estimates, we refer to “pattern similarity” given that we compare the similarity of activity patterns between the first target presentation and either a repeat, correct, or incorrect lure, respectively. All pattern similarity values are thus calculated relative to the first target. Unless otherwise specified, all statistical tests are two-tailed and corrected for multiple comparisons across ROIs using the false discovery rate (FDR) on the level of contrasts and stimulus type.

#### Characterizing response profiles across all ROIs with univariate and multivariate estimates of brain activity

First, we sought to characterize regions in terms of their response profiles to each trial type to uncover potential mechanisms through which they contribute to successful mnemonic discrimination. We reasoned that the strongest evidence we could obtain for pattern separation-like responses in the univariate brain data would be a combination of i) lower activity during a repeated target compared to the first target presentation (‘repetition suppression’), ii) greater activity during correct lures compared to repeats (‘lure-related novelty responses’), iii) no reduced activity in response to correct lures when compared to novel stimuli (resistance to feature overlap), and iii) greater activity during correct than incorrect lures ^31,32,61^. Of note, whole-brain voxel-wise clusters for correct lures > repeats and correct > incorrect lures on their own are insufficient to provide the same level of evidence for potential pattern separation, because these analyses leave open the question whether a region suppressed its activity upon presentation of the lure compared to the first presentation of a target ^19,72^. If such suppression did occur, the purported “pattern separation” region did not fully treat the lure like a novel stimulus and evidence for pattern separation should be considered weaker than in cases where no such suppression took place. This issue can be mitigated by comparing responses across all trial types.

Complementing these univariate results, the strongest evidence for pattern separation would fulfil further criteria based on multivariate analysis showing iv) reduced pattern similarity for correct lures compared to repeated targets which suggests that a region is actively orthogonalizing the neural representations of similar lures compared to the target; and v) reduced pattern similarity for correct lures relative to incorrect lures, indicative of successful pattern separation at the behavioral level. Having defined these uni- and multivariate fMRI proxies, we could determine how many pattern separation criteria each region meets.

Univariate trial-level *β*-estimates were subjected to a linear mixed model with *lmer* from the *lme4* package with fixed effects of ROI and trial type (first object target, repeated object target, correct object lure, incorrect object lure, and the equivalent for scenes) and random effect of participant ^137^. Estimated marginal means (EMMs) were obtained using *emmeans* from the *emmeans* package ^138^. We obtained *t*-statistics and Cohen’s *d* as effect size for each ROI for the pairwise comparisons of firsts vs. repeats, firsts vs. correct lures, correct lures vs. repeats, and correct vs. incorrect lures

For the multivariate approach, we ran a linear mixed model on item-specific pattern similarity which included fixed effects for trial type (repeats, correct lures, incorrect lures), stimulus (objects, scenes), and ROI, and random intercepts for subject and item, thereby controlling for individual differences in mean pattern similarity and systematic variance related to the specific stimulus. We obtained EMMs and performed planned pairwise contrasts for repeats vs. correct lures and correct vs. incorrect lures. For the correct > incorrect lures contrast, a negative value indicates that pattern similarity ERS is lower for lures that were correctly rejected than for those that were falsely recognised, a proxy for successful pattern separation at a behavioral level (e.g., greater neural differentiation between similar items supports correct lure rejection; see ^34,80^). Negative values in the correct lures > repeats contrast indicate that pattern similarity between correct lures and first targets was lower than similarity between first targets and repeats, which may reflect orthogonalisation of the original representation of the lure vis-a-vis the target.

#### Modulating target-lure similarity to estimate region-wise resistance to feature overlap

Lures vary in their degree to which they share features with their corresponding target. Theoretical models of hippocampal pattern separation, particularly from univariate fMRI studies ^11^, posit that regions capable of pattern separation do not track feature similarity in a linear fashion. Rather, if a region successfully orthogonalises lure representations, trial-level activity during first target presentations should resemble activity levels of lures with low to moderate feature overlap. When a region is no longer capable of distinguishing targets and increasingly similar lures, a sharp drop in activity is expected with activity levels to subsequent trials resembling that observed during repeats. We hypothesised that DG specifically (and possibly CA3) maintain similar activity levels during firsts and moderate-to-high lures, while cortical MTL regions may be capable of responding to low similarity lures in a similar fashion as a first target presentation. Similarly, we expected that pattern separation as indexed by neural similarity in CA3 and DG should become more evident as target-lure similarity increases from low to moderate (and possibly high, although this may result in a switch to pattern reinstatement as indicated by increased neural similarity).

As described above, we split lures into similarity bins based on performance measures in an independent cohort. This approach avoids the issue of circularity where bins are selected based on the same sample for which their brain activity is measured. We followed a similar procedure as in prior studies ^34,36,80^. Specifically, we defined several models in line with various potential interpretations of the neural mechanisms behind univariate changes in brain activity across trial types. We fit all models separately for each ROI by stimulus type combination. We identified the best fitting model based on the Akaike Information Criterion. Candidate models included either i) the intercept only, or a ii) linear, iii) quadratic, or iv) cubic effect of similarity as well as models in which we recoded trial types to test whether brain activity was most indicate of v) strong suppression to any kind of feature overlap (1=first target, 0=all other trial types), vi) suppression to moderate feature overlap (2=first, 1=low similarity lure, 0=all other trial types), vii) resistance to low degrees of feature overlap (1=first, 1=low similarity lure, 0=all other trial types), viii) resistance to moderate degrees of feature overlap (1=first, 1=low similarity lure, 1=moderate similarity lure, 0=all other trial types), or xi) resistance to high degrees of feature overlap (1=all except for repeats, 0=repeats). Accordingly, if a region is best fit by a model with a flat line between firsts and high similarity lures, followed by a steep drop in activity when presented with a repeat, this would suggest lure-related novelty responses regardless of the amount of feature overlap. In contrast, a steep drop in activity from firsts to low similarity lures and a subsequent, less pronounced change or stable activity from low similarity lure lures to repeats suggests a strong preference for complete novelty.

Next, we tested our hypothesis that higher levels of feature overlap tend to engage hippocampal pattern separation, while low degrees of feature overlap can be handled by extra-hippocampal regions. We defined a hippocampal and an extra-hippocampal MTL mask and for each mask, extracted the number of above-threshold voxels in the multivariate searchlight maps for each similarity bin for the correct vs. incorrect lure contrast. We then conducted a chi-square test to determine whether the proportion of significant voxels within each anatomical division differed from what would be expected based on the total number of significant voxels in that bin. This proportion was calculated as the number of above-threshold voxels in a region divided by the total number of significant voxels across both hippocampal and extra-hippocampal regions. This approach allowed us to go beyond the binary memory outcomes of the previous model and test whether the graded similarity structure of the stimuli influences reinstatement in a manner predicted by pattern separation theory.

#### Assessing the relationship between brain responses and intra- and inter-individual variability in memory performance

Prior analyses contrasted all trials in a given condition across all participants. Next, we sought to determine whether brain responses track behavior on a trial-by-trial basis and explain inter-individual variability in memory performance. We formulated the following hypotheses: lures with lower pattern similarity are more likely correctly rejected as new; individuals with greater average univariate activity during correctly rejected lures compared to repeats and incorrectly endorsed lures, respectively, tend to also have better mnemonic discrimination; and individuals with lower pattern similarity between first targets and lures compared to first targets and repeats have better memory performance. To keep the number of comparisons low, our planned analysis focused on correlations for all CA3 and DG ROIs. We conducted an exploratory analysis across all ROIs (see Supplementary Materials).

We tested the first hypothesis by running a generalised mixed linear model (*glmer* in R) for each ROI focused on lure trials, predicting whether the lure was a correct rejection or a false alarm. Fixed effects were neural pattern similarity and stimulus type. The model included random intercepts on the stimulus and subject level. To test the second hypothesis on univariate brain activity estimates, we focused on contrast *β*-estimate for the correct lure vs. repeat and correct vs. incorrect lures comparisons. The behavioral outcome was mean corrected hit rate for object and scene discrimination, respectively. We used Pearson correlations between corrected hit rates and contrast *β*-estimates. For our third hypothesis on neural pattern similarity, we correlated subject-level pattern similarity contrast estimates with corrected hit rates. A negative correlation was interpreted as consistent with a pattern separation account, reflecting lower neural similarity for correct lures (relative to repetitions and incorrect lures) in individuals with better lure discrimination performance.

In an exploratory analysis, we identified the best combination of predictors for inter-individual variability in memory performance. Based on the correlations, we chose the top 20 uni- and multivariate brain measures, respectively, that were most strongly correlated (negatively or positively) with corrected hit rates. We set the maximum number of predictors per model to three (given the sample size and limitations on model size) and tested all possible combinations using linear regressions and five fold cross-validation to obtain the mean root squared error for selection of the best fitting model. We used this metric instead of AIC because not all models contained the exact same cases given that for some measures, extreme outliers had to be removed.

#### Uncovering the relationship between univariate contrasts of response magnitude and neural pattern similarity

To examine the correspondence between univariate and multivariate indices of pattern separation, we computed Pearson correlations between subject-specific estimates from both approaches. We focused on the correct lure vs. repeats and correct vs. incorrect lures contrasts. Univariate estimates were subject-level contrast *β* statistics. To obtain subject-level multivariate estimates, we fit the same linear mixed-effects model for each subject individually with pattern similarity scores as dependent variable, with trial type (repeat, correct lure, incorrect lure), stimulus category (object, scene), and ROI as fixed effects, and random intercepts per stimulus. We then extracted EMMs per subject, ROI, and stimulus type to quantify multivariate pattern separation strength. To keep the number of comparisons low, our planned analysis focused on correlations for all CA3 and DG ROIs. We also conducted an exploratory analysis across all ROIs (see Supplementary Materials).

## KEY RESOURCES TABLE

Advanced Normalization Tools (ANTs)

Automated Segmentation of Hippocampal Subfields (ASHS)

FMRIB Software Library (FSL)

ITKsnap

MRIcroGL Presentation

R, RStudio

Statistical Parametric Mapping 12 (SPM12)

GLMsingle

CoSMo Multivariate Pattern Analysis (CoSMoMVPA)

Statistical nonParametric Mapping 13 (SnPM13)

## Acknowledgements

We thank the Leibniz Institute for Neurobiology in Magdeburg for providing access to the 7 Tesla MRI Scanner and Anne Hochkeppler and Regina Schwarzer for support with manual segmentations of the medial temporal lobe subregions. We are grateful to Dharshan Kumaran for stimulating discussions of the manuscript.

This work was supported by the Deutsche Forschungsgemeinschaft (DFG, German Research Foundation) – Project-ID 42589994 (E.D. and D.B.). D.B. has received funding from the European Union’s Horizon 2020 research and innovation programme under the Marie Skłodowska-Curie grant agreement No. 843074. H.M.G. received funding by the BrightFocus Foundation, the DZNE Foundation, the Joachim Herz Foundation and acknowledges the support of St John’s College Cambridge. X.P. received a fellowship from the Deutsche Forschungsgemeinschaft (DFG, German Research Foundation) - Project-ID 550158511 and was further supported by Medical Research Council MR/W01971X/1 (PI: Helen C. Barron).

