## Supplementary Material for "Dynamic shifts between pattern separation and reinstatement in human CA3 and dentate gyrus"

1. *Memory performance*

Variability in mnemonic discrimination performance was driven by correct rejections of lures, rather than hits, consistent with the idea that distinguishing overlapping events rather than novelty detection posed the main memory challenge ($F(1,132) = 171.11$, $p < .001$, $\eta^2 =$ .56;). There was also a moderate effect of category on corrected hit rates with scenes being more difficult ($F(1,66) = 4.16$, $p = .045$, $\eta^2 = .06$), despite categories having been matched for difficulty based independent ratings.

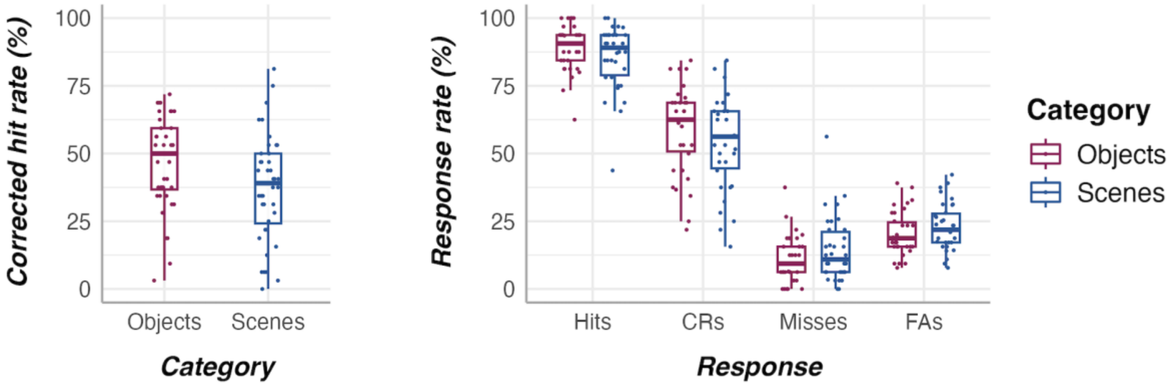


**Supplementary Figure 1. Memory performance in object and scene mnemonic discrimination tasks for corrected hit rates and response types. Left: Asterisk indicates a main effect of category. Right: Asterisks indicate main effects of response type across both categories taken together. Notes. CRs: correct rejections; FAs: false alarms. ****p***<.05.**

**Supplementary Table 1. Results from a repeated-measures ANOVA with within-subjects factor of stimulus type (objects, scenes) and corrected hit rates (hits – false alarms) as outcome.**

Repeated-measures ANOVA: Corrected Hit Rates

| Effect | Degrees of freedom | F-value | p-value | Partial eta squared |
|---|---|---|---|---|
| Sex (male, female) | 1 | 3.951 | 0.051 | 0.054 |
| Age | 1 | 0.994 | 0.322 | 0.014 |
| Stimulus Type (objects, scenes) | 1 | 4.347 | 0.041 | 0.059 |
| Residuals | 64 | NA | NA | NA |

**Supplementary Table 2. Results from a repeated-measures ANOVA with within-subjects factors of stimulus type (objects, scenes) and response type (correct rejections, hits) and response rate as outcome.**

Repeated-measures ANOVA: Response Rates

| Effect | Degrees of freedom | F-value | p-value | Partial eta squared |
|---|---|---|---|---|
| Sex (male, female) | 1 | 2.850 | 0.094 | 0.009 |
| Age | 1 | 0.828 | 0.364 | 0.003 |
| Stimulus Type (objects, scenes) | 1 | 3.534 | 0.062 | 0.011 |
| Response Type (correct rejections, hits) | 1 | 173.281 | 0.000 | 0.558 |
| Interactions of Stimulus and Response Type | 1 | 0.234 | 0.629 | 0.001 |
| Residuals | 130 | NA | NA | NA |

2. *Region of interest analysis for effects of trial type on univariate and multivariate brain responses*

**Supplementary Table 3. Overview of pairwise comparisons of estimated marginal means for a mixed linear model of trial type (first objects, first scenes, correct object lures, correct scene lures, incorrect object lures, incorrect scene lures, repeated objects, repeated scenes) and region of interest for univariate fMRI results (beta estimates for each trial type obtained from first-level general linear models).**

| Region | Stimulus | Comparison | Estimated marginal mean for differences in trial-wise beta estimates | Standard error | Cohen's d | Uncorrected p-value | Corrected p-value |
|---|---|---|---|---|---|---|---|
| AMY (L) | Objects | Lure correct > Repeat | 0.637 | 0.480 | 0.327 | 0.184 | 0.224 |
| AMY (L) | Objects | Lure correct > Lure incorrect | 1.577 | 0.484 | 0.809 | 0.001 | 0.006 |
| AMY (L) | Objects | First > Repeat | 1.516 | 0.480 | 0.778 | 0.002 | 0.003 |
| AMY (L) | Objects | First > Lure correct | 0.879 | 0.480 | 0.451 | 0.067 | 0.183 |
| AMY (L) | Scenes | Lure correct > Repeat | -0.003 | 0.484 | -0.002 | 0.994 | 0.994 |
| AMY (L) | Scenes | Lure correct > Lure incorrect | 0.626 | 0.484 | 0.321 | 0.195 | 0.651 |
| AMY (L) | Scenes | First > Repeat | 1.247 | 0.484 | 0.640 | 0.010 | 0.017 |
| AMY (L) | Scenes | First > Lure correct | 1.251 | 0.480 | 0.642 | 0.009 | 0.046 |
| AMY (R) | Objects | Lure correct > Repeat | 1.114 | 0.488 | 0.571 | 0.022 | 0.067 |
| AMY (R) | Objects | Lure correct > Lure incorrect | 1.499 | 0.480 | 0.769 | 0.002 | 0.008 |
| AMY (R) | Objects | First > Repeat | 1.745 | 0.488 | 0.895 | 0.000 | 0.001 |
| AMY (R) | Objects | First > Lure correct | 0.631 | 0.480 | 0.324 | 0.189 | 0.471 |
| AMY (R) | Scenes | Lure correct > Repeat | 0.192 | 0.484 | 0.098 | 0.692 | 0.768 |
| AMY (R) | Scenes | Lure correct > Lure incorrect | 0.431 | 0.484 | 0.221 | 0.373 | 0.673 |
| AMY (R) | Scenes | First > Repeat | 1.111 | 0.487 | 0.570 | 0.023 | 0.031 |
| AMY (R) | Scenes | First > Lure correct | 0.919 | 0.484 | 0.472 | 0.057 | 0.132 |
| BA35 (L) | Objects | Lure correct > Repeat | 0.806 | 0.495 | 0.414 | 0.103 | 0.155 |
| BA35 (L) | Objects | Lure correct > Lure incorrect | 0.535 | 0.495 | 0.274 | 0.280 | 0.323 |
| BA35 (L) | Objects | First > Repeat | 0.891 | 0.495 | 0.457 | 0.072 | 0.077 |
| BA35 (L) | Objects | First > Lure correct | 0.085 | 0.495 | 0.043 | 0.864 | 0.931 |
| BA35 (L) | Scenes | Lure correct > Repeat | 0.389 | 0.495 | 0.200 | 0.432 | 0.518 |
| BA35 (L) | Scenes | Lure correct > Lure incorrect | 0.435 | 0.495 | 0.223 | 0.380 | 0.673 |
| BA35 (L) | Scenes | First > Repeat | 0.745 | 0.495 | 0.382 | 0.132 | 0.137 |
| BA35 (L) | Scenes | First > Lure correct | 0.356 | 0.495 | 0.183 | 0.472 | 0.675 |
| BA35 (R) | Objects | Lure correct > Repeat | 0.539 | 0.495 | 0.276 | 0.277 | 0.296 |
| BA35 (R) | Objects | Lure correct > Lure incorrect | 0.338 | 0.495 | 0.173 | 0.495 | 0.512 |
| BA35 (R) | Objects | First > Repeat | 0.868 | 0.495 | 0.445 | 0.080 | 0.082 |

| Region | Stimulus | Comparison | Estimated marginal mean for differences in trial-wise beta estimates | Standard error | Cohen's d | Uncorrected p-value | Corrected p-value |
|---|---|---|---|---|---|---|---|
| BA35 (R) | Objects | First > Lure correct | 0.329 | 0.495 | 0.169 | 0.506 | 0.867 |
| BA35 (R) | Scenes | Lure correct > Repeat | 0.868 | 0.495 | 0.445 | 0.080 | 0.217 |
| BA35 (R) | Scenes | Lure correct > Lure incorrect | 0.807 | 0.495 | 0.414 | 0.103 | 0.651 |
| BA35 (R) | Scenes | First > Repeat | 0.748 | 0.495 | 0.384 | 0.131 | 0.137 |
| BA35 (R) | Scenes | First > Lure correct | -0.120 | 0.495 | -0.062 | 0.808 | 0.836 |
| BA36 (L) | Objects | Lure correct > Repeat | 0.654 | 0.495 | 0.335 | 0.187 | 0.224 |
| BA36 (L) | Objects | Lure correct > Lure incorrect | 0.488 | 0.495 | 0.251 | 0.324 | 0.360 |
| BA36 (L) | Objects | First > Repeat | 0.893 | 0.495 | 0.458 | 0.071 | 0.077 |
| BA36 (L) | Objects | First > Lure correct | 0.239 | 0.495 | 0.123 | 0.629 | 0.931 |
| BA36 (L) | Scenes | Lure correct > Repeat | 0.221 | 0.499 | 0.113 | 0.659 | 0.760 |
| BA36 (L) | Scenes | Lure correct > Lure incorrect | 0.135 | 0.495 | 0.069 | 0.785 | 0.812 |
| BA36 (L) | Scenes | First > Repeat | 0.835 | 0.499 | 0.429 | 0.094 | 0.105 |
| BA36 (L) | Scenes | First > Lure correct | 0.615 | 0.495 | 0.315 | 0.214 | 0.402 |
| BA36 (R) | Objects | Lure correct > Repeat | 0.686 | 0.495 | 0.352 | 0.166 | 0.224 |
| BA36 (R) | Objects | Lure correct > Lure incorrect | 0.690 | 0.495 | 0.354 | 0.163 | 0.223 |
| BA36 (R) | Objects | First > Repeat | 0.625 | 0.495 | 0.321 | 0.207 | 0.207 |
| BA36 (R) | Objects | First > Lure correct | -0.061 | 0.495 | -0.031 | 0.902 | 0.931 |
| BA36 (R) | Scenes | Lure correct > Repeat | 0.593 | 0.495 | 0.304 | 0.231 | 0.356 |
| BA36 (R) | Scenes | Lure correct > Lure incorrect | 0.592 | 0.495 | 0.304 | 0.232 | 0.651 |
| BA36 (R) | Scenes | First > Repeat | 0.435 | 0.499 | 0.223 | 0.384 | 0.384 |
| BA36 (R) | Scenes | First > Lure correct | -0.158 | 0.499 | -0.081 | 0.751 | 0.805 |
| ALERC (L) | Objects | Lure correct > Repeat | 0.911 | 0.480 | 0.467 | 0.058 | 0.102 |
| ALERC (L) | Objects | Lure correct > Lure incorrect | 0.916 | 0.480 | 0.470 | 0.056 | 0.130 |
| ALERC (L) | Objects | First > Repeat | 1.305 | 0.480 | 0.670 | 0.007 | 0.010 |
| ALERC (L) | Objects | First > Lure correct | 0.394 | 0.480 | 0.202 | 0.411 | 0.822 |
| ALERC (L) | Scenes | Lure correct > Repeat | 0.567 | 0.480 | 0.291 | 0.238 | 0.356 |
| ALERC (L) | Scenes | Lure correct > Lure incorrect | 0.831 | 0.480 | 0.427 | 0.083 | 0.651 |
| ALERC (L) | Scenes | First > Repeat | 0.952 | 0.480 | 0.489 | 0.047 | 0.059 |
| ALERC (L) | Scenes | First > Lure correct | 0.386 | 0.480 | 0.198 | 0.422 | 0.666 |
| ALERC (R) | Objects | Lure correct > Repeat | 1.035 | 0.480 | 0.531 | 0.031 | 0.078 |
| ALERC (R) | Objects | Lure correct > Lure incorrect | 0.995 | 0.480 | 0.510 | 0.038 | 0.115 |
| ALERC (R) | Objects | First > Repeat | 1.169 | 0.480 | 0.600 | 0.015 | 0.021 |

| Region | Stimulus | Comparison | Estimated marginal mean for differences in trial-wise beta estimates | Standard error | Cohen's d | Uncorrected p-value | Corrected p-value |
|---|---|---|---|---|---|---|---|
| ALERC (R) | Objects | First > Lure correct | 0.134 | 0.480 | 0.069 | 0.780 | 0.931 |
| ALERC (R) | Scenes | Lure correct > Repeat | 1.005 | 0.480 | 0.516 | 0.036 | 0.181 |
| ALERC (R) | Scenes | Lure correct > Lure incorrect | 1.224 | 0.480 | 0.628 | 0.011 | 0.323 |
| ALERC (R) | Scenes | First > Repeat | 1.158 | 0.480 | 0.594 | 0.016 | 0.024 |
| ALERC (R) | Scenes | First > Lure correct | 0.153 | 0.480 | 0.078 | 0.750 | 0.805 |
| PMERC (L) | Objects | Lure correct > Repeat | 0.921 | 0.480 | 0.472 | 0.055 | 0.102 |
| PMERC (L) | Objects | Lure correct > Lure incorrect | 0.189 | 0.480 | 0.097 | 0.693 | 0.693 |
| PMERC (L) | Objects | First > Repeat | 1.344 | 0.484 | 0.690 | 0.005 | 0.009 |
| PMERC (L) | Objects | First > Lure correct | 0.423 | 0.484 | 0.217 | 0.381 | 0.817 |
| PMERC (L) | Scenes | Lure correct > Repeat | 0.688 | 0.480 | 0.353 | 0.152 | 0.303 |
| PMERC (L) | Scenes | Lure correct > Lure incorrect | 0.561 | 0.480 | 0.288 | 0.243 | 0.651 |
| PMERC (L) | Scenes | First > Repeat | 0.899 | 0.484 | 0.461 | 0.063 | 0.073 |
| PMERC (L) | Scenes | First > Lure correct | 0.211 | 0.484 | 0.108 | 0.662 | 0.794 |
| PMERC (R) | Objects | Lure correct > Repeat | 1.015 | 0.480 | 0.521 | 0.034 | 0.079 |
| PMERC (R) | Objects | Lure correct > Lure incorrect | 0.951 | 0.480 | 0.488 | 0.048 | 0.119 |
| PMERC (R) | Objects | First > Repeat | 1.057 | 0.480 | 0.542 | 0.028 | 0.033 |
| PMERC (R) | Objects | First > Lure correct | 0.042 | 0.480 | 0.021 | 0.931 | 0.931 |
| PMERC (R) | Scenes | Lure correct > Repeat | 0.926 | 0.480 | 0.475 | 0.054 | 0.202 |
| PMERC (R) | Scenes | Lure correct > Lure incorrect | 0.604 | 0.480 | 0.310 | 0.208 | 0.651 |
| PMERC (R) | Scenes | First > Repeat | 1.167 | 0.480 | 0.599 | 0.015 | 0.024 |
| PMERC (R) | Scenes | First > Lure correct | 0.242 | 0.480 | 0.124 | 0.615 | 0.768 |
| PHC (L) | Objects | Lure correct > Repeat | 0.914 | 0.499 | 0.469 | 0.067 | 0.112 |
| PHC (L) | Objects | Lure correct > Lure incorrect | 0.615 | 0.499 | 0.316 | 0.218 | 0.261 |
| PHC (L) | Objects | First > Repeat | 2.334 | 0.495 | 1.197 | 0.000 | 0.000 |
| PHC (L) | Objects | First > Lure correct | 1.420 | 0.499 | 0.728 | 0.004 | 0.017 |
| PHC (L) | Scenes | Lure correct > Repeat | 0.729 | 0.495 | 0.374 | 0.141 | 0.303 |
| PHC (L) | Scenes | Lure correct > Lure incorrect | 0.215 | 0.495 | 0.110 | 0.664 | 0.797 |
| PHC (L) | Scenes | First > Repeat | 1.708 | 0.495 | 0.876 | 0.001 | 0.002 |
| PHC (L) | Scenes | First > Lure correct | 0.979 | 0.495 | 0.502 | 0.048 | 0.131 |
| PHC (R) | Objects | Lure correct > Repeat | 0.618 | 0.503 | 0.317 | 0.220 | 0.244 |
| PHC (R) | Objects | Lure correct > Lure incorrect | -0.477 | 0.499 | -0.244 | 0.340 | 0.364 |
| PHC (R) | Objects | First > Repeat | 2.427 | 0.499 | 1.245 | 0.000 | 0.000 |

| Region | Stimulus | Comparison | Estimated marginal mean for differences in trial-wise beta estimates | Standard error | Cohen's d | Uncorrected p-value | Corrected p-value |
|---|---|---|---|---|---|---|---|
| PHC (R) | Objects | First > Lure correct | 1.809 | 0.499 | 0.928 | 0.000 | 0.002 |
| PHC (R) | Scenes | Lure correct > Repeat | 1.183 | 0.495 | 0.607 | 0.017 | 0.127 |
| PHC (R) | Scenes | Lure correct > Lure incorrect | 0.727 | 0.495 | 0.373 | 0.142 | 0.651 |
| PHC (R) | Scenes | First > Repeat | 1.640 | 0.495 | 0.841 | 0.001 | 0.003 |
| PHC (R) | Scenes | First > Lure correct | 0.457 | 0.495 | 0.234 | 0.356 | 0.594 |
| aSUB (L) | Objects | Lure correct > Repeat | 0.685 | 0.503 | 0.351 | 0.174 | 0.224 |
| aSUB (L) | Objects | Lure correct > Lure incorrect | 0.816 | 0.503 | 0.419 | 0.105 | 0.166 |
| aSUB (L) | Objects | First > Repeat | 1.744 | 0.503 | 0.895 | 0.001 | 0.001 |
| aSUB (L) | Objects | First > Lure correct | 1.060 | 0.503 | 0.544 | 0.035 | 0.106 |
| aSUB (L) | Scenes | Lure correct > Repeat | 0.625 | 0.503 | 0.321 | 0.214 | 0.356 |
| aSUB (L) | Scenes | Lure correct > Lure incorrect | 0.430 | 0.503 | 0.220 | 0.393 | 0.673 |
| aSUB (L) | Scenes | First > Repeat | 1.554 | 0.508 | 0.797 | 0.002 | 0.005 |
| aSUB (L) | Scenes | First > Lure correct | 0.929 | 0.508 | 0.477 | 0.067 | 0.144 |
| aSUB (R) | Objects | Lure correct > Repeat | 0.985 | 0.503 | 0.505 | 0.050 | 0.102 |
| aSUB (R) | Objects | Lure correct > Lure incorrect | 1.195 | 0.503 | 0.613 | 0.018 | 0.059 |
| aSUB (R) | Objects | First > Repeat | 2.278 | 0.508 | 1.169 | 0.000 | 0.000 |
| aSUB (R) | Objects | First > Lure correct | 1.293 | 0.508 | 0.663 | 0.011 | 0.036 |
| aSUB (R) | Scenes | Lure correct > Repeat | 0.932 | 0.503 | 0.478 | 0.064 | 0.213 |
| aSUB (R) | Scenes | Lure correct > Lure incorrect | 0.779 | 0.503 | 0.400 | 0.122 | 0.651 |
| aSUB (R) | Scenes | First > Repeat | 1.901 | 0.503 | 0.975 | 0.000 | 0.001 |
| aSUB (R) | Scenes | First > Lure correct | 0.969 | 0.503 | 0.497 | 0.054 | 0.132 |
| aCA1 (L) | Objects | Lure correct > Repeat | 0.487 | 0.503 | 0.250 | 0.334 | 0.334 |
| aCA1 (L) | Objects | Lure correct > Lure incorrect | 0.650 | 0.503 | 0.333 | 0.197 | 0.246 |
| aCA1 (L) | Objects | First > Repeat | 1.951 | 0.503 | 1.001 | 0.000 | 0.000 |
| aCA1 (L) | Objects | First > Lure correct | 1.464 | 0.503 | 0.751 | 0.004 | 0.016 |
| aCA1 (L) | Scenes | Lure correct > Repeat | 0.575 | 0.503 | 0.295 | 0.253 | 0.361 |
| aCA1 (L) | Scenes | Lure correct > Lure incorrect | 0.382 | 0.503 | 0.196 | 0.447 | 0.674 |
| aCA1 (L) | Scenes | First > Repeat | 1.838 | 0.503 | 0.943 | 0.000 | 0.001 |
| aCA1 (L) | Scenes | First > Lure correct | 1.262 | 0.503 | 0.648 | 0.012 | 0.052 |
| aCA1 (R) | Objects | Lure correct > Repeat | 0.720 | 0.503 | 0.369 | 0.153 | 0.218 |
| aCA1 (R) | Objects | Lure correct > Lure incorrect | 0.800 | 0.503 | 0.410 | 0.112 | 0.168 |
| aCA1 (R) | Objects | First > Repeat | 2.381 | 0.503 | 1.222 | 0.000 | 0.000 |

| Region | Stimulus | Comparison | Estimated marginal mean for differences in trial-wise beta estimates | Standard error | Cohen's d | Uncorrected p-value | Corrected p-value |
|---|---|---|---|---|---|---|---|
| aCA1 (R) | Objects | First > Lure correct | 1.662 | 0.503 | 0.852 | 0.001 | 0.005 |
| aCA1 (R) | Scenes | Lure correct > Repeat | 0.644 | 0.508 | 0.330 | 0.205 | 0.356 |
| aCA1 (R) | Scenes | Lure correct > Lure incorrect | 0.754 | 0.508 | 0.387 | 0.137 | 0.651 |
| aCA1 (R) | Scenes | First > Repeat | 2.161 | 0.503 | 1.109 | 0.000 | 0.000 |
| aCA1 (R) | Scenes | First > Lure correct | 1.518 | 0.508 | 0.779 | 0.003 | 0.017 |
| aCA3 (L) | Objects | Lure correct > Repeat | 0.532 | 0.503 | 0.273 | 0.291 | 0.301 |
| aCA3 (L) | Objects | Lure correct > Lure incorrect | 0.675 | 0.503 | 0.346 | 0.180 | 0.235 |
| aCA3 (L) | Objects | First > Repeat | 3.610 | 0.503 | 1.852 | 0.000 | 0.000 |
| aCA3 (L) | Objects | First > Lure correct | 3.078 | 0.503 | 1.579 | 0.000 | 0.000 |
| aCA3 (L) | Scenes | Lure correct > Repeat | -0.059 | 0.503 | -0.030 | 0.906 | 0.955 |
| aCA3 (L) | Scenes | Lure correct > Lure incorrect | -0.152 | 0.503 | -0.078 | 0.762 | 0.812 |
| aCA3 (L) | Scenes | First > Repeat | 2.784 | 0.503 | 1.428 | 0.000 | 0.000 |
| aCA3 (L) | Scenes | First > Lure correct | 2.843 | 0.503 | 1.459 | 0.000 | 0.000 |
| aCA3 (R) | Objects | Lure correct > Repeat | 1.215 | 0.503 | 0.623 | 0.016 | 0.053 |
| aCA3 (R) | Objects | Lure correct > Lure incorrect | 2.493 | 0.503 | 1.279 | 0.000 | 0.000 |
| aCA3 (R) | Objects | First > Repeat | 4.305 | 0.503 | 2.208 | 0.000 | 0.000 |
| aCA3 (R) | Objects | First > Lure correct | 3.090 | 0.503 | 1.585 | 0.000 | 0.000 |
| aCA3 (R) | Scenes | Lure correct > Repeat | 0.048 | 0.503 | 0.025 | 0.924 | 0.955 |
| aCA3 (R) | Scenes | Lure correct > Lure incorrect | 0.236 | 0.503 | 0.121 | 0.640 | 0.797 |
| aCA3 (R) | Scenes | First > Repeat | 3.613 | 0.503 | 1.854 | 0.000 | 0.000 |
| aCA3 (R) | Scenes | First > Lure correct | 3.565 | 0.503 | 1.829 | 0.000 | 0.000 |
| aDG (L) | Objects | Lure correct > Repeat | 0.624 | 0.503 | 0.320 | 0.215 | 0.244 |
| aDG (L) | Objects | Lure correct > Lure incorrect | 0.823 | 0.503 | 0.422 | 0.102 | 0.166 |
| aDG (L) | Objects | First > Repeat | 3.004 | 0.503 | 1.541 | 0.000 | 0.000 |
| aDG (L) | Objects | First > Lure correct | 2.381 | 0.503 | 1.221 | 0.000 | 0.000 |
| aDG (L) | Scenes | Lure correct > Repeat | 0.781 | 0.503 | 0.400 | 0.121 | 0.279 |
| aDG (L) | Scenes | Lure correct > Lure incorrect | 0.546 | 0.503 | 0.280 | 0.278 | 0.651 |
| aDG (L) | Scenes | First > Repeat | 2.586 | 0.503 | 1.327 | 0.000 | 0.000 |
| aDG (L) | Scenes | First > Lure correct | 1.805 | 0.503 | 0.926 | 0.000 | 0.003 |
| aDG (R) | Objects | Lure correct > Repeat | 1.117 | 0.503 | 0.573 | 0.026 | 0.072 |
| aDG (R) | Objects | Lure correct > Lure incorrect | 2.114 | 0.503 | 1.085 | 0.000 | 0.000 |
| aDG (R) | Objects | First > Repeat | 3.668 | 0.503 | 1.882 | 0.000 | 0.000 |

| Region | Stimulus | Comparison | Estimated marginal mean for differences in trial-wise beta estimates | Standard error | Cohen's d | Uncorrected p-value | Corrected p-value |
|---|---|---|---|---|---|---|---|
| aDG (R) | Objects | First > Lure correct | 2.551 | 0.503 | 1.309 | 0.000 | 0.000 |
| aDG (R) | Scenes | Lure correct > Repeat | 0.805 | 0.508 | 0.413 | 0.113 | 0.279 |
| aDG (R) | Scenes | Lure correct > Lure incorrect | 0.365 | 0.508 | 0.187 | 0.472 | 0.674 |
| aDG (R) | Scenes | First > Repeat | 3.186 | 0.503 | 1.634 | 0.000 | 0.000 |
| aDG (R) | Scenes | First > Lure correct | 2.380 | 0.508 | 1.221 | 0.000 | 0.000 |
| pSUB (L) | Objects | Lure correct > Repeat | 1.241 | 0.508 | 0.637 | 0.015 | 0.053 |
| pSUB (L) | Objects | Lure correct > Lure incorrect | 1.367 | 0.508 | 0.701 | 0.007 | 0.027 |
| pSUB (L) | Objects | First > Repeat | 1.053 | 0.508 | 0.540 | 0.038 | 0.044 |
| pSUB (L) | Objects | First > Lure correct | -0.188 | 0.503 | -0.097 | 0.708 | 0.931 |
| pSUB (L) | Scenes | Lure correct > Repeat | 0.650 | 0.508 | 0.333 | 0.200 | 0.356 |
| pSUB (L) | Scenes | Lure correct > Lure incorrect | -0.137 | 0.503 | -0.070 | 0.785 | 0.812 |
| pSUB (L) | Scenes | First > Repeat | 0.990 | 0.508 | 0.508 | 0.051 | 0.062 |
| pSUB (L) | Scenes | First > Lure correct | 0.340 | 0.503 | 0.174 | 0.500 | 0.682 |
| pSUB (R) | Objects | Lure correct > Repeat | 1.433 | 0.508 | 0.735 | 0.005 | 0.029 |
| pSUB (R) | Objects | Lure correct > Lure incorrect | 0.948 | 0.508 | 0.486 | 0.062 | 0.132 |
| pSUB (R) | Objects | First > Repeat | 1.522 | 0.503 | 0.781 | 0.003 | 0.004 |
| pSUB (R) | Objects | First > Lure correct | 0.089 | 0.508 | 0.046 | 0.861 | 0.931 |
| pSUB (R) | Scenes | Lure correct > Repeat | 1.316 | 0.503 | 0.675 | 0.009 | 0.104 |
| pSUB (R) | Scenes | Lure correct > Lure incorrect | 0.420 | 0.503 | 0.215 | 0.404 | 0.673 |
| pSUB (R) | Scenes | First > Repeat | 1.328 | 0.503 | 0.681 | 0.008 | 0.016 |
| pSUB (R) | Scenes | First > Lure correct | 0.012 | 0.503 | 0.006 | 0.982 | 0.982 |
| pCA1 (L) | Objects | Lure correct > Repeat | 0.975 | 0.503 | 0.500 | 0.053 | 0.102 |
| pCA1 (L) | Objects | Lure correct > Lure incorrect | 0.902 | 0.503 | 0.463 | 0.073 | 0.134 |
| pCA1 (L) | Objects | First > Repeat | 1.164 | 0.503 | 0.597 | 0.021 | 0.027 |
| pCA1 (L) | Objects | First > Lure correct | 0.189 | 0.503 | 0.097 | 0.707 | 0.931 |
| pCA1 (L) | Scenes | Lure correct > Repeat | 0.542 | 0.503 | 0.278 | 0.282 | 0.375 |
| pCA1 (L) | Scenes | Lure correct > Lure incorrect | 0.140 | 0.508 | 0.072 | 0.783 | 0.812 |
| pCA1 (L) | Scenes | First > Repeat | 1.186 | 0.503 | 0.608 | 0.018 | 0.026 |
| pCA1 (L) | Scenes | First > Lure correct | 0.644 | 0.503 | 0.330 | 0.201 | 0.401 |
| pCA1 (R) | Objects | Lure correct > Repeat | 1.282 | 0.503 | 0.658 | 0.011 | 0.053 |
| pCA1 (R) | Objects | Lure correct > Lure incorrect | 1.017 | 0.503 | 0.522 | 0.043 | 0.118 |
| pCA1 (R) | Objects | First > Repeat | 1.436 | 0.503 | 0.737 | 0.004 | 0.007 |

| Region | Stimulus | Comparison | Estimated marginal mean for differences in trial-wise beta estimates | Standard error | Cohen's d | Uncorrected p-value | Corrected p-value |
|---|---|---|---|---|---|---|---|
| pCA1 (R) | Objects | First > Lure correct | 0.154 | 0.503 | 0.079 | 0.759 | 0.931 |
| pCA1 (R) | Scenes | Lure correct > Repeat | 0.997 | 0.503 | 0.511 | 0.048 | 0.202 |
| pCA1 (R) | Scenes | Lure correct > Lure incorrect | 0.541 | 0.503 | 0.278 | 0.282 | 0.651 |
| pCA1 (R) | Scenes | First > Repeat | 1.371 | 0.503 | 0.703 | 0.006 | 0.013 |
| pCA1 (R) | Scenes | First > Lure correct | 0.374 | 0.503 | 0.192 | 0.457 | 0.675 |
| pCA3 (L) | Objects | Lure correct > Repeat | 1.465 | 0.503 | 0.751 | 0.004 | 0.027 |
| pCA3 (L) | Objects | Lure correct > Lure incorrect | 0.740 | 0.508 | 0.380 | 0.145 | 0.207 |
| pCA3 (L) | Objects | First > Repeat | 2.017 | 0.503 | 1.035 | 0.000 | 0.000 |
| pCA3 (L) | Objects | First > Lure correct | 0.552 | 0.503 | 0.283 | 0.273 | 0.630 |
| pCA3 (L) | Scenes | Lure correct > Repeat | 0.403 | 0.503 | 0.207 | 0.423 | 0.518 |
| pCA3 (L) | Scenes | Lure correct > Lure incorrect | -0.379 | 0.503 | -0.194 | 0.452 | 0.674 |
| pCA3 (L) | Scenes | First > Repeat | 1.592 | 0.503 | 0.817 | 0.002 | 0.004 |
| pCA3 (L) | Scenes | First > Lure correct | 1.189 | 0.503 | 0.610 | 0.018 | 0.068 |
| pCA3 (R) | Objects | Lure correct > Repeat | 2.182 | 0.503 | 1.119 | 0.000 | 0.000 |
| pCA3 (R) | Objects | Lure correct > Lure incorrect | 2.265 | 0.503 | 1.162 | 0.000 | 0.000 |
| pCA3 (R) | Objects | First > Repeat | 2.505 | 0.503 | 1.285 | 0.000 | 0.000 |
| pCA3 (R) | Objects | First > Lure correct | 0.324 | 0.503 | 0.166 | 0.520 | 0.867 |
| pCA3 (R) | Scenes | Lure correct > Repeat | 0.897 | 0.503 | 0.460 | 0.075 | 0.217 |
| pCA3 (R) | Scenes | Lure correct > Lure incorrect | 0.225 | 0.503 | 0.115 | 0.655 | 0.797 |
| pCA3 (R) | Scenes | First > Repeat | 1.996 | 0.503 | 1.024 | 0.000 | 0.000 |
| pCA3 (R) | Scenes | First > Lure correct | 1.099 | 0.503 | 0.564 | 0.029 | 0.088 |
| pDG (L) | Objects | Lure correct > Repeat | 1.772 | 0.503 | 0.909 | 0.000 | 0.004 |
| pDG (L) | Objects | Lure correct > Lure incorrect | 1.804 | 0.503 | 0.925 | 0.000 | 0.002 |
| pDG (L) | Objects | First > Repeat | 2.007 | 0.508 | 1.030 | 0.000 | 0.000 |
| pDG (L) | Objects | First > Lure correct | 0.235 | 0.508 | 0.121 | 0.644 | 0.931 |
| pDG (L) | Scenes | Lure correct > Repeat | 0.536 | 0.503 | 0.275 | 0.287 | 0.375 |
| pDG (L) | Scenes | Lure correct > Lure incorrect | 0.075 | 0.503 | 0.038 | 0.881 | 0.881 |
| pDG (L) | Scenes | First > Repeat | 1.632 | 0.503 | 0.837 | 0.001 | 0.003 |
| pDG (L) | Scenes | First > Lure correct | 1.096 | 0.503 | 0.562 | 0.029 | 0.088 |
| pDG (R) | Objects | Lure correct > Repeat | 2.410 | 0.503 | 1.236 | 0.000 | 0.000 |
| pDG (R) | Objects | Lure correct > Lure incorrect | 2.318 | 0.503 | 1.189 | 0.000 | 0.000 |
| pDG (R) | Objects | First > Repeat | 2.340 | 0.503 | 1.200 | 0.000 | 0.000 |

| Region | Stimulus | Comparison | Estimated marginal mean for differences in trial-wise beta estimates | Standard error | Cohen's d | Uncorrected p-value | Corrected p-value |
|---|---|---|---|---|---|---|---|
| pDG (R) | Objects | First > Lure correct | -0.070 | 0.503 | -0.036 | 0.890 | 0.931 |
| pDG (R) | Scenes | Lure correct > Repeat | 1.664 | 0.508 | 0.854 | 0.001 | 0.031 |
| pDG (R) | Scenes | Lure correct > Lure incorrect | 0.522 | 0.508 | 0.268 | 0.304 | 0.652 |
| pDG (R) | Scenes | First > Repeat | 2.189 | 0.503 | 1.123 | 0.000 | 0.000 |
| pDG (R) | Scenes | First > Lure correct | 0.524 | 0.508 | 0.269 | 0.302 | 0.533 |
| TAIL (L) | Objects | Lure correct > Repeat | 1.257 | 0.508 | 0.645 | 0.013 | 0.053 |
| TAIL (L) | Objects | Lure correct > Lure incorrect | 0.887 | 0.499 | 0.455 | 0.076 | 0.134 |
| TAIL (L) | Objects | First > Repeat | 1.160 | 0.504 | 0.595 | 0.021 | 0.027 |
| TAIL (L) | Objects | First > Lure correct | -0.097 | 0.499 | -0.050 | 0.846 | 0.931 |
| TAIL (L) | Scenes | Lure correct > Repeat | 1.103 | 0.495 | 0.566 | 0.026 | 0.156 |
| TAIL (L) | Scenes | Lure correct > Lure incorrect | 0.311 | 0.495 | 0.160 | 0.530 | 0.722 |
| TAIL (L) | Scenes | First > Repeat | 1.283 | 0.495 | 0.658 | 0.010 | 0.017 |
| TAIL (L) | Scenes | First > Lure correct | 0.181 | 0.495 | 0.093 | 0.715 | 0.805 |
| TAIL (R) | Objects | Lure correct > Repeat | 0.879 | 0.499 | 0.451 | 0.078 | 0.124 |
| TAIL (R) | Objects | Lure correct > Lure incorrect | 0.917 | 0.499 | 0.470 | 0.066 | 0.133 |
| TAIL (R) | Objects | First > Repeat | 1.201 | 0.495 | 0.616 | 0.015 | 0.021 |
| TAIL (R) | Objects | First > Lure correct | 0.321 | 0.499 | 0.165 | 0.520 | 0.867 |
| TAIL (R) | Scenes | Lure correct > Repeat | 1.268 | 0.495 | 0.651 | 0.010 | 0.104 |
| TAIL (R) | Scenes | Lure correct > Lure incorrect | 0.555 | 0.495 | 0.285 | 0.262 | 0.651 |
| TAIL (R) | Scenes | First > Repeat | 0.986 | 0.495 | 0.506 | 0.046 | 0.059 |
| TAIL (R) | Scenes | First > Lure correct | -0.282 | 0.495 | -0.145 | 0.569 | 0.742 |

**Notes. '*a*': anterior hippocampus; '*p*', posterior hippocampus. ALERC: anterolateral entorhinal cortex; AMY: amygdala; B: bilateral; BA: Brodmann area; CA: cornu ammonis; DG: dentate gyrus; L: left; PHC: parahippocampal cortex; PMERC: posterior-medial ERC; R: right; SUB: subiculum.**

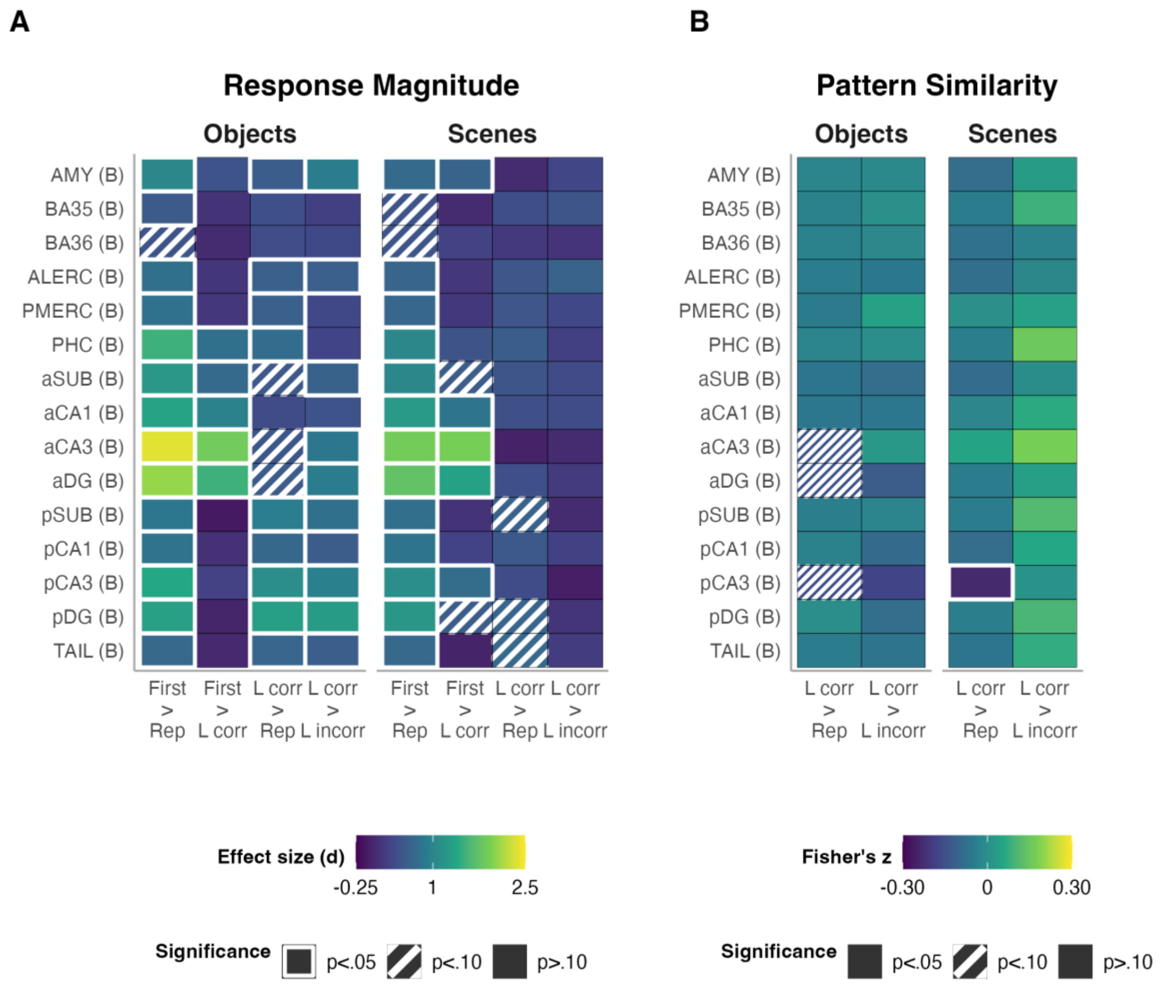


**Supplementary Figure 2. Characterizing response patterns across trial types provides insights into the nature of information processing in medial temporal lobe regions. All regions of interest were collapsed across hemispheres. All p-values were corrected for multiple comparisons on the contrast and stimulus level. A. Analyses were conducted on trial-wise model *beta* estimates from each participant's first-level general linear model, extracted from and averaged across all voxels within the anatomical mask for a region of interest. Coloured tiles show statistically significant pairwise comparisons between individual trial types across all regions of interest and split by stimulus type in a linear mixed model including a region by trial type interaction and participant as random effect. Comparisons were derived from estimated marginal means and were corrected for multiple comparisons at the contrast and stimulus level. Effect sizes are Cohen's *d*. Decreased activity during repeated targets and correct lures relative to the first target suggests sensitivity to feature overlap and a preference for novelty. Similar activity between first targets and correct lures combined with greater activity during correct lures compared to repeats and greater activity during correct vs. incorrect lures is more in line the interpretation of pattern separation according to prior fMRI studies using univariate approaches (Bakker et al., 2008; Berron et al., 2018; Reagh et al., 2018). B. Multivariate analysis on neural pattern similarity. Analyses were conducted on trial-wise beta estimates from each participant's first-level general linear model, extracted from all voxels within the anatomical mask of each region of interest. Effects were corrected for multiple comparisons at the contrast and stimulus level. Decreased neural pattern similarity during correct lure trials relative to repeated targets or incorrect lures is in line with theoretical accounts of pattern separation. Pearson correlation coefficients were Fisher *z*-transformed prior to analysis to normalize their distribution and stabilize variance for statistical testing. Notes. The prefix '*a*' denotes anterior hippocampus; '*p*', posterior hippocampus. ALERC: anterolateral entorhinal cortex; AMY: amygdala; B: bilateral; BA: Brodmann area; CA: cornu ammonis; DG: dentate gyrus; L corr: correct rejection of the lure; L incorr: false alarm to lure; PHC: parahippocampal cortex; PMERC: posterior-medial entorhinal cortex; Rep: repetition of the target item; SUB: subiculum.**

**Supplementary Table 4. Overview of pairwise comparisons of estimated marginal means for a mixed linear model of trial type (repeat, correct lure, incorrect lure by stimulus domain) and region of interest for multivariate fMRI results as measured with neural pattern separation (Fisher's *z*). Pattern similarity always refers to the similarity between the baseline (first target presentations) and a given trial type. The correct lures < repeats contrast thus refers to lower pattern similarity between correct lures and firsts compared to pattern similarity between first and repeated targets.**

| Region | Stimulus | Comparison | Estimated marginal mean for differences in Fisher's z-transformed r | Standard error | Cohen's d | Uncorrected p-value | Corrected p-value |
|---|---|---|---|---|---|---|---|
| AMY (L) | Objects | Lure correct > Repeat | -0.005 | 0.007 | -0.037 | 0.493 | 0.812 |
| AMY (L) | Objects | Lure correct > Lure incorrect | -0.001 | 0.008 | -0.011 | 0.855 | 0.997 |
| AMY (L) | Scenes | Lure correct > Repeat | -0.007 | 0.007 | -0.057 | 0.300 | 0.475 |
| AMY (L) | Scenes | Lure correct > Lure incorrect | 0.005 | 0.008 | 0.038 | 0.536 | 0.874 |
| AMY (R) | Objects | Lure correct > Repeat | -0.004 | 0.007 | -0.033 | 0.550 | 0.824 |
| AMY (R) | Objects | Lure correct > Lure incorrect | -0.002 | 0.008 | -0.014 | 0.829 | 0.997 |
| AMY (R) | Scenes | Lure correct > Repeat | -0.013 | 0.007 | -0.101 | 0.068 | 0.292 |
| AMY (R) | Scenes | Lure correct > Lure incorrect | 0.001 | 0.008 | 0.009 | 0.887 | 0.959 |
| BA35 (L) | Objects | Lure correct > Repeat | -0.001 | 0.007 | -0.009 | 0.875 | 0.938 |
| BA35 (L) | Objects | Lure correct > Lure incorrect | 0.001 | 0.008 | 0.007 | 0.915 | 0.997 |
| BA35 (L) | Scenes | Lure correct > Repeat | -0.006 | 0.007 | -0.044 | 0.426 | 0.532 |
| BA35 (L) | Scenes | Lure correct > Lure incorrect | 0.009 | 0.008 | 0.073 | 0.242 | 0.759 |
| BA35 (R) | Objects | Lure correct > Repeat | -0.003 | 0.007 | -0.020 | 0.718 | 0.840 |
| BA35 (R) | Objects | Lure correct > Lure incorrect | 0.000 | 0.008 | 0.000 | 0.997 | 0.997 |
| BA35 (R) | Scenes | Lure correct > Repeat | -0.007 | 0.007 | -0.056 | 0.323 | 0.475 |
| BA35 (R) | Scenes | Lure correct > Lure incorrect | 0.006 | 0.008 | 0.045 | 0.470 | 0.874 |
| BA36 (L) | Objects | Lure correct > Repeat | -0.007 | 0.007 | -0.054 | 0.321 | 0.812 |
| BA36 (L) | Objects | Lure correct > Lure incorrect | -0.005 | 0.008 | -0.043 | 0.497 | 0.884 |
| BA36 (L) | Scenes | Lure correct > Repeat | -0.014 | 0.007 | -0.113 | 0.044 | 0.218 |
| BA36 (L) | Scenes | Lure correct > Lure incorrect | -0.009 | 0.008 | -0.072 | 0.253 | 0.759 |
| BA36 (R) | Objects | Lure correct > Repeat | -0.003 | 0.007 | -0.027 | 0.616 | 0.830 |
| BA36 (R) | Objects | Lure correct > Lure incorrect | -0.001 | 0.008 | -0.004 | 0.950 | 0.997 |
| BA36 (R) | Scenes | Lure correct > Repeat | -0.008 | 0.007 | -0.062 | 0.268 | 0.473 |
| BA36 (R) | Scenes | Lure correct > Lure incorrect | -0.001 | 0.008 | -0.010 | 0.873 | 0.959 |
| ALERC (L) | Objects | Lure correct > Repeat | -0.008 | 0.007 | -0.064 | 0.241 | 0.804 |
| ALERC (L) | Objects | Lure correct > Lure incorrect | -0.005 | 0.008 | -0.037 | 0.560 | 0.884 |
| ALERC (L) | Scenes | Lure correct > Repeat | -0.012 | 0.007 | -0.093 | 0.097 | 0.292 |

| Region | Stimulus | Comparison | Estimated marginal mean for differences in Fisher's z-transformed r | Standard error | Cohen's d | Uncorrected p-value | Corrected p-value |
|---|---|---|---|---|---|---|---|
| ALERC (L) | Scenes | Lure correct > Lure incorrect | -0.004 | 0.008 | -0.029 | 0.648 | 0.877 |
| ALERC (R) | Objects | Lure correct > Repeat | -0.001 | 0.007 | -0.005 | 0.922 | 0.954 |
| ALERC (R) | Objects | Lure correct > Lure incorrect | -0.006 | 0.008 | -0.048 | 0.449 | 0.884 |
| ALERC (R) | Scenes | Lure correct > Repeat | -0.008 | 0.007 | -0.066 | 0.240 | 0.461 |
| ALERC (R) | Scenes | Lure correct > Lure incorrect | 0.000 | 0.008 | -0.003 | 0.961 | 0.961 |
| PMERC (L) | Objects | Lure correct > Repeat | -0.002 | 0.007 | -0.016 | 0.768 | 0.853 |
| PMERC (L) | Objects | Lure correct > Lure incorrect | 0.004 | 0.008 | 0.029 | 0.640 | 0.960 |
| PMERC (L) | Scenes | Lure correct > Repeat | -0.009 | 0.007 | -0.073 | 0.190 | 0.408 |
| PMERC (L) | Scenes | Lure correct > Lure incorrect | -0.001 | 0.008 | -0.005 | 0.936 | 0.961 |
| PMERC (R) | Objects | Lure correct > Repeat | -0.009 | 0.007 | -0.068 | 0.214 | 0.804 |
| PMERC (R) | Objects | Lure correct > Lure incorrect | 0.005 | 0.008 | 0.039 | 0.540 | 0.884 |
| PMERC (R) | Scenes | Lure correct > Repeat | 0.002 | 0.007 | 0.016 | 0.770 | 0.825 |
| PMERC (R) | Scenes | Lure correct > Lure incorrect | 0.006 | 0.008 | 0.049 | 0.430 | 0.874 |
| PHC (L) | Objects | Lure correct > Repeat | -0.005 | 0.007 | -0.040 | 0.461 | 0.812 |
| PHC (L) | Objects | Lure correct > Lure incorrect | -0.001 | 0.008 | -0.009 | 0.888 | 0.997 |
| PHC (L) | Scenes | Lure correct > Repeat | -0.004 | 0.007 | -0.028 | 0.620 | 0.715 |
| PHC (L) | Scenes | Lure correct > Lure incorrect | 0.019 | 0.008 | 0.145 | 0.022 | 0.332 |
| PHC (R) | Objects | Lure correct > Repeat | -0.005 | 0.007 | -0.036 | 0.515 | 0.812 |
| PHC (R) | Objects | Lure correct > Lure incorrect | 0.000 | 0.008 | 0.002 | 0.973 | 0.997 |
| PHC (R) | Scenes | Lure correct > Repeat | -0.009 | 0.007 | -0.067 | 0.246 | 0.461 |
| PHC (R) | Scenes | Lure correct > Lure incorrect | 0.010 | 0.008 | 0.075 | 0.244 | 0.759 |
| aSUB (L) | Objects | Lure correct > Repeat | -0.009 | 0.007 | -0.073 | 0.195 | 0.804 |
| aSUB (L) | Objects | Lure correct > Lure incorrect | -0.010 | 0.008 | -0.078 | 0.234 | 0.884 |
| aSUB (L) | Scenes | Lure correct > Repeat | -0.005 | 0.007 | -0.043 | 0.462 | 0.554 |
| aSUB (L) | Scenes | Lure correct > Lure incorrect | 0.004 | 0.008 | 0.033 | 0.612 | 0.874 |
| aSUB (R) | Objects | Lure correct > Repeat | -0.009 | 0.007 | -0.067 | 0.229 | 0.804 |
| aSUB (R) | Objects | Lure correct > Lure incorrect | -0.009 | 0.008 | -0.074 | 0.256 | 0.884 |
| aSUB (R) | Scenes | Lure correct > Repeat | -0.015 | 0.007 | -0.117 | 0.042 | 0.218 |
| aSUB (R) | Scenes | Lure correct > Lure incorrect | -0.006 | 0.008 | -0.046 | 0.477 | 0.874 |
| aCA1 (L) | Objects | Lure correct > Repeat | -0.006 | 0.007 | -0.043 | 0.448 | 0.812 |
| aCA1 (L) | Objects | Lure correct > Lure incorrect | -0.003 | 0.008 | -0.023 | 0.729 | 0.997 |

| Region | Stimulus | Comparison | Estimated marginal mean for differences in Fisher's z-transformed r | Standard error | Cohen's d | Uncorrected p-value | Corrected p-value |
|---|---|---|---|---|---|---|---|
| aCA1 (L) | Scenes | Lure correct > Repeat | -0.007 | 0.007 | -0.057 | 0.333 | 0.475 |
| aCA1 (L) | Scenes | Lure correct > Lure incorrect | 0.002 | 0.008 | 0.014 | 0.832 | 0.959 |
| aCA1 (R) | Objects | Lure correct > Repeat | -0.007 | 0.007 | -0.055 | 0.324 | 0.812 |
| aCA1 (R) | Objects | Lure correct > Lure incorrect | -0.009 | 0.008 | -0.073 | 0.263 | 0.884 |
| aCA1 (R) | Scenes | Lure correct > Repeat | -0.003 | 0.007 | -0.020 | 0.733 | 0.814 |
| aCA1 (R) | Scenes | Lure correct > Lure incorrect | 0.012 | 0.008 | 0.096 | 0.145 | 0.759 |
| aCA3 (L) | Objects | Lure correct > Repeat | 0.000 | 0.007 | 0.002 | 0.965 | 0.965 |
| aCA3 (L) | Objects | Lure correct > Lure incorrect | 0.010 | 0.008 | 0.076 | 0.241 | 0.884 |
| aCA3 (L) | Scenes | Lure correct > Repeat | -0.016 | 0.007 | -0.126 | 0.029 | 0.218 |
| aCA3 (L) | Scenes | Lure correct > Lure incorrect | 0.001 | 0.008 | 0.008 | 0.895 | 0.959 |
| aCA3 (R) | Objects | Lure correct > Repeat | -0.030 | 0.007 | -0.236 | 0.000 | 0.001 |
| aCA3 (R) | Objects | Lure correct > Lure incorrect | -0.005 | 0.008 | -0.042 | 0.520 | 0.884 |
| aCA3 (R) | Scenes | Lure correct > Repeat | 0.023 | 0.007 | 0.181 | 0.002 | 0.041 |
| aCA3 (R) | Scenes | Lure correct > Lure incorrect | 0.034 | 0.008 | 0.266 | 0.000 | 0.001 |
| aDG (L) | Objects | Lure correct > Repeat | -0.009 | 0.007 | -0.071 | 0.208 | 0.804 |
| aDG (L) | Objects | Lure correct > Lure incorrect | -0.009 | 0.008 | -0.067 | 0.303 | 0.884 |
| aDG (L) | Scenes | Lure correct > Repeat | -0.011 | 0.007 | -0.086 | 0.136 | 0.333 |
| aDG (L) | Scenes | Lure correct > Lure incorrect | -0.005 | 0.008 | -0.037 | 0.570 | 0.874 |
| aDG (R) | Objects | Lure correct > Repeat | -0.021 | 0.007 | -0.163 | 0.004 | 0.045 |
| aDG (R) | Objects | Lure correct > Lure incorrect | -0.017 | 0.009 | -0.133 | 0.048 | 0.718 |
| aDG (R) | Scenes | Lure correct > Repeat | 0.000 | 0.008 | 0.002 | 0.970 | 0.987 |
| aDG (R) | Scenes | Lure correct > Lure incorrect | 0.011 | 0.009 | 0.084 | 0.208 | 0.759 |
| pSUB (L) | Objects | Lure correct > Repeat | -0.003 | 0.007 | -0.024 | 0.674 | 0.840 |
| pSUB (L) | Objects | Lure correct > Lure incorrect | 0.002 | 0.008 | 0.016 | 0.809 | 0.997 |
| pSUB (L) | Scenes | Lure correct > Repeat | 0.000 | 0.007 | -0.001 | 0.987 | 0.987 |
| pSUB (L) | Scenes | Lure correct > Lure incorrect | 0.014 | 0.008 | 0.108 | 0.093 | 0.697 |
| pSUB (R) | Objects | Lure correct > Repeat | -0.005 | 0.007 | -0.039 | 0.486 | 0.812 |
| pSUB (R) | Objects | Lure correct > Lure incorrect | -0.008 | 0.008 | -0.064 | 0.334 | 0.884 |
| pSUB (R) | Scenes | Lure correct > Repeat | -0.011 | 0.007 | -0.086 | 0.144 | 0.333 |
| pSUB (R) | Scenes | Lure correct > Lure incorrect | 0.009 | 0.008 | 0.068 | 0.300 | 0.818 |
| pCA1 (L) | Objects | Lure correct > Repeat | -0.002 | 0.007 | -0.019 | 0.728 | 0.840 |

| Region | Stimulus | Comparison | Estimated marginal mean for differences in Fisher's z-transformed r | Standard error | Cohen's d | Uncorrected p-value | Corrected p-value |
|---|---|---|---|---|---|---|---|
| pCA1 (L) | Objects | Lure correct > Lure incorrect | -0.002 | 0.008 | -0.017 | 0.799 | 0.997 |
| pCA1 (L) | Scenes | Lure correct > Repeat | -0.007 | 0.007 | -0.057 | 0.332 | 0.475 |
| pCA1 (L) | Scenes | Lure correct > Lure incorrect | 0.007 | 0.008 | 0.056 | 0.391 | 0.874 |
| pCA1 (R) | Objects | Lure correct > Repeat | -0.007 | 0.007 | -0.053 | 0.347 | 0.812 |
| pCA1 (R) | Objects | Lure correct > Lure incorrect | -0.015 | 0.008 | -0.116 | 0.079 | 0.788 |
| pCA1 (R) | Scenes | Lure correct > Repeat | -0.013 | 0.007 | -0.103 | 0.080 | 0.292 |
| pCA1 (R) | Scenes | Lure correct > Lure incorrect | 0.005 | 0.008 | 0.036 | 0.587 | 0.874 |
| pCA3 (L) | Objects | Lure correct > Repeat | -0.022 | 0.007 | -0.170 | 0.002 | 0.034 |
| pCA3 (L) | Objects | Lure correct > Lure incorrect | -0.006 | 0.008 | -0.045 | 0.486 | 0.884 |
| pCA3 (L) | Scenes | Lure correct > Repeat | -0.018 | 0.007 | -0.140 | 0.015 | 0.155 |
| pCA3 (L) | Scenes | Lure correct > Lure incorrect | 0.003 | 0.008 | 0.025 | 0.702 | 0.877 |
| pCA3 (R) | Objects | Lure correct > Repeat | -0.012 | 0.007 | -0.095 | 0.094 | 0.702 |
| pCA3 (R) | Objects | Lure correct > Lure incorrect | -0.032 | 0.008 | -0.251 | 0.000 | 0.004 |
| pCA3 (R) | Scenes | Lure correct > Repeat | -0.022 | 0.007 | -0.175 | 0.003 | 0.041 |
| pCA3 (R) | Scenes | Lure correct > Lure incorrect | -0.003 | 0.008 | -0.027 | 0.677 | 0.877 |
| pDG (L) | Objects | Lure correct > Repeat | 0.003 | 0.007 | 0.027 | 0.636 | 0.830 |
| pDG (L) | Objects | Lure correct > Lure incorrect | -0.010 | 0.008 | -0.076 | 0.243 | 0.884 |
| pDG (L) | Scenes | Lure correct > Repeat | -0.011 | 0.007 | -0.088 | 0.132 | 0.333 |
| pDG (L) | Scenes | Lure correct > Lure incorrect | 0.006 | 0.008 | 0.047 | 0.471 | 0.874 |
| pDG (R) | Objects | Lure correct > Repeat | -0.005 | 0.007 | -0.040 | 0.473 | 0.812 |
| pDG (R) | Objects | Lure correct > Lure incorrect | -0.007 | 0.008 | -0.058 | 0.368 | 0.884 |
| pDG (R) | Scenes | Lure correct > Repeat | -0.007 | 0.007 | -0.054 | 0.348 | 0.475 |
| pDG (R) | Scenes | Lure correct > Lure incorrect | 0.017 | 0.008 | 0.134 | 0.041 | 0.408 |
| TAIL (L) | Objects | Lure correct > Repeat | -0.004 | 0.007 | -0.030 | 0.581 | 0.829 |
| TAIL (L) | Objects | Lure correct > Lure incorrect | -0.007 | 0.008 | -0.056 | 0.370 | 0.884 |
| TAIL (L) | Scenes | Lure correct > Repeat | -0.006 | 0.007 | -0.046 | 0.410 | 0.532 |
| TAIL (L) | Scenes | Lure correct > Lure incorrect | 0.010 | 0.008 | 0.076 | 0.223 | 0.759 |
| TAIL (R) | Objects | Lure correct > Repeat | -0.006 | 0.007 | -0.049 | 0.370 | 0.812 |
| TAIL (R) | Objects | Lure correct > Lure incorrect | -0.006 | 0.008 | -0.045 | 0.471 | 0.884 |
| TAIL (R) | Scenes | Lure correct > Repeat | -0.012 | 0.007 | -0.094 | 0.093 | 0.292 |
| TAIL (R) | Scenes | Lure correct > Lure incorrect | 0.005 | 0.008 | 0.038 | 0.540 | 0.874 |

**Notes. '*a'* denotes anterior, '*p'* posterior hippocampus. ALERC: anterolateral entorhinal cortex; AMY: amygdala; B: bilateral; BA: Brodmann area; CA: cornu ammonis; DG: dentate gyrus; L: left; PHC: parahippocampal cortex; PMERC: posterior-medial ERC; R: right; SUB: subiculum.**

3. *Further details on univariate and multivariate cluster-level analysis in template space*

*3.1. Univariate repetition suppression*

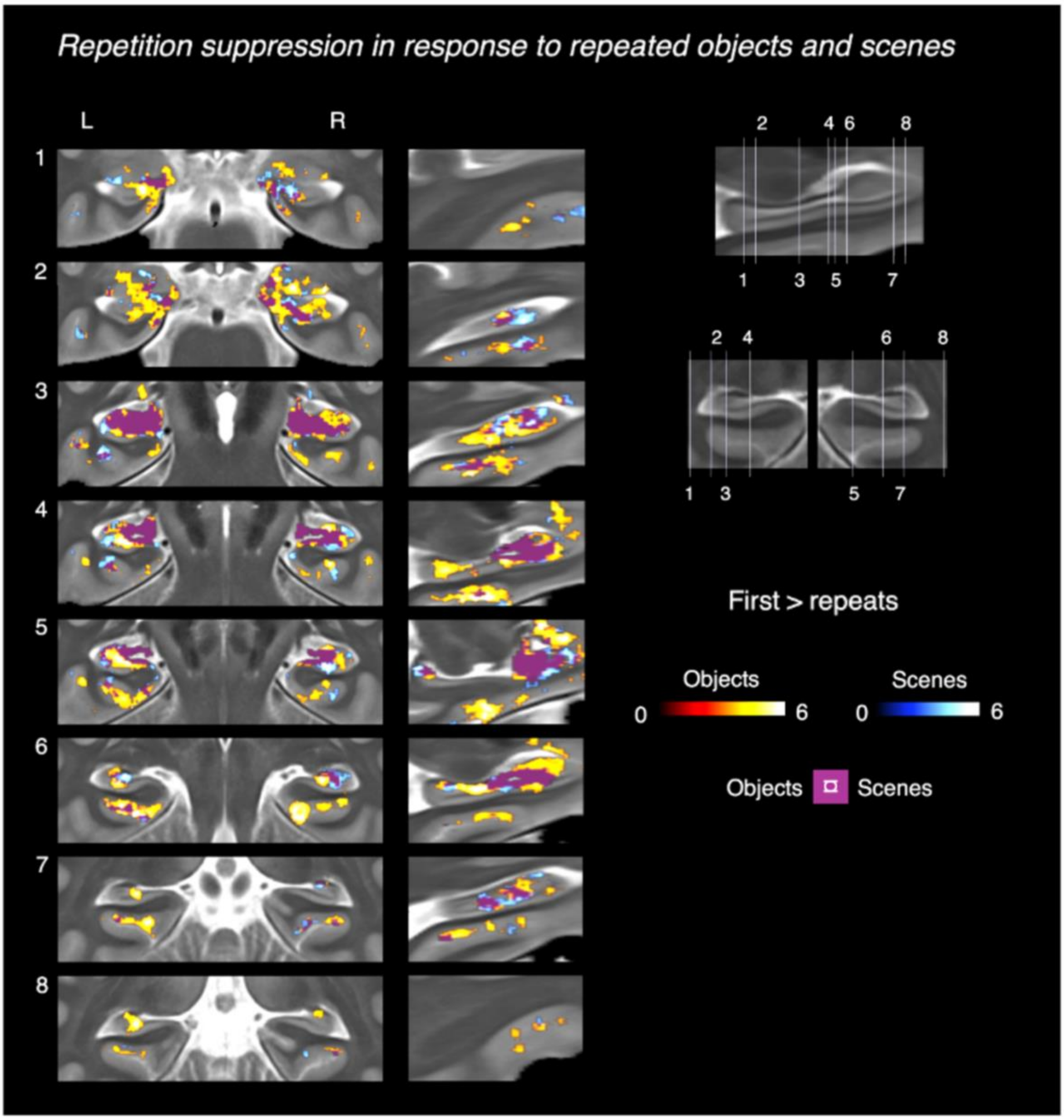


**Supplementary Figure 3. Repetition suppression effects for the contrast of first vs. repeated targets in a mass-univariate second-level analysis in sample template space. Overlap between object and scene repetition suppression effects are shown in purple. Effects were obtained based on uncorrected *p*-values on a voxel level and FDR correction at the cluster-level.**

*3.2. Trial type activity patterns of voxels from significant clusters in the univariate analysis*

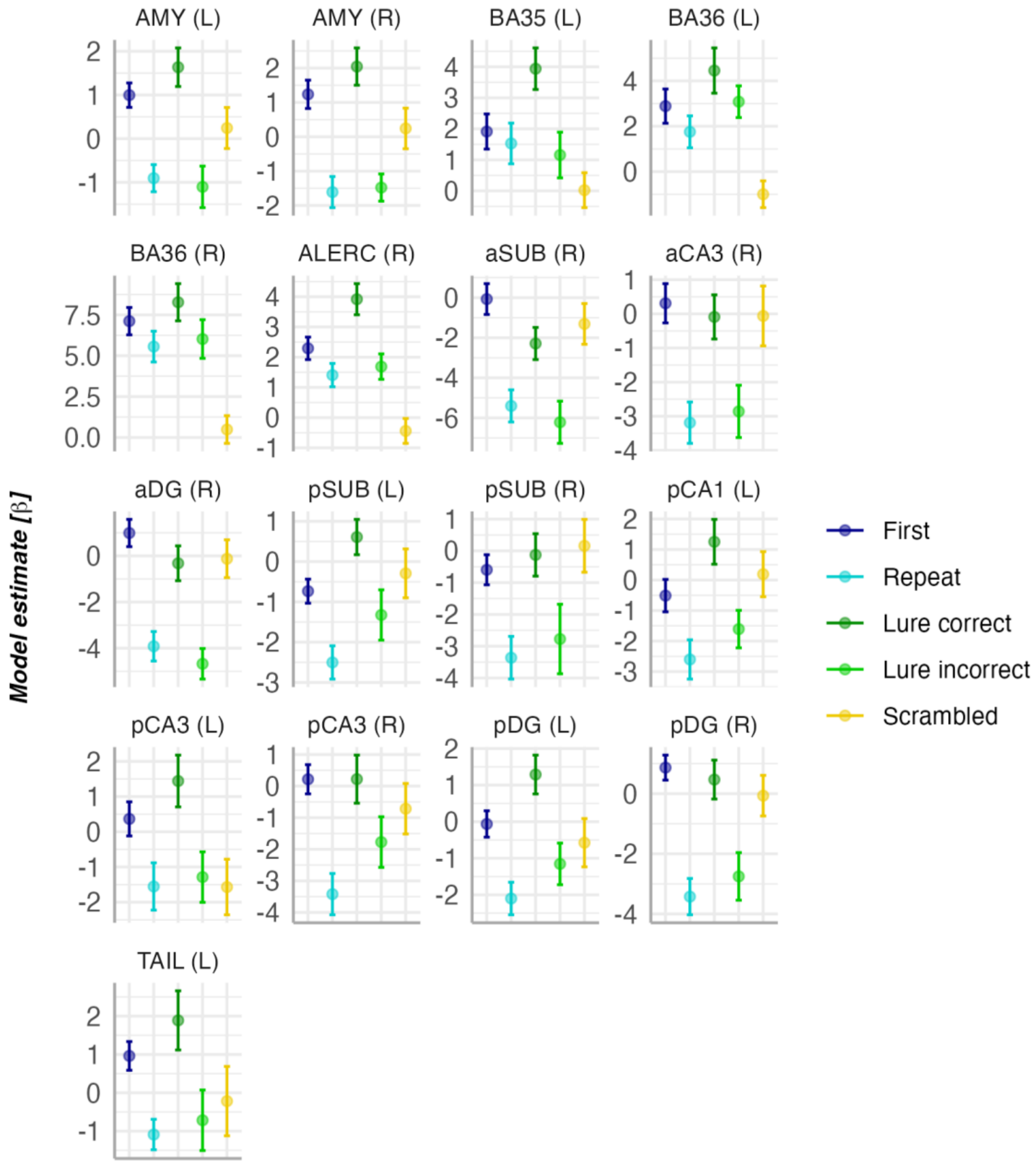


**Supplementary Figure 4. Model estimates for different trial types extracted from active clusters with object lure-related novelty responses (correct lures > repeats) presented in the main manuscript. Active clusters were masked with anatomically defined masks to obtain above-threshold voxels in each anatomical region of interest (where applicable). Trial types are first object presentations, repeated target presentations, correct object lures, incorrect scene lures, and scrambled images. Note. a: anterior; ALERC: anterolateral entorhinal cortex; AMY: amygdala; BA35: Brodmann area 35; BA36: Brodmann area 36; CA: cornu ammonis; DG: dentate gyrus; p: posterior; SUB: subiculum; TAIL: hippocampal tail**

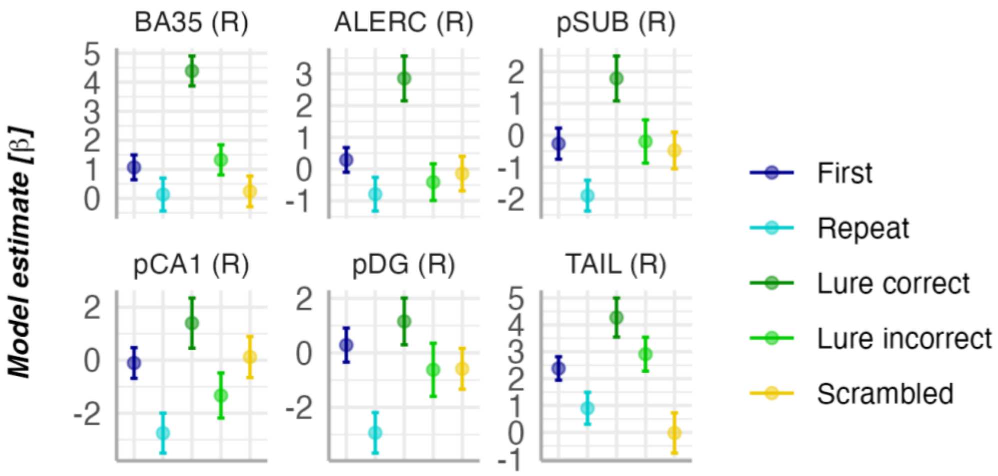


**Supplementary Figure 5. Model estimates for different trial types extracted from active clusters with lure-related novelty responses (correct lures > repeats) in the scene task presented in the main manuscript. Active clusters were masked with anatomically defined masks to obtain above-threshold voxels in each anatomical region of interest (where applicable). Different trial types are first scene presentations, repeated target presentations, correct scene lures, incorrect scene lures, and scrambled images. Note. BA35: Brodmann area 35; ALERC: anterolateral entorhinal cortex; pCA1: posterior cornu ammonis-1; pDG: posterior dentate gyrus; pSUB: posterior subiculum; TAIL: hippocampal tail.**

*3.3. Univariate lure-related novelty effects independent of behavioural outcome*

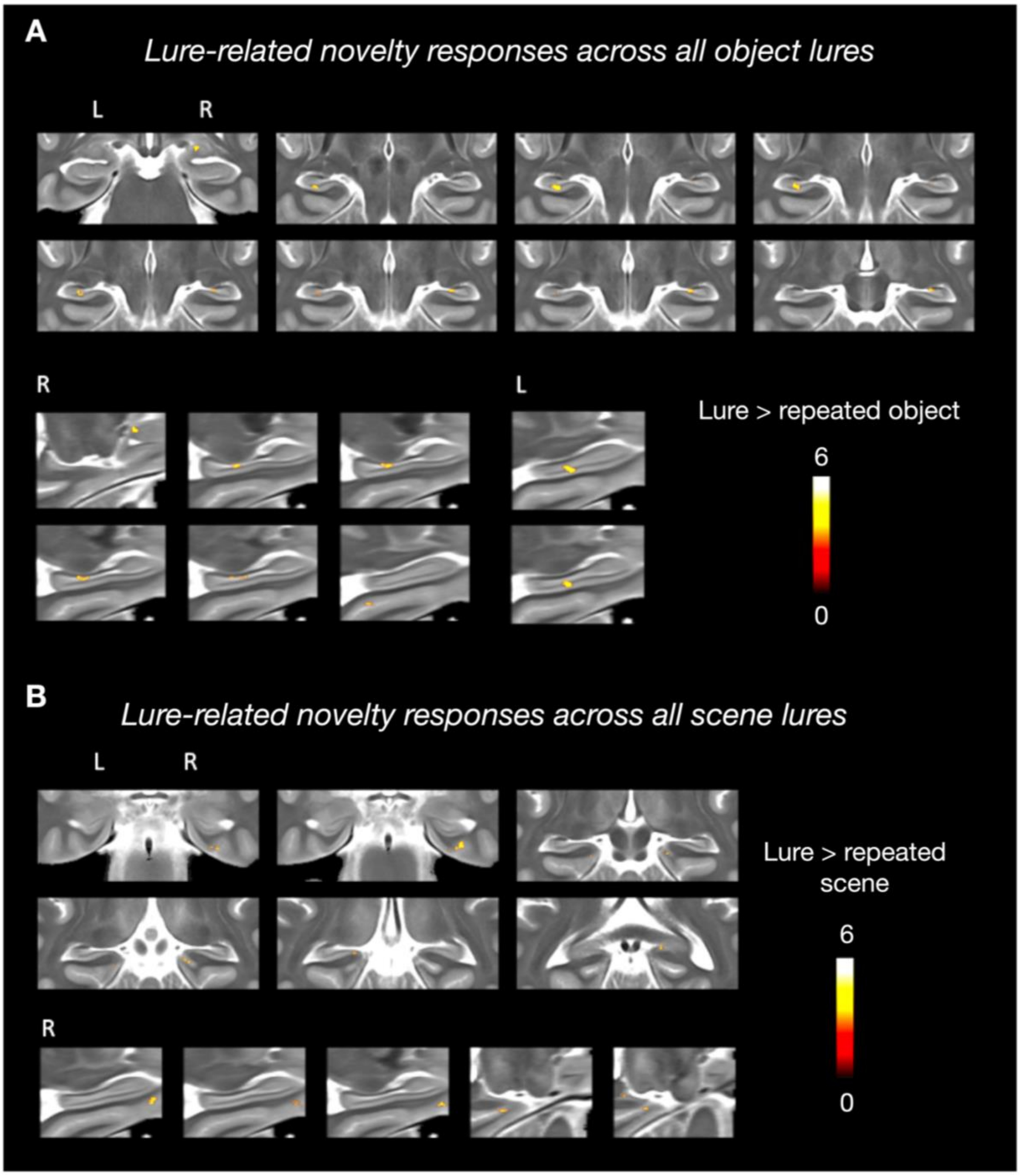


**Supplementary Figure 6. A. Results from the mass-univariate voxelwise analysis showing regions with greater activity during presentation of object lures (regardless of memory performance) compared to repeated object stimuli. B. Regions with greater activity during lures compared to repeats in the scene task. Results were thresholded based on uncorrected *p*-values on a voxel level and correction at the cluster-level.**

*3.4. Multivariate lure-related novelty effects independent of behavioural outcome*

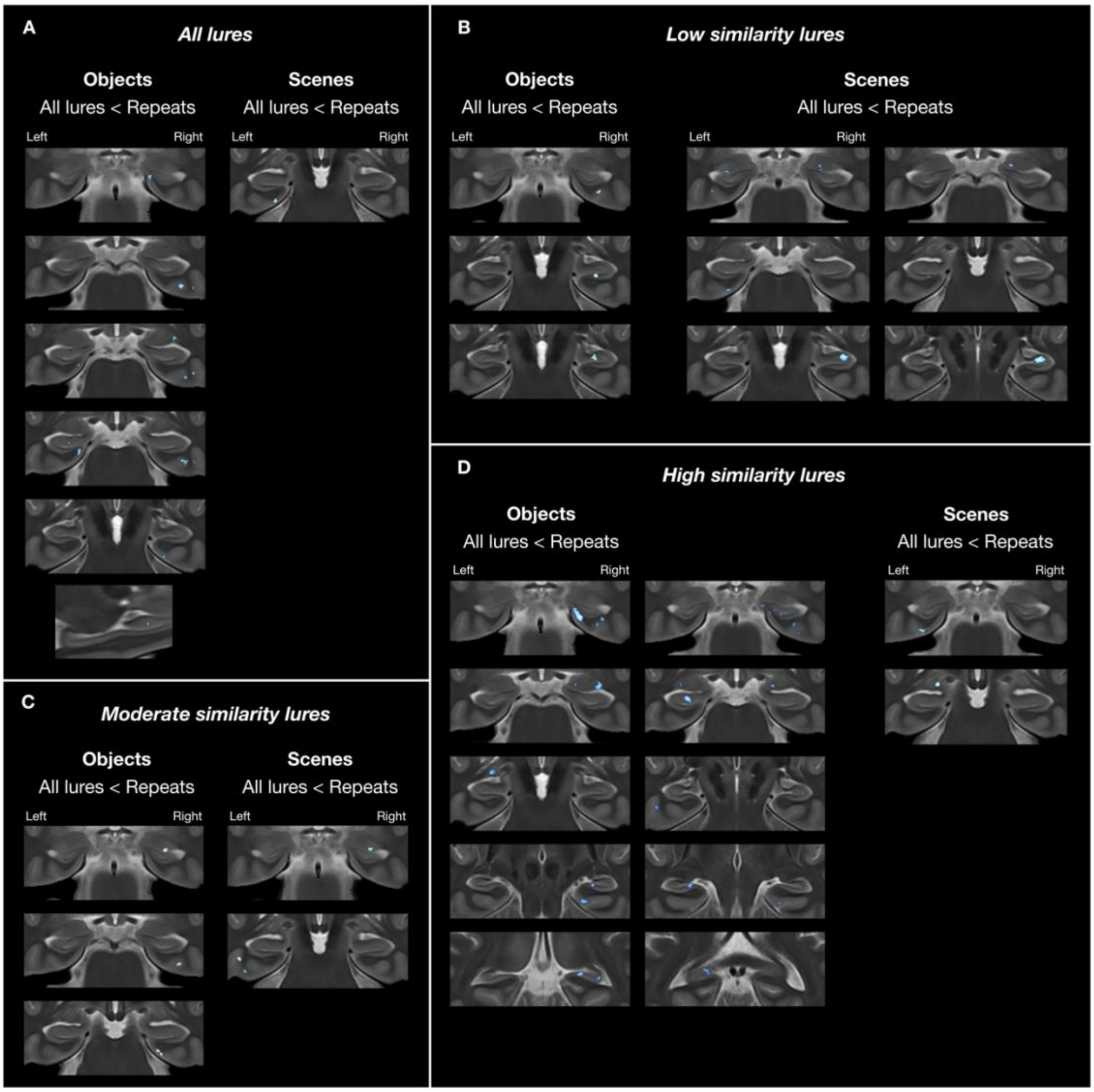


**Supplementary Figure 7. Searchlight analysis for multivariate pattern similarity for the contrast of all lures vs. repeats. Neural pattern refers to the similarity between the baseline (first target presentations) and a given trial type. A. Regions with lower pattern similarity during lures compared to repeats across all lures regardless of memory performance and target-lure similarity. B. Clusters with lower pattern similarity during low similarity lures (regardless of memory performance) compared to repeats. C. Clusters with lower pattern similarity during moderate similarity lures (regardless of memory performance) compared to repeats. D. Clusters with lower pattern similarity during high similarity lures (regardless of memory performance) compared to repeats.**

4. *Univariate brain responses as a function of stimulus similarity*

*4.1. Model curves by region*

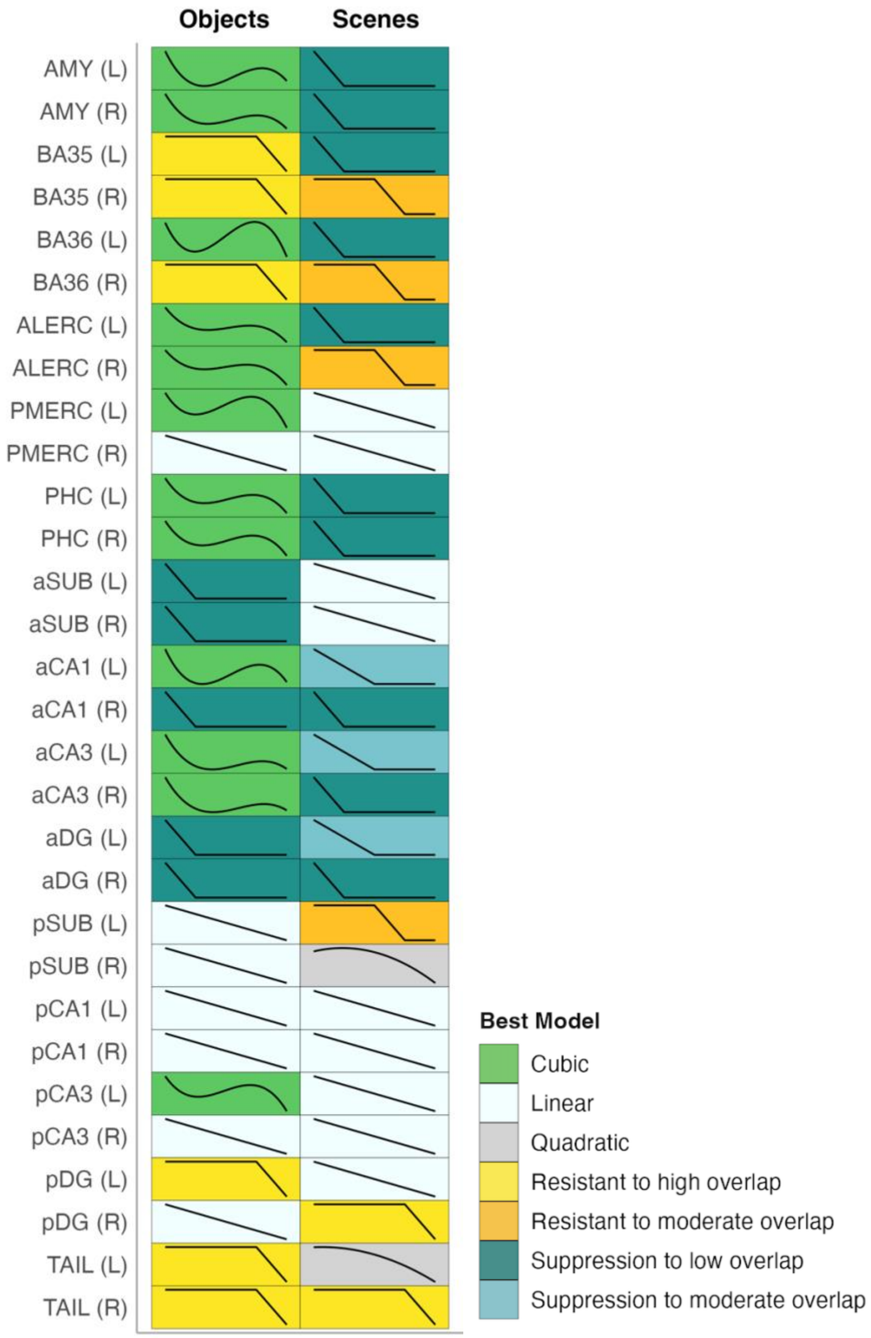


**Supplementary Figure 8. Overview of best models describing univariate brain activity across trial types (first presentation of a target, low similarity lure, moderate similarity lure, high similarity lure, repeated target). Regions that are highly sensitive to low levels of feature interference show strong suppression of activity from first to low similarity lows. Regions that are sensitive to moderate levels of feature interference show a linear decline of activity from first to low and moderate similarity lures, followed by similar activity levels to high similarity lures and repeats. Regions that show stable activity levels for first stimuli, low and moderate similarity lures followed by activity suppression during repeats are resistant to moderate feature overlap. Regions that show stable activity levels for first stimuli, low, moderate, and high similarity lures followed by activity suppression during repeats are resistant to high feature overlap. Notes. The prefix '*a'* denotes anterior hippocampus; '*p'*, posterior hippocampus. ALERC: anterolateral entorhinal cortex; AMY: amygdala; B: bilateral; BA: Brodmann area; CA: cornu ammonis; DG: dentate**

**gyrus; L: left; PHC: parahippocampal cortex; PMERC: posterior-medial entorhinal cortex; R: right; SUB: subiculum.**

With respect to the curve fitting findings, note that although right BA35 and BA36 were also best fit by a model where activity remains high across similarity bins, Bayesian Information Criterion favored a model with just an intercept, suggesting potential overfitting in the best AIC model.

### *4.2 Model selection statistics by region*

**Supplementary Table 5. Overview of the best and second-best models describing univariate brain activity as a function of target-lure feature overlap. The best model was selected based on Akaike Information Criterion.**

| ROI | Stimulus | Best model | Second best model | AIC of best model | AIC second best model - AIC best model |
|---|---|---|---|---|---|
| ALERC (L) | Objects | Cubic | Suppression to low overlap | 543.00 | 2.16 |
| ALERC (L) | Scenes | Suppression to low overlap | Quadratic | 574.80 | 0.78 |
| PMERC (L) | Objects | Cubic | Resistant to high overlap | 655.86 | 1.53 |
| PMERC (L) | Scenes | Linear | Resistant to moderate overlap | 669.42 | 0.72 |
| PHC (L) | Objects | Cubic | Suppression to low overlap | 658.88 | 4.32 |
| PHC (L) | Scenes | Suppression to low overlap | Linear | 686.72 | 0.44 |
| AMY (L) | Objects | Cubic | Suppression to low overlap | 603.61 | 4.86 |
| AMY (L) | Scenes | Suppression to low overlap | Quadratic | 596.70 | 4.92 |
| BA35 (L) | Objects | Resistant to high overlap | Linear | 575.70 | 2.53 |
| BA35 (L) | Scenes | Suppression to low overlap | Linear | 585.06 | 0.02 |
| BA36 (L) | Objects | Cubic | Resistant to high overlap | 491.91 | 10.15 |
| BA36 (L) | Scenes | Suppression to low overlap | Linear | 496.93 | 0.26 |
| aSUB (L) | Objects | Suppression to low overlap | Cubic | 556.93 | 0.18 |
| aSUB (L) | Scenes | Linear | Quadratic | 514.96 | 0.42 |
| aCA1 (L) | Objects | Cubic | Suppression to low overlap | 538.66 | 10.13 |
| aCA1 (L) | Scenes | Suppression to moderate overlap | Quadratic | 569.89 | 0.42 |
| aCA3 (L) | Objects | Cubic | Suppression to low overlap | 682.78 | 0.50 |
| aCA3 (L) | Scenes | Suppression to moderate overlap | Suppression to low overlap | 699.55 | 0.63 |
| aDG (L) | Objects | Suppression to low overlap | Cubic | 660.95 | 1.08 |
| aDG (L) | Scenes | Suppression to moderate overlap | Suppression to low overlap | 651.86 | 0.06 |
| pSUB (L) | Objects | Linear | Resistant to moderate overlap | 607.95 | 1.61 |
| pSUB (L) | Scenes | Resistant to moderate overlap | Quadratic | 630.35 | 1.57 |
| pCA1 (L) | Objects | Linear | Resistant to high overlap | 620.34 | 0.15 |
| pCA1 (L) | Scenes | Linear | Suppression to low overlap | 613.67 | 1.14 |
| pCA3 (L) | Objects | Cubic | Resistant to high overlap | 667.23 | 0.12 |
| pCA3 (L) | Scenes | Linear | Suppression to low overlap | 759.06 | 0.35 |
| pDG (L) | Objects | Resistant to high overlap | Cubic | 620.29 | 1.04 |
| pDG (L) | Scenes | Linear | Resistant to moderate overlap | 709.68 | 0.42 |
| TAIL (L) | Objects | Resistant to high overlap | Cubic | 566.44 | 2.23 |

| ROI | Stimulus | Best model | Second best model | AIC of best model | AIC second best model - AIC best model |
|---|---|---|---|---|---|
| TAIL (L) | Scenes | Quadratic | Linear | 612.66 | 0.15 |
| ALERC (R) | Objects | Cubic | Linear | 576.59 | 0.16 |
| ALERC (R) | Scenes | Resistant to moderate overlap | Cubic | 589.25 | 7.18 |
| PMERC (R) | Objects | Linear | Suppression to moderate overlap | 640.93 | 2.71 |
| PMERC (R) | Scenes | Linear | Quadratic | 686.93 | 1.48 |
| PHC (R) | Objects | Cubic | Suppression to low overlap | 727.36 | 1.50 |
| PHC (R) | Scenes | Suppression to low overlap | Cubic | 792.30 | 0.34 |
| AMY (R) | Objects | Cubic | Suppression to moderate overlap | 618.73 | 3.33 |
| AMY (R) | Scenes | Suppression to low overlap | Quadratic | 569.33 | 2.83 |
| BA35 (R) | Objects | Resistant to high overlap | Linear | 641.09 | 1.14 |
| BA35 (R) | Scenes | Resistant to moderate overlap | Linear | 619.91 | 1.12 |
| BA36 (R) | Objects | Resistant to high overlap | Cubic | 549.96 | 0.57 |
| BA36 (R) | Scenes | Resistant to moderate overlap | Linear | 504.25 | 2.21 |
| aSUB (R) | Objects | Suppression to low overlap | Cubic | 608.21 | 1.23 |
| aSUB (R) | Scenes | Linear | Quadratic | 607.33 | 1.17 |
| aCA1 (R) | Objects | Suppression to low overlap | Cubic | 604.84 | 2.49 |
| aCA1 (R) | Scenes | Suppression to low overlap | Cubic | 574.14 | 5.20 |
| aCA3 (R) | Objects | Cubic | Suppression to low overlap | 761.83 | 0.75 |
| aCA3 (R) | Scenes | Suppression to low overlap | Cubic | 753.88 | 5.89 |
| aDG (R) | Objects | Suppression to low overlap | Cubic | 713.06 | 1.21 |
| aDG (R) | Scenes | Suppression to low overlap | Cubic | 697.44 | 4.46 |
| pSUB (R) | Objects | Linear | Resistant to high overlap | 661.89 | 1.12 |
| pSUB (R) | Scenes | Quadratic | Cubic | 660.80 | 0.96 |
| pCA1 (R) | Objects | Linear | Resistant to low overlap | 689.94 | 1.44 |
| pCA1 (R) | Scenes | Linear | Resistant to high overlap | 658.50 | 1.25 |
| pCA3 (R) | Objects | Linear | Resistant to low overlap | 765.64 | 0.40 |
| pCA3 (R) | Scenes | Linear | Suppression to low overlap | 768.28 | 1.19 |
| pDG (R) | Objects | Linear | Resistant to low overlap | 739.33 | 1.11 |
| pDG (R) | Scenes | Resistant to high overlap | Quadratic | 724.59 | 0.61 |
| TAIL (R) | Objects | Resistant to high overlap | Linear | 676.69 | 2.13 |
| TAIL (R) | Scenes | Resistant to high overlap | Quadratic | 696.00 | 2.61 |
| ALERC (B) | Objects | Cubic | Linear | 508.55 | 2.53 |
| ALERC (B) | Scenes | Resistant to moderate overlap | Linear | 530.29 | 2.66 |

| ROI | Stimulus | Best model | Second best model | AIC of best model | AIC second best model - AIC best model |
|---|---|---|---|---|---|
| PMERC (B) | Objects | Cubic | Resistant to high overlap | 587.57 | 1.43 |
| PMERC (B) | Scenes | Linear | Quadratic | 603.90 | 1.85 |
| PHC (B) | Objects | Cubic | Suppression to low overlap | 664.05 | 5.71 |
| PHC (B) | Scenes | Suppression to low overlap | Cubic | 691.11 | 0.63 |
| AMY (B) | Objects | Cubic | Suppression to low overlap | 583.76 | 5.24 |
| AMY (B) | Scenes | Suppression to low overlap | Quadratic | 557.31 | 3.80 |
| BA35 (B) | Objects | Resistant to high overlap | Cubic | 551.82 | 1.57 |
| BA35 (B) | Scenes | Resistant to moderate overlap | Linear | 530.74 | 1.06 |
| BA36 (B) | Objects | Cubic | Resistant to high overlap | 487.41 | 5.37 |
| BA36 (B) | Scenes | Linear | Quadratic | 466.80 | 1.59 |
| aSUB (B) | Objects | Cubic | Suppression to low overlap | 535.77 | 0.16 |
| aSUB (B) | Scenes | Linear | Quadratic | 503.91 | 0.62 |
| aCA1 (B) | Objects | Cubic | Suppression to low overlap | 534.41 | 0.44 |
| aCA1 (B) | Scenes | Suppression to low overlap | Cubic | 531.35 | 2.97 |
| aCA3 (B) | Objects | Cubic | Suppression to low overlap | 667.18 | 1.34 |
| aCA3 (B) | Scenes | Suppression to low overlap | Cubic | 645.89 | 6.93 |
| aDG (B) | Objects | Cubic | Suppression to low overlap | 638.19 | 0.21 |
| aDG (B) | Scenes | Suppression to low overlap | Cubic | 620.40 | 2.96 |
| pSUB (B) | Objects | Linear | Cubic | 563.95 | 2.88 |
| pSUB (B) | Scenes | Quadratic | Cubic | 596.99 | 0.72 |
| pCA1 (B) | Objects | Linear | Cubic | 569.50 | 1.48 |
| pCA1 (B) | Scenes | Linear | Quadratic | 569.79 | 1.90 |
| pCA3 (B) | Objects | Linear | Cubic | 672.55 | 0.77 |
| pCA3 (B) | Scenes | Linear | Suppression to low overlap | 713.70 | 0.69 |
| pDG (B) | Objects | Linear | Cubic | 665.64 | 1.53 |
| pDG (B) | Scenes | Linear | Quadratic | 673.51 | 1.40 |
| TAIL (B) | Objects | Resistant to high overlap | Cubic | 596.47 | 2.80 |
| TAIL (B) | Scenes | Resistant to high overlap | Quadratic | 616.89 | 0.14 |

**Notes. The prefix '*a*' denotes anterior hippocampus; '*p*', posterior hippocampus. ALERC: anterolateral entorhinal cortex; AMY: amygdala; B: bilateral; BA: Brodmann area; CA: cornu ammonis; DG: dentate gyrus; L: left; PHC: parahippocampal cortex; PMERC: posterior-medial entorhinal cortex; R: right; SUB: subiculum.**

*4.3 Voxel-wise clusters of univariate lure-related novelty as a function of stimulus similarity*

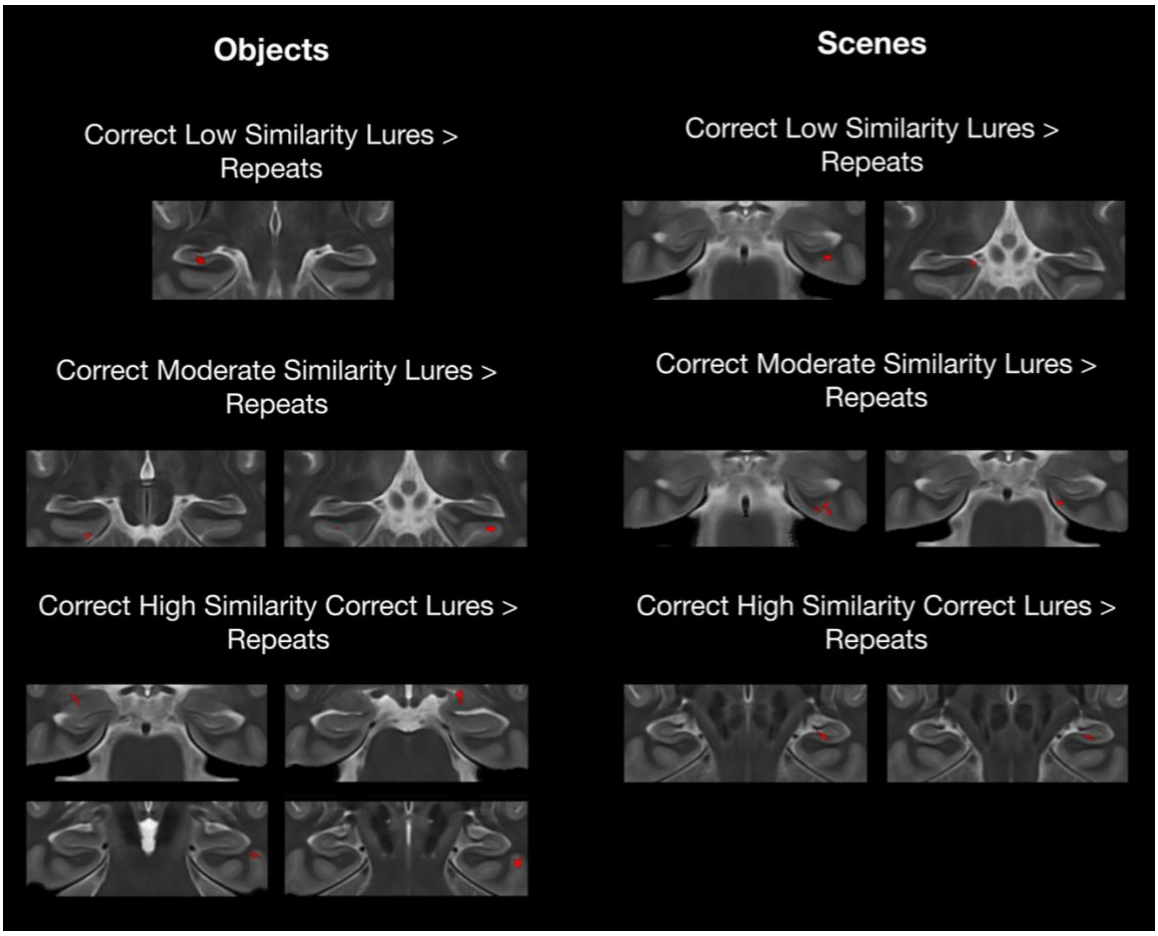


**Supplementary Figure 9. Univariate lure-related novelty responses for various similarity bins for objects and scenes.**

5. *Regions with higher pattern similarity between correct lures and firsts compared to incorrect and firsts in the scene mnemonic discrimination task*

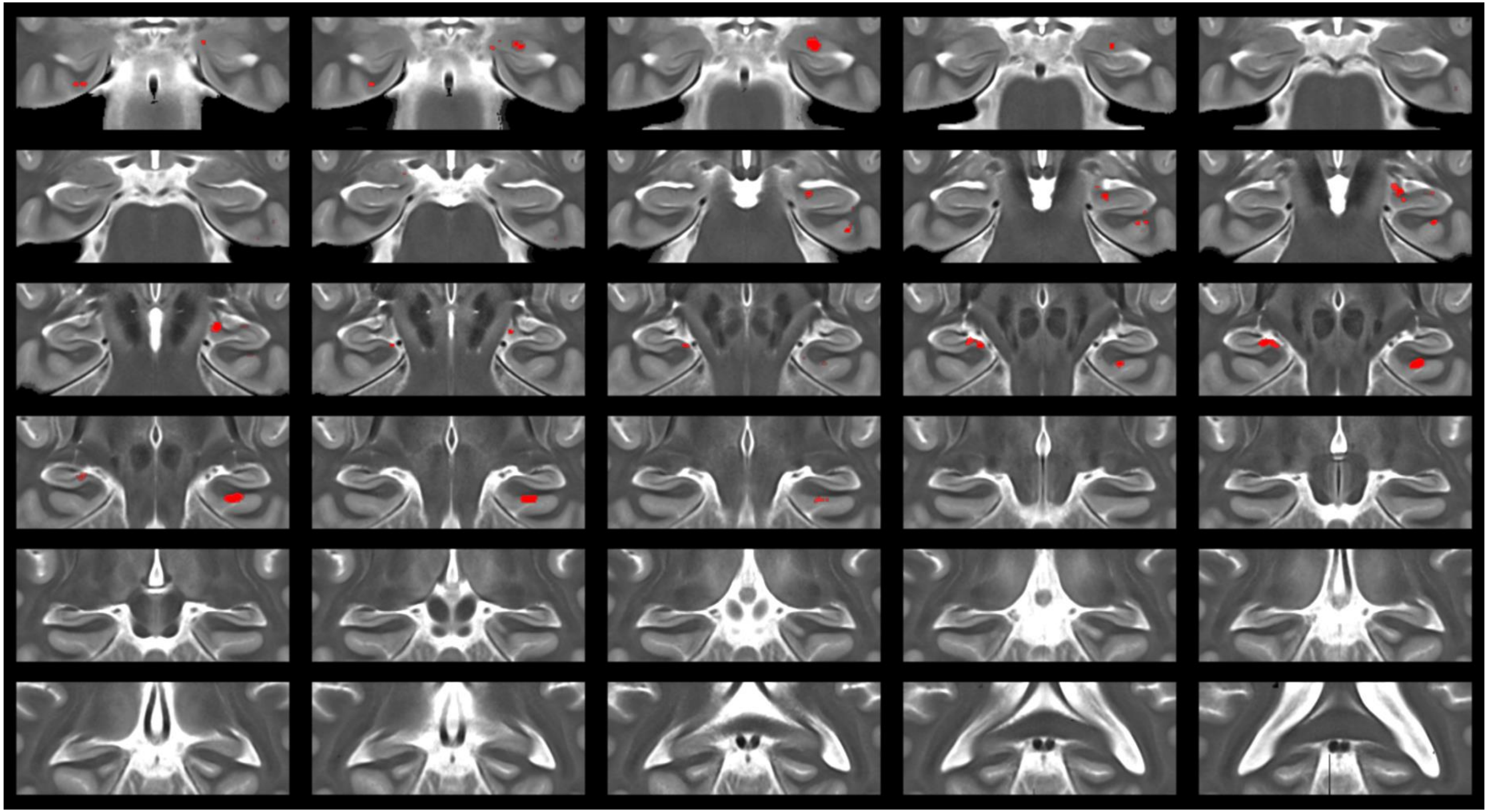

**Supplementary Figure 10. Higher pattern similarity during correct compared to incorrect scene lures when collapsing across all lure similarity bins.**

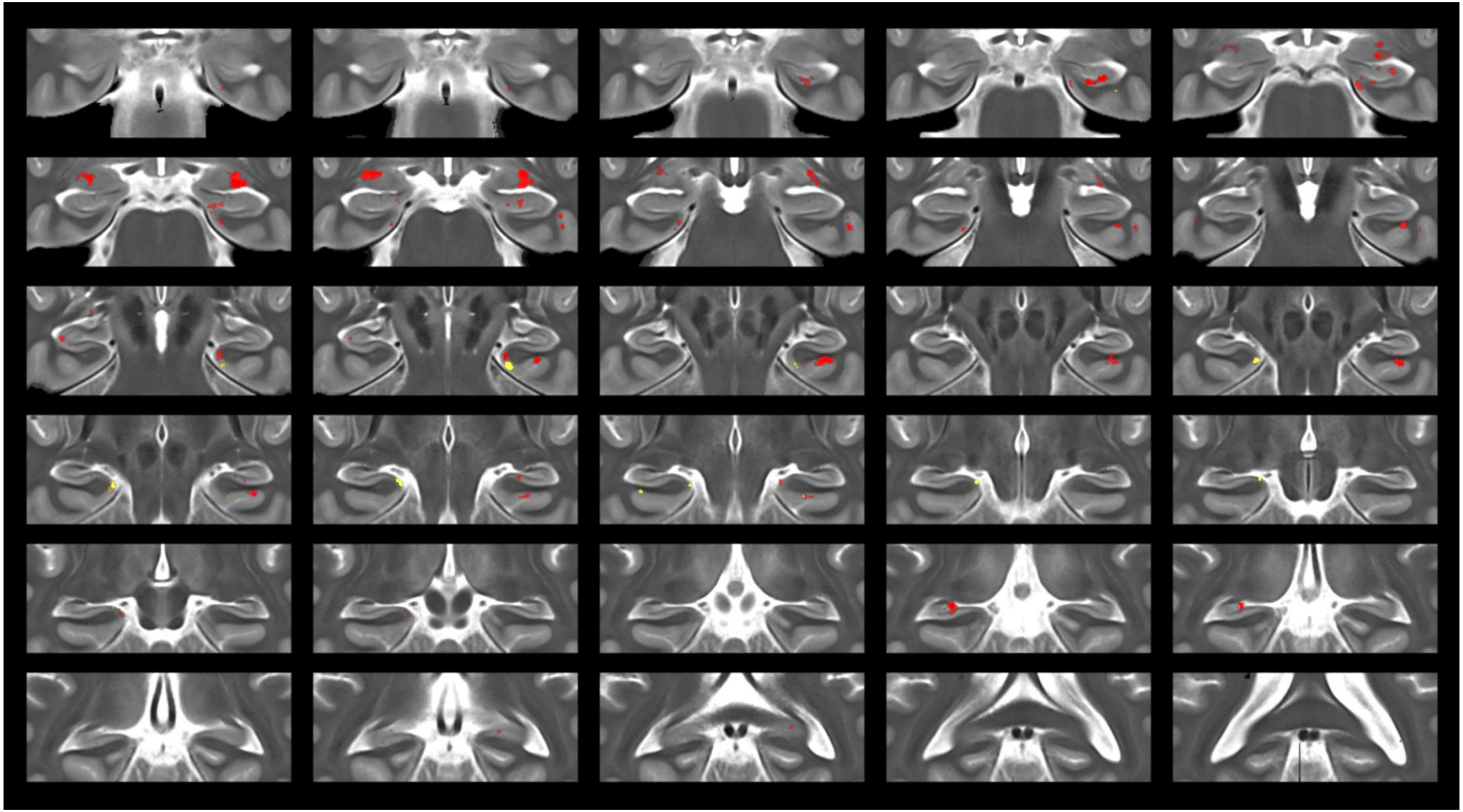

**Supplementary Figure 11. Higher pattern similarity during correct compared to incorrect scene lures for moderate similarity lures (yellow) and high similarity lures (red). There were no clusters with higher pattern similarity during correct vs. incorrect low similarity lures.**

6. *Brain-behaviour associations*

*6.1. Trial-by-trial neural similarity and lure discrimination*

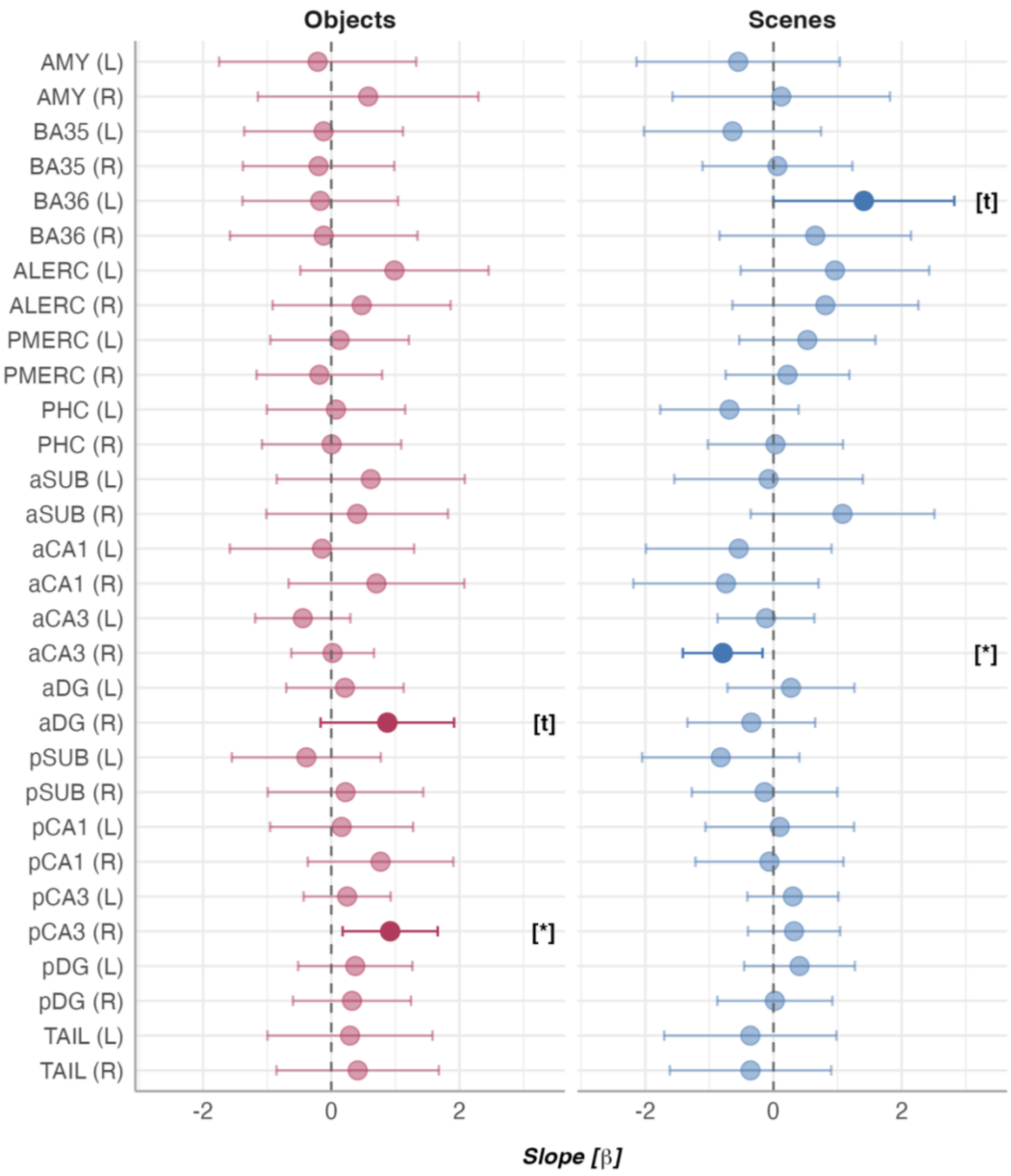


**Supplementary Figure 12. Association between trial-wise neural pattern similarity and lure discrimination. The slope here refers to the effect of neural pattern similarity between first target presentations and corresponding lure presentations on discrimination performance during lure trials, derived from a linear mixed model with response type (correct rejection of a lure vs. false alarm to a lure) and stimulus type (objects, scenes) as fixed effects and random intercepts for subject and stimulus. Notes. The prefix '*a*' denotes anterior hippocampus; '*p*', posterior hippocampus. ALERC: anterolateral entorhinal cortex; AMY: amygdala; B: bilateral; BA: Brodmann area; CA: cornu ammonis; DG: dentate gyrus; L: left; PHC: parahippocampal cortex; PMERC: posterior-medial entorhinal cortex; R: right; SUB: subiculum. $^{t}p<.10$, $^{*}p<.05$. Asterisks in square brackets refer to uncorrected *p*-values.**

### 6.2. Inter-individual differences in memory performance

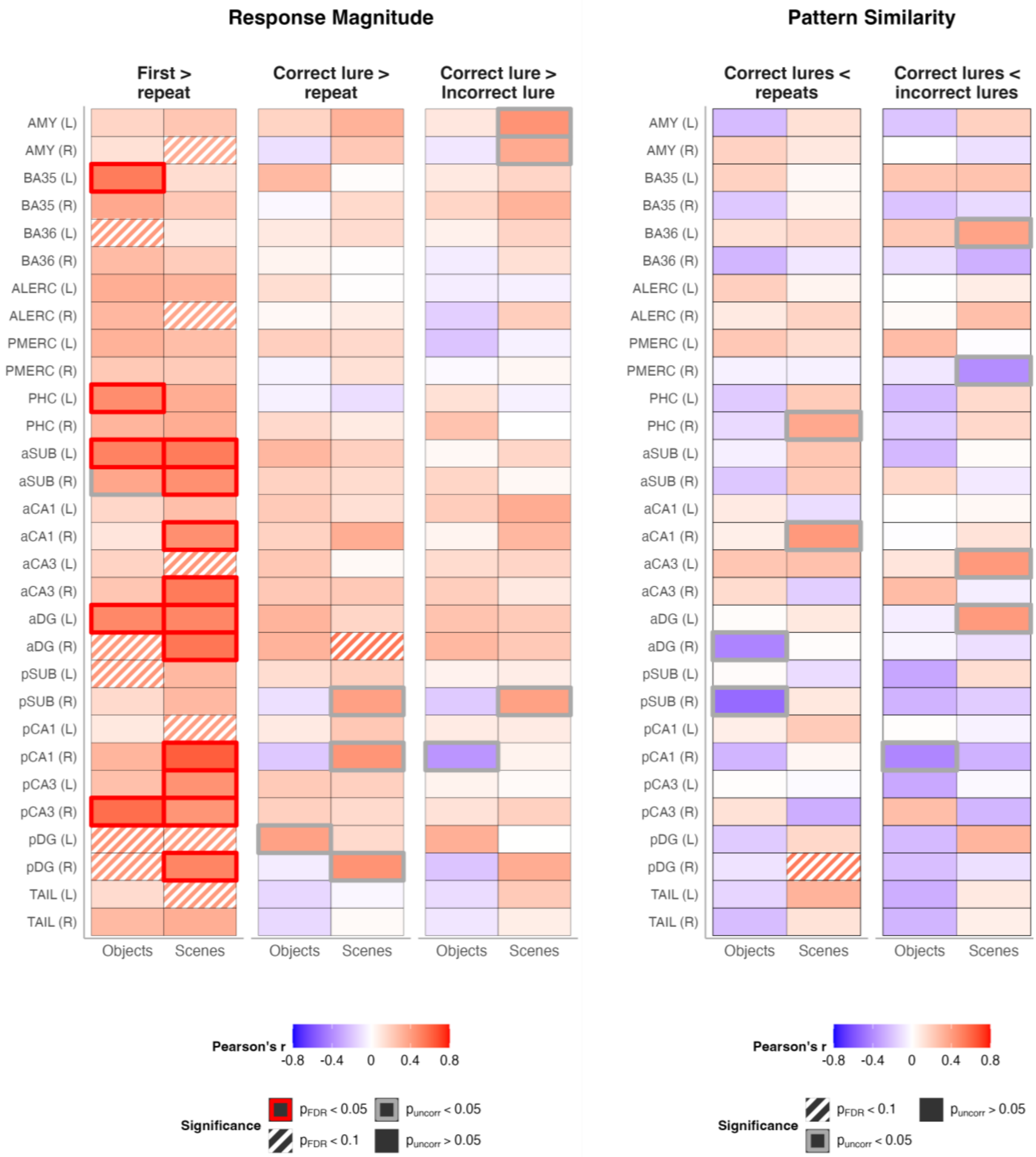


**Supplementary Figure 13. Overview of brain-behaviour correlations for univariate and multivariate measures of brain activity during the main contrasts of interest (correct lures vs. repeats; correct lures vs. incorrect lures) and for the sake of completeness, the univariate contrast of first vs. repeated targets. All *p*-values were FDR corrected at the contrast and stimulus type level. The behavioural measure was the corrected hit rate during the object and scene task, respectively. Univariate effects were contrast beta values from first-level analyses. Multivariate effects were differences in neural pattern similarity. Neural pattern similarity for repeats, correct lures, and incorrect lures here refers to the similarity between the baseline (first target presentations) and a given trial type. The correct lures < repeats contrast thus refers to lower pattern similarity between correct lures and firsts compared to pattern similarity between first and repeated targets.**

**Supplementary Table 6. Statistics for brain-behaviour correlations for univariate effects which represent contrast beta values from first-level analyses. *P*-values were corrected at the level of contrasts and stimulus type.**

| Region | Stimulus | Contrast | Pearson's *r* | DF | Uncorrected *p*-value | Corrected *p*-value |
|---|---|---|---|---|---|---|
| AMY (L) | Scenes | First > repeat | 0.237 | 30 | 0.192 | 0.230 |
| AMY (L) | Scenes | Correct lure > Incorrect lure | 0.442 | 31 | 0.010 | 0.303 |
| AMY (L) | Scenes | Correct lure > repeat | 0.321 | 31 | 0.068 | 0.342 |
| AMY (L) | Objects | First > repeat | 0.178 | 31 | 0.323 | 0.418 |
| AMY (L) | Objects | Correct lure > Incorrect lure | 0.100 | 31 | 0.580 | 0.869 |
| AMY (L) | Objects | Correct lure > repeat | 0.182 | 31 | 0.311 | 0.635 |
| AMY (R) | Scenes | First > repeat | 0.340 | 31 | 0.053 | 0.100 |
| AMY (R) | Scenes | Correct lure > Incorrect lure | 0.351 | 30 | 0.049 | 0.406 |
| AMY (R) | Scenes | Correct lure > repeat | 0.228 | 31 | 0.202 | 0.701 |
| AMY (R) | Objects | First > repeat | 0.128 | 31 | 0.478 | 0.512 |
| AMY (R) | Objects | Correct lure > Incorrect lure | -0.080 | 30 | 0.662 | 0.869 |
| AMY (R) | Objects | Correct lure > repeat | -0.106 | 31 | 0.558 | 0.782 |
| BA35 (L) | Scenes | First > repeat | 0.141 | 29 | 0.450 | 0.465 |
| BA35 (L) | Scenes | Correct lure > Incorrect lure | 0.174 | 29 | 0.348 | 0.691 |
| BA35 (L) | Scenes | Correct lure > repeat | 0.011 | 29 | 0.953 | 0.971 |
| BA35 (L) | Objects | First > repeat | 0.530 | 29 | 0.002 | 0.033 |
| BA35 (L) | Objects | Correct lure > Incorrect lure | 0.093 | 29 | 0.619 | 0.869 |
| BA35 (L) | Objects | Correct lure > repeat | 0.291 | 29 | 0.113 | 0.635 |
| BA35 (R) | Scenes | First > repeat | 0.228 | 29 | 0.217 | 0.251 |
| BA35 (R) | Scenes | Correct lure > Incorrect lure | 0.317 | 29 | 0.083 | 0.413 |
| BA35 (R) | Scenes | Correct lure > repeat | 0.157 | 29 | 0.399 | 0.701 |
| BA35 (R) | Objects | First > repeat | 0.358 | 28 | 0.052 | 0.126 |
| BA35 (R) | Objects | Correct lure > Incorrect lure | 0.173 | 29 | 0.352 | 0.844 |
| BA35 (R) | Objects | Correct lure > repeat | -0.027 | 29 | 0.885 | 0.885 |
| BA36 (L) | Scenes | First > repeat | 0.102 | 29 | 0.584 | 0.584 |
| BA36 (L) | Scenes | Correct lure > Incorrect lure | 0.178 | 29 | 0.338 | 0.691 |
| BA36 (L) | Scenes | Correct lure > repeat | 0.145 | 28 | 0.444 | 0.701 |
| BA36 (L) | Objects | First > repeat | 0.412 | 29 | 0.021 | 0.065 |
| BA36 (L) | Objects | Correct lure > Incorrect lure | 0.059 | 29 | 0.753 | 0.869 |
| BA36 (L) | Objects | Correct lure > repeat | 0.091 | 29 | 0.627 | 0.812 |
| BA36 (R) | Scenes | First > repeat | 0.212 | 28 | 0.260 | 0.278 |

| Region | Stimulus | Contrast | Pearson's *r* | DF | Uncorrected *p*-value | Corrected *p*-value |
|---|---|---|---|---|---|---|
| BA36 (R) | Scenes | Correct lure > Incorrect lure | 0.129 | 29 | 0.489 | 0.864 |
| BA36 (R) | Scenes | Correct lure > repeat | 0.007 | 28 | 0.969 | 0.971 |
| BA36 (R) | Objects | First > repeat | 0.282 | 29 | 0.124 | 0.196 |
| BA36 (R) | Objects | Correct lure > Incorrect lure | -0.066 | 29 | 0.725 | 0.869 |
| BA36 (R) | Objects | Correct lure > repeat | 0.045 | 29 | 0.812 | 0.879 |
| ALERC (L) | Scenes | First > repeat | 0.312 | 31 | 0.077 | 0.116 |
| ALERC (L) | Scenes | Correct lure > Incorrect lure | -0.049 | 30 | 0.789 | 0.967 |
| ALERC (L) | Scenes | Correct lure > repeat | 0.007 | 31 | 0.971 | 0.971 |
| ALERC (L) | Objects | First > repeat | 0.338 | 31 | 0.055 | 0.126 |
| ALERC (L) | Objects | Correct lure > Incorrect lure | -0.063 | 31 | 0.730 | 0.869 |
| ALERC (L) | Objects | Correct lure > repeat | 0.140 | 31 | 0.437 | 0.739 |
| ALERC (R) | Scenes | First > repeat | 0.366 | 30 | 0.040 | 0.079 |
| ALERC (R) | Scenes | Correct lure > Incorrect lure | 0.205 | 30 | 0.261 | 0.691 |
| ALERC (R) | Scenes | Correct lure > repeat | 0.075 | 30 | 0.682 | 0.853 |
| ALERC (R) | Objects | First > repeat | 0.299 | 31 | 0.091 | 0.171 |
| ALERC (R) | Objects | Correct lure > Incorrect lure | -0.163 | 31 | 0.366 | 0.844 |
| ALERC (R) | Objects | Correct lure > repeat | 0.029 | 31 | 0.874 | 0.885 |
| PMERC (L) | Scenes | First > repeat | 0.272 | 31 | 0.125 | 0.164 |
| PMERC (L) | Scenes | Correct lure > Incorrect lure | -0.047 | 31 | 0.793 | 0.967 |
| PMERC (L) | Scenes | Correct lure > repeat | 0.158 | 31 | 0.379 | 0.701 |
| PMERC (L) | Objects | First > repeat | 0.321 | 31 | 0.069 | 0.148 |
| PMERC (L) | Objects | Correct lure > Incorrect lure | -0.202 | 31 | 0.259 | 0.844 |
| PMERC (L) | Objects | Correct lure > repeat | 0.199 | 31 | 0.267 | 0.635 |
| PMERC (R) | Scenes | First > repeat | 0.218 | 30 | 0.230 | 0.255 |
| PMERC (R) | Scenes | Correct lure > Incorrect lure | 0.033 | 30 | 0.859 | 0.973 |
| PMERC (R) | Scenes | Correct lure > repeat | 0.128 | 31 | 0.479 | 0.718 |
| PMERC (R) | Objects | First > repeat | 0.228 | 30 | 0.210 | 0.287 |
| PMERC (R) | Objects | Correct lure > Incorrect lure | -0.020 | 31 | 0.913 | 0.913 |
| PMERC (R) | Objects | Correct lure > repeat | -0.041 | 31 | 0.821 | 0.879 |
| PHC (L) | Scenes | First > repeat | 0.340 | 29 | 0.061 | 0.100 |
| PHC (L) | Scenes | Correct lure > Incorrect lure | -0.046 | 29 | 0.806 | 0.967 |
| PHC (L) | Scenes | Correct lure > repeat | -0.114 | 29 | 0.543 | 0.740 |
| PHC (L) | Objects | First > repeat | 0.465 | 29 | 0.008 | 0.050 |

| Region | Stimulus | Contrast | Pearson's *r* | DF | Uncorrected *p*-value | Corrected *p*-value |
|---|---|---|---|---|---|---|
| PHC (L) | Objects | Correct lure > Incorrect lure | 0.125 | 28 | 0.509 | 0.869 |
| PHC (L) | Objects | Correct lure > repeat | -0.048 | 29 | 0.798 | 0.879 |
| PHC (R) | Scenes | First > repeat | 0.342 | 29 | 0.060 | 0.100 |
| PHC (R) | Scenes | Correct lure > Incorrect lure | 0.001 | 29 | 0.994 | 0.994 |
| PHC (R) | Scenes | Correct lure > repeat | 0.086 | 29 | 0.647 | 0.844 |
| PHC (R) | Objects | First > repeat | 0.309 | 29 | 0.091 | 0.171 |
| PHC (R) | Objects | Correct lure > Incorrect lure | 0.249 | 29 | 0.177 | 0.844 |
| PHC (R) | Objects | Correct lure > repeat | 0.158 | 29 | 0.397 | 0.739 |
| aSUB (L) | Scenes | First > repeat | 0.529 | 28 | 0.003 | 0.020 |
| aSUB (L) | Scenes | Correct lure > Incorrect lure | 0.170 | 28 | 0.368 | 0.691 |
| aSUB (L) | Scenes | Correct lure > repeat | 0.197 | 28 | 0.297 | 0.701 |
| aSUB (L) | Objects | First > repeat | 0.505 | 28 | 0.004 | 0.044 |
| aSUB (L) | Objects | Correct lure > Incorrect lure | 0.030 | 28 | 0.876 | 0.906 |
| aSUB (L) | Objects | Correct lure > repeat | 0.298 | 28 | 0.109 | 0.635 |
| aSUB (R) | Scenes | First > repeat | 0.455 | 27 | 0.013 | 0.044 |
| aSUB (R) | Scenes | Correct lure > Incorrect lure | 0.030 | 28 | 0.875 | 0.973 |
| aSUB (R) | Scenes | Correct lure > repeat | 0.148 | 28 | 0.435 | 0.701 |
| aSUB (R) | Objects | First > repeat | 0.367 | 28 | 0.046 | 0.125 |
| aSUB (R) | Objects | Correct lure > Incorrect lure | 0.173 | 28 | 0.360 | 0.844 |
| aSUB (R) | Objects | Correct lure > repeat | 0.190 | 28 | 0.316 | 0.635 |
| aCA1 (L) | Scenes | First > repeat | 0.265 | 28 | 0.156 | 0.195 |
| aCA1 (L) | Scenes | Correct lure > Incorrect lure | 0.344 | 27 | 0.068 | 0.406 |
| aCA1 (L) | Scenes | Correct lure > repeat | 0.130 | 27 | 0.503 | 0.718 |
| aCA1 (L) | Objects | First > repeat | 0.167 | 28 | 0.377 | 0.452 |
| aCA1 (L) | Objects | Correct lure > Incorrect lure | 0.207 | 28 | 0.272 | 0.844 |
| aCA1 (L) | Objects | Correct lure > repeat | 0.222 | 28 | 0.239 | 0.635 |
| aCA1 (R) | Scenes | First > repeat | 0.459 | 28 | 0.011 | 0.044 |
| aCA1 (R) | Scenes | Correct lure > Incorrect lure | 0.300 | 28 | 0.107 | 0.458 |
| aCA1 (R) | Scenes | Correct lure > repeat | 0.340 | 28 | 0.066 | 0.342 |
| aCA1 (R) | Objects | First > repeat | 0.108 | 28 | 0.571 | 0.591 |
| aCA1 (R) | Objects | Correct lure > Incorrect lure | 0.045 | 28 | 0.811 | 0.869 |
| aCA1 (R) | Objects | Correct lure > repeat | 0.189 | 28 | 0.318 | 0.635 |
| aCA3 (L) | Scenes | First > repeat | 0.428 | 27 | 0.021 | 0.054 |

| Region | Stimulus | Contrast | Pearson's *r* | DF | Uncorrected *p*-value | Corrected *p*-value |
|---|---|---|---|---|---|---|
| aCA3 (L) | Scenes | Correct lure > Incorrect lure | 0.175 | 28 | 0.355 | 0.691 |
| aCA3 (L) | Scenes | Correct lure > repeat | 0.026 | 28 | 0.891 | 0.971 |
| aCA3 (L) | Objects | First > repeat | 0.182 | 28 | 0.335 | 0.418 |
| aCA3 (L) | Objects | Correct lure > Incorrect lure | 0.148 | 28 | 0.435 | 0.869 |
| aCA3 (L) | Objects | Correct lure > repeat | 0.235 | 28 | 0.211 | 0.635 |
| aCA3 (R) | Scenes | First > repeat | 0.531 | 28 | 0.003 | 0.020 |
| aCA3 (R) | Scenes | Correct lure > Incorrect lure | 0.093 | 28 | 0.623 | 0.967 |
| aCA3 (R) | Scenes | Correct lure > repeat | 0.225 | 28 | 0.232 | 0.701 |
| aCA3 (R) | Objects | First > repeat | 0.238 | 28 | 0.206 | 0.287 |
| aCA3 (R) | Objects | Correct lure > Incorrect lure | 0.204 | 28 | 0.279 | 0.844 |
| aCA3 (R) | Objects | Correct lure > repeat | 0.228 | 28 | 0.225 | 0.635 |
| aDG (L) | Scenes | First > repeat | 0.491 | 28 | 0.006 | 0.029 |
| aDG (L) | Scenes | Correct lure > Incorrect lure | 0.217 | 27 | 0.257 | 0.691 |
| aDG (L) | Scenes | Correct lure > repeat | 0.173 | 28 | 0.362 | 0.701 |
| aDG (L) | Objects | First > repeat | 0.485 | 28 | 0.007 | 0.050 |
| aDG (L) | Objects | Correct lure > Incorrect lure | 0.250 | 28 | 0.183 | 0.844 |
| aDG (L) | Objects | Correct lure > repeat | 0.313 | 28 | 0.092 | 0.635 |
| aDG (R) | Scenes | First > repeat | 0.549 | 28 | 0.002 | 0.020 |
| aDG (R) | Scenes | Correct lure > Incorrect lure | 0.224 | 27 | 0.242 | 0.691 |
| aDG (R) | Scenes | Correct lure > repeat | 0.546 | 27 | 0.002 | 0.065 |
| aDG (R) | Objects | First > repeat | 0.417 | 28 | 0.022 | 0.065 |
| aDG (R) | Objects | Correct lure > Incorrect lure | 0.294 | 28 | 0.114 | 0.844 |
| aDG (R) | Objects | Correct lure > repeat | 0.314 | 28 | 0.091 | 0.635 |
| pSUB (L) | Scenes | First > repeat | 0.296 | 28 | 0.112 | 0.158 |
| pSUB (L) | Scenes | Correct lure > Incorrect lure | 0.074 | 28 | 0.696 | 0.967 |
| pSUB (L) | Scenes | Correct lure > repeat | 0.198 | 28 | 0.294 | 0.701 |
| pSUB (L) | Objects | First > repeat | 0.421 | 28 | 0.020 | 0.065 |
| pSUB (L) | Objects | Correct lure > Incorrect lure | 0.053 | 28 | 0.781 | 0.869 |
| pSUB (L) | Objects | Correct lure > repeat | 0.141 | 28 | 0.457 | 0.739 |
| pSUB (R) | Scenes | First > repeat | 0.293 | 28 | 0.116 | 0.158 |
| pSUB (R) | Scenes | Correct lure > Incorrect lure | 0.390 | 28 | 0.033 | 0.406 |
| pSUB (R) | Scenes | Correct lure > repeat | 0.390 | 28 | 0.033 | 0.249 |
| pSUB (R) | Objects | First > repeat | 0.153 | 27 | 0.428 | 0.475 |

| Region | Stimulus | Contrast | Pearson's *r* | DF | Uncorrected *p*-value | Corrected *p*-value |
|---|---|---|---|---|---|---|
| pSUB (R) | Objects | Correct lure > Incorrect lure | -0.182 | 28 | 0.337 | 0.844 |
| pSUB (R) | Objects | Correct lure > repeat | -0.107 | 28 | 0.574 | 0.782 |
| pCA1 (L) | Scenes | First > repeat | 0.398 | 28 | 0.029 | 0.065 |
| pCA1 (L) | Scenes | Correct lure > Incorrect lure | 0.082 | 28 | 0.665 | 0.967 |
| pCA1 (L) | Scenes | Correct lure > repeat | 0.229 | 28 | 0.223 | 0.701 |
| pCA1 (L) | Objects | First > repeat | 0.094 | 28 | 0.621 | 0.621 |
| pCA1 (L) | Objects | Correct lure > Incorrect lure | 0.082 | 28 | 0.666 | 0.869 |
| pCA1 (L) | Objects | Correct lure > repeat | 0.086 | 28 | 0.650 | 0.812 |
| pCA1 (R) | Scenes | First > repeat | 0.631 | 28 | 0.000 | 0.006 |
| pCA1 (R) | Scenes | Correct lure > Incorrect lure | 0.052 | 28 | 0.786 | 0.967 |
| pCA1 (R) | Scenes | Correct lure > repeat | 0.434 | 28 | 0.017 | 0.165 |
| pCA1 (R) | Objects | First > repeat | 0.307 | 28 | 0.098 | 0.174 |
| pCA1 (R) | Objects | Correct lure > Incorrect lure | -0.364 | 28 | 0.048 | 0.844 |
| pCA1 (R) | Objects | Correct lure > repeat | -0.189 | 28 | 0.316 | 0.635 |
| pCA3 (L) | Scenes | First > repeat | 0.447 | 28 | 0.013 | 0.044 |
| pCA3 (L) | Scenes | Correct lure > Incorrect lure | 0.022 | 28 | 0.910 | 0.975 |
| pCA3 (L) | Scenes | Correct lure > repeat | 0.196 | 28 | 0.298 | 0.701 |
| pCA3 (L) | Objects | First > repeat | 0.263 | 28 | 0.161 | 0.241 |
| pCA3 (L) | Objects | Correct lure > Incorrect lure | 0.052 | 28 | 0.785 | 0.869 |
| pCA3 (L) | Objects | Correct lure > repeat | 0.220 | 28 | 0.242 | 0.635 |
| pCA3 (R) | Scenes | First > repeat | 0.436 | 28 | 0.016 | 0.048 |
| pCA3 (R) | Scenes | Correct lure > Incorrect lure | 0.192 | 28 | 0.310 | 0.691 |
| pCA3 (R) | Scenes | Correct lure > repeat | 0.153 | 28 | 0.418 | 0.701 |
| pCA3 (R) | Objects | First > repeat | 0.577 | 28 | 0.001 | 0.025 |
| pCA3 (R) | Objects | Correct lure > Incorrect lure | 0.121 | 28 | 0.525 | 0.869 |
| pCA3 (R) | Objects | Correct lure > repeat | 0.195 | 28 | 0.301 | 0.635 |
| pDG (L) | Scenes | First > repeat | 0.396 | 28 | 0.030 | 0.065 |
| pDG (L) | Scenes | Correct lure > Incorrect lure | 0.002 | 28 | 0.993 | 0.994 |
| pDG (L) | Scenes | Correct lure > repeat | 0.155 | 28 | 0.413 | 0.701 |
| pDG (L) | Objects | First > repeat | 0.439 | 28 | 0.015 | 0.065 |
| pDG (L) | Objects | Correct lure > Incorrect lure | 0.333 | 28 | 0.073 | 0.844 |
| pDG (L) | Objects | Correct lure > repeat | 0.387 | 28 | 0.034 | 0.635 |
| pDG (R) | Scenes | First > repeat | 0.497 | 28 | 0.005 | 0.029 |

| Region | Stimulus | Contrast | Pearson's *r* | DF | Uncorrected *p*-value | Corrected *p*-value |
|---|---|---|---|---|---|---|
| pDG (R) | Scenes | Correct lure > Incorrect lure | 0.345 | 27 | 0.067 | 0.406 |
| pDG (R) | Scenes | Correct lure > repeat | 0.443 | 27 | 0.016 | 0.165 |
| pDG (R) | Objects | First > repeat | 0.419 | 28 | 0.021 | 0.065 |
| pDG (R) | Objects | Correct lure > Incorrect lure | -0.200 | 28 | 0.289 | 0.844 |
| pDG (R) | Objects | Correct lure > repeat | -0.066 | 28 | 0.731 | 0.877 |
| TAIL (L) | Scenes | First > repeat | 0.411 | 29 | 0.022 | 0.054 |
| TAIL (L) | Scenes | Correct lure > Incorrect lure | 0.220 | 28 | 0.243 | 0.691 |
| TAIL (L) | Scenes | Correct lure > repeat | -0.029 | 29 | 0.877 | 0.971 |
| TAIL (L) | Objects | First > repeat | 0.149 | 29 | 0.424 | 0.475 |
| TAIL (L) | Objects | Correct lure > Incorrect lure | -0.118 | 29 | 0.529 | 0.869 |
| TAIL (L) | Objects | Correct lure > repeat | -0.136 | 28 | 0.474 | 0.739 |
| TAIL (R) | Scenes | First > repeat | 0.337 | 29 | 0.063 | 0.100 |
| TAIL (R) | Scenes | Correct lure > Incorrect lure | 0.073 | 29 | 0.695 | 0.967 |
| TAIL (R) | Scenes | Correct lure > repeat | 0.022 | 29 | 0.905 | 0.971 |
| TAIL (R) | Objects | First > repeat | 0.287 | 29 | 0.117 | 0.195 |
| TAIL (R) | Objects | Correct lure > Incorrect lure | -0.085 | 29 | 0.651 | 0.869 |
| TAIL (R) | Objects | Correct lure > repeat | -0.128 | 29 | 0.493 | 0.739 |

**Notes. The prefix '*a'* denotes anterior hippocampus; '*p'*, posterior hippocampus. ALERC: anterolateral entorhinal cortex; AMY: amygdala; B: bilateral; BA: Brodmann area; CA: cornu ammonis; DG: dentate gyrus; L: left; PHC: parahippocampal cortex; PMERC: posterior-medial entorhinal cortex; R: right; SUB: subiculum.**

**Supplementary Table 7. Brain-behaviour correlations for multivariate effects. Multivariate effects were differences in neural pattern similarity. Neural pattern similarity for repeats, correct lures, and incorrect lures here refers to the similarity between the baseline (first target presentations) and a given trial type. The correct lures < repeats contrast thus refers to lower pattern similarity between correct lures and firsts compared to pattern similarity between first and repeated targets. *P*-values were corrected at the level of contrasts and stimulus type.**

| Region | Stimulus | Contrast | Pearson's r | DF | Uncorrected p-value | Corrected p-value |
|---|---|---|---|---|---|---|
| AMY (L) | Objects | Correct lures < repeats | -0.231 | 31 | 0.196 | 0.672 |
| AMY (L) | Objects | Correct lures < incorrect lures | -0.198 | 31 | 0.270 | 0.476 |
| AMY (L) | Scenes | Correct lures < repeats | 0.126 | 32 | 0.477 | 0.848 |
| AMY (L) | Scenes | Correct lures < incorrect lures | 0.194 | 32 | 0.273 | 0.744 |
| AMY (R) | Objects | Correct lures < repeats | 0.187 | 30 | 0.306 | 0.672 |
| AMY (R) | Objects | Correct lures < incorrect lures | -0.001 | 31 | 0.994 | 0.994 |
| AMY (R) | Scenes | Correct lures < repeats | 0.095 | 32 | 0.592 | 0.848 |
| AMY (R) | Scenes | Correct lures < incorrect lures | -0.106 | 32 | 0.551 | 0.868 |
| BA35 (L) | Objects | Correct lures < repeats | 0.190 | 29 | 0.307 | 0.672 |
| BA35 (L) | Objects | Correct lures < incorrect lures | 0.238 | 30 | 0.190 | 0.446 |
| BA35 (L) | Scenes | Correct lures < repeats | 0.028 | 31 | 0.875 | 0.938 |
| BA35 (L) | Scenes | Correct lures < incorrect lures | 0.251 | 31 | 0.159 | 0.529 |
| BA35 (R) | Objects | Correct lures < repeats | -0.188 | 31 | 0.294 | 0.672 |
| BA35 (R) | Objects | Correct lures < incorrect lures | -0.206 | 31 | 0.250 | 0.469 |
| BA35 (R) | Scenes | Correct lures < repeats | 0.047 | 30 | 0.799 | 0.924 |
| BA35 (R) | Scenes | Correct lures < incorrect lures | -0.127 | 31 | 0.482 | 0.868 |
| BA36 (L) | Objects | Correct lures < repeats | 0.127 | 31 | 0.481 | 0.779 |
| BA36 (L) | Objects | Correct lures < incorrect lures | 0.225 | 31 | 0.208 | 0.446 |
| BA36 (L) | Scenes | Correct lures < repeats | 0.158 | 31 | 0.380 | 0.835 |
| BA36 (L) | Scenes | Correct lures < incorrect lures | 0.380 | 31 | 0.029 | 0.219 |
| BA36 (R) | Objects | Correct lures < repeats | -0.252 | 32 | 0.151 | 0.672 |
| BA36 (R) | Objects | Correct lures < incorrect lures | -0.115 | 32 | 0.517 | 0.776 |
| BA36 (R) | Scenes | Correct lures < repeats | -0.080 | 31 | 0.657 | 0.857 |
| BA36 (R) | Scenes | Correct lures < incorrect lures | -0.272 | 31 | 0.126 | 0.529 |
| ALERC (L) | Objects | Correct lures < repeats | 0.199 | 30 | 0.276 | 0.672 |
| ALERC (L) | Objects | Correct lures < incorrect lures | 0.005 | 30 | 0.978 | 0.994 |
| ALERC (L) | Scenes | Correct lures < repeats | 0.046 | 31 | 0.799 | 0.924 |
| ALERC (L) | Scenes | Correct lures < incorrect lures | 0.078 | 31 | 0.667 | 0.881 |
| ALERC (R) | Objects | Correct lures < repeats | 0.089 | 29 | 0.635 | 0.848 |

| Region | Stimulus | Contrast | Pearson's r | DF | Uncorrected p-value | Corrected p-value |
|---|---|---|---|---|---|---|
| ALERC (R) | Objects | Correct lures < incorrect lures | 0.022 | 31 | 0.905 | 0.994 |
| ALERC (R) | Scenes | Correct lures < repeats | 0.180 | 31 | 0.315 | 0.835 |
| ALERC (R) | Scenes | Correct lures < incorrect lures | 0.266 | 31 | 0.135 | 0.529 |
| PMERC (L) | Objects | Correct lures < repeats | 0.229 | 30 | 0.208 | 0.672 |
| PMERC (L) | Objects | Correct lures < incorrect lures | 0.277 | 31 | 0.119 | 0.446 |
| PMERC (L) | Scenes | Correct lures < repeats | 0.143 | 31 | 0.427 | 0.848 |
| PMERC (L) | Scenes | Correct lures < incorrect lures | -0.009 | 31 | 0.961 | 0.961 |
| PMERC (R) | Objects | Correct lures < repeats | -0.048 | 30 | 0.796 | 0.884 |
| PMERC (R) | Objects | Correct lures < incorrect lures | -0.081 | 30 | 0.659 | 0.898 |
| PMERC (R) | Scenes | Correct lures < repeats | -0.046 | 30 | 0.801 | 0.924 |
| PMERC (R) | Scenes | Correct lures < incorrect lures | -0.394 | 31 | 0.023 | 0.219 |
| PHC (L) | Objects | Correct lures < repeats | -0.181 | 31 | 0.314 | 0.672 |
| PHC (L) | Objects | Correct lures < incorrect lures | -0.240 | 31 | 0.178 | 0.446 |
| PHC (L) | Scenes | Correct lures < repeats | 0.215 | 30 | 0.237 | 0.741 |
| PHC (L) | Scenes | Correct lures < incorrect lures | 0.155 | 30 | 0.397 | 0.851 |
| PHC (R) | Objects | Correct lures < repeats | -0.125 | 29 | 0.502 | 0.779 |
| PHC (R) | Objects | Correct lures < incorrect lures | -0.171 | 29 | 0.357 | 0.594 |
| PHC (R) | Scenes | Correct lures < repeats | 0.359 | 29 | 0.047 | 0.473 |
| PHC (R) | Scenes | Correct lures < incorrect lures | 0.162 | 29 | 0.383 | 0.851 |
| aSUB (L) | Objects | Correct lures < repeats | -0.054 | 28 | 0.776 | 0.884 |
| aSUB (L) | Objects | Correct lures < incorrect lures | -0.243 | 28 | 0.196 | 0.446 |
| aSUB (L) | Scenes | Correct lures < repeats | 0.239 | 28 | 0.203 | 0.741 |
| aSUB (L) | Scenes | Correct lures < incorrect lures | 0.019 | 29 | 0.921 | 0.952 |
| aSUB (R) | Objects | Correct lures < repeats | -0.192 | 29 | 0.300 | 0.672 |
| aSUB (R) | Objects | Correct lures < incorrect lures | 0.161 | 29 | 0.387 | 0.610 |
| aSUB (R) | Scenes | Correct lures < repeats | 0.221 | 29 | 0.232 | 0.741 |
| aSUB (R) | Scenes | Correct lures < incorrect lures | -0.074 | 29 | 0.693 | 0.881 |
| aCA1 (L) | Objects | Correct lures < repeats | 0.086 | 28 | 0.650 | 0.848 |
| aCA1 (L) | Objects | Correct lures < incorrect lures | 0.003 | 28 | 0.988 | 0.994 |
| aCA1 (L) | Scenes | Correct lures < repeats | -0.117 | 28 | 0.538 | 0.848 |
| aCA1 (L) | Scenes | Correct lures < incorrect lures | 0.029 | 28 | 0.878 | 0.952 |
| aCA1 (R) | Objects | Correct lures < repeats | 0.066 | 28 | 0.727 | 0.872 |
| aCA1 (R) | Objects | Correct lures < incorrect lures | -0.008 | 29 | 0.966 | 0.994 |

| Region | Stimulus | Contrast | Pearson's r | DF | Uncorrected p-value | Corrected p-value |
|---|---|---|---|---|---|---|
| aCA1 (R) | Scenes | Correct lures < repeats | 0.419 | 28 | 0.021 | 0.319 |
| aCA1 (R) | Scenes | Correct lures < incorrect lures | 0.118 | 28 | 0.536 | 0.868 |
| aCA3 (L) | Objects | Correct lures < repeats | 0.239 | 29 | 0.196 | 0.672 |
| aCA3 (L) | Objects | Correct lures < incorrect lures | 0.109 | 29 | 0.560 | 0.800 |
| aCA3 (L) | Scenes | Correct lures < repeats | 0.258 | 29 | 0.161 | 0.741 |
| aCA3 (L) | Scenes | Correct lures < incorrect lures | 0.418 | 29 | 0.019 | 0.219 |
| aCA3 (R) | Objects | Correct lures < repeats | 0.147 | 29 | 0.430 | 0.762 |
| aCA3 (R) | Objects | Correct lures < incorrect lures | 0.280 | 29 | 0.128 | 0.446 |
| aCA3 (R) | Scenes | Correct lures < repeats | -0.166 | 28 | 0.380 | 0.835 |
| aCA3 (R) | Scenes | Correct lures < incorrect lures | -0.058 | 29 | 0.757 | 0.908 |
| aDG (L) | Objects | Correct lures < repeats | 0.014 | 28 | 0.940 | 0.961 |
| aDG (L) | Objects | Correct lures < incorrect lures | -0.061 | 29 | 0.746 | 0.972 |
| aDG (L) | Scenes | Correct lures < repeats | 0.094 | 28 | 0.622 | 0.848 |
| aDG (L) | Scenes | Correct lures < incorrect lures | 0.416 | 29 | 0.020 | 0.219 |
| aDG (R) | Objects | Correct lures < repeats | -0.426 | 27 | 0.021 | 0.318 |
| aDG (R) | Objects | Correct lures < incorrect lures | -0.038 | 27 | 0.844 | 0.994 |
| aDG (R) | Scenes | Correct lures < repeats | 0.012 | 27 | 0.950 | 0.950 |
| aDG (R) | Scenes | Correct lures < incorrect lures | -0.108 | 27 | 0.577 | 0.868 |
| pSUB (L) | Objects | Correct lures < repeats | 0.016 | 28 | 0.931 | 0.961 |
| pSUB (L) | Objects | Correct lures < incorrect lures | -0.306 | 28 | 0.100 | 0.446 |
| pSUB (L) | Scenes | Correct lures < repeats | -0.120 | 28 | 0.527 | 0.848 |
| pSUB (L) | Scenes | Correct lures < incorrect lures | 0.138 | 29 | 0.459 | 0.868 |
| pSUB (R) | Objects | Correct lures < repeats | -0.517 | 28 | 0.003 | 0.103 |
| pSUB (R) | Objects | Correct lures < incorrect lures | -0.260 | 28 | 0.165 | 0.446 |
| pSUB (R) | Scenes | Correct lures < repeats | 0.098 | 28 | 0.606 | 0.848 |
| pSUB (R) | Scenes | Correct lures < incorrect lures | -0.172 | 28 | 0.362 | 0.851 |
| pCA1 (L) | Objects | Correct lures < repeats | 0.072 | 29 | 0.701 | 0.872 |
| pCA1 (L) | Objects | Correct lures < incorrect lures | 0.008 | 29 | 0.965 | 0.994 |
| pCA1 (L) | Scenes | Correct lures < repeats | 0.218 | 28 | 0.247 | 0.741 |
| pCA1 (L) | Scenes | Correct lures < incorrect lures | -0.044 | 28 | 0.817 | 0.943 |
| pCA1 (R) | Objects | Correct lures < repeats | -0.255 | 28 | 0.174 | 0.672 |
| pCA1 (R) | Objects | Correct lures < incorrect lures | -0.416 | 28 | 0.022 | 0.446 |
| pCA1 (R) | Scenes | Correct lures < repeats | 0.037 | 28 | 0.847 | 0.938 |

| Region | Stimulus | Contrast | Pearson's r | DF | Uncorrected p-value | Corrected p-value |
|---|---|---|---|---|---|---|
| pCA1 (R) | Scenes | Correct lures < incorrect lures | -0.266 | 28 | 0.156 | 0.529 |
| pCA3 (L) | Objects | Correct lures < repeats | 0.009 | 29 | 0.961 | 0.961 |
| pCA3 (L) | Objects | Correct lures < incorrect lures | -0.298 | 29 | 0.103 | 0.446 |
| pCA3 (L) | Scenes | Correct lures < repeats | -0.017 | 29 | 0.926 | 0.950 |
| pCA3 (L) | Scenes | Correct lures < incorrect lures | -0.023 | 29 | 0.903 | 0.952 |
| pCA3 (R) | Objects | Correct lures < repeats | 0.122 | 28 | 0.519 | 0.779 |
| pCA3 (R) | Objects | Correct lures < incorrect lures | 0.270 | 28 | 0.149 | 0.446 |
| pCA3 (R) | Scenes | Correct lures < repeats | -0.277 | 28 | 0.138 | 0.741 |
| pCA3 (R) | Scenes | Correct lures < incorrect lures | -0.250 | 28 | 0.183 | 0.550 |
| pDG (L) | Objects | Correct lures < repeats | -0.177 | 28 | 0.350 | 0.699 |
| pDG (L) | Objects | Correct lures < incorrect lures | -0.235 | 29 | 0.203 | 0.446 |
| pDG (L) | Scenes | Correct lures < repeats | 0.163 | 28 | 0.390 | 0.835 |
| pDG (L) | Scenes | Correct lures < incorrect lures | 0.306 | 28 | 0.100 | 0.529 |
| pDG (R) | Objects | Correct lures < repeats | -0.095 | 29 | 0.609 | 0.848 |
| pDG (R) | Objects | Correct lures < incorrect lures | -0.222 | 29 | 0.230 | 0.461 |
| pDG (R) | Scenes | Correct lures < repeats | 0.529 | 28 | 0.003 | 0.079 |
| pDG (R) | Scenes | Correct lures < incorrect lures | -0.105 | 28 | 0.581 | 0.868 |
| TAIL (L) | Objects | Correct lures < repeats | -0.142 | 31 | 0.432 | 0.762 |
| TAIL (L) | Objects | Correct lures < incorrect lures | -0.279 | 31 | 0.116 | 0.446 |
| TAIL (L) | Scenes | Correct lures < repeats | 0.316 | 31 | 0.073 | 0.547 |
| TAIL (L) | Scenes | Correct lures < incorrect lures | 0.093 | 31 | 0.608 | 0.868 |
| TAIL (R) | Objects | Correct lures < repeats | -0.225 | 31 | 0.208 | 0.672 |
| TAIL (R) | Objects | Correct lures < incorrect lures | -0.255 | 31 | 0.151 | 0.446 |
| TAIL (R) | Scenes | Correct lures < repeats | 0.119 | 31 | 0.508 | 0.848 |
| TAIL (R) | Scenes | Correct lures < incorrect lures | 0.068 | 31 | 0.705 | 0.881 |

**Notes. The prefix '*a'* denotes anterior hippocampus; '*p'*, posterior hippocampus. ALERC: anterolateral entorhinal cortex; AMY: amygdala; B: bilateral; BA: Brodmann area; CA: cornu ammonis; DG: dentate gyrus; L: left; PHC: parahippocampal cortex; PMERC: posterior-medial entorhinal cortex; R: right; SUB: subiculum.**

*6.3 Model selection procedure to identify unique contributions of MTL cortical and hippocampal subfield response magnitude and neural similarity to inter-individual variability in memory performance*

Greater left pDG univariate lure-related novelty responses, lower right pSUB pattern similarity for correct lures than repeats, and lower pCA1 activity during correct vs. incorrect lures predicted better object memory ($F(3, 26) = 7.75$, $p < .001$, ${}_{p}R^2 = 0.41$). Better scene memory was best explained by stronger lure-related novelty responses in right aDG, reduced lure-related novelty in left pCA3, and greater neural pattern similarity in right pSUB for correct lures over repeats ($F(3, 23) = 15.5$, $p<.001$, ${}_{p}R^2 = 0.67$).

These findings point to a direct behavioral relevance for hippocampal activation strength and pattern similarity for inter-individual variability in memory fidelity across visual domains. Importantly, stronger univariate responses and lower neural pattern similarity for correct lures are not always better: while greater lure-related novelty responses in DG consistently correlated with better object and scene performance, less pronounced novelty responses in CA1 and CA3, respectively, were also beneficial. Moreover, the opposite effect of pattern similarity on object and scene memory in the same region (right pSUB) suggests that different mechanisms may underpin successful lure discrimination in these two tasks.

**Supplementary Table 8. Results from a multiple regression model fitting procedure to identify the best combination of predictors for object and scene mnemonic discrimination, respectively.**

| **Object Mnemonic Discrimination [Corrected Hit Rate]** | | | | |
|---|---|---|---|---|
| **Model Term** | **Beta Estimate** | **Standard Error** | ***t*-value** | ***p*-value** |
| (Intercept) | −0.064 | 0.143 | −0.445 | 0.660 |
| Left pDG (Response Magnitude Correct Lure > Repeat) | 0.468 | 0.152 | 3.088 | 0.005 |
| Right pCA1 (Response Magnitude Correct > Incorrect Lure) | −0.342 | 0.165 | −2.068 | 0.049 |
| Right pSUB (Pattern Similarity Correct Lure > Repeat) | −0.357 | 0.161 | −2.225 | 0.035 |
| **Scene Mnemonic Discrimination [Corrected Hit Rate]** | | | | |
| (Intercept) | 0.123 | 0.128 | 0.963 | 0.345 |
| Left pCA3 (Response Magnitude Correct Lure > Repeat) | −0.447 | 0.163 | −2.750 | 0.011 |
| Right aDG (Response Magnitude Correct Lure > Repeat) | 0.989 | 0.161 | 6.145 | <0.001 |
| Right pSUB (Pattern Similarity Correct Lure > Repeat) | 0.617 | 0.148 | 4.159 | <0.001 |

Note: Model estimates are standardised. Models were selected based on a five-fold cross-validation, choosing the model with the lowest root mean squared error. aDG: anterior dentate gyrus; pCA1: posterior cornu ammonis-1; pCA3: posterior CA3; pDG: posterior dentate gyrus; pSUB: posterior subiculum. Note that the top 25 models for objects all included the dentate gyrus, while the top 25 for scenes included either the DG or the hippocampal tail (which is predominantly occupied by the DG subfield).

### 7. *Associations between univariate and multivariate measures of brain activity*

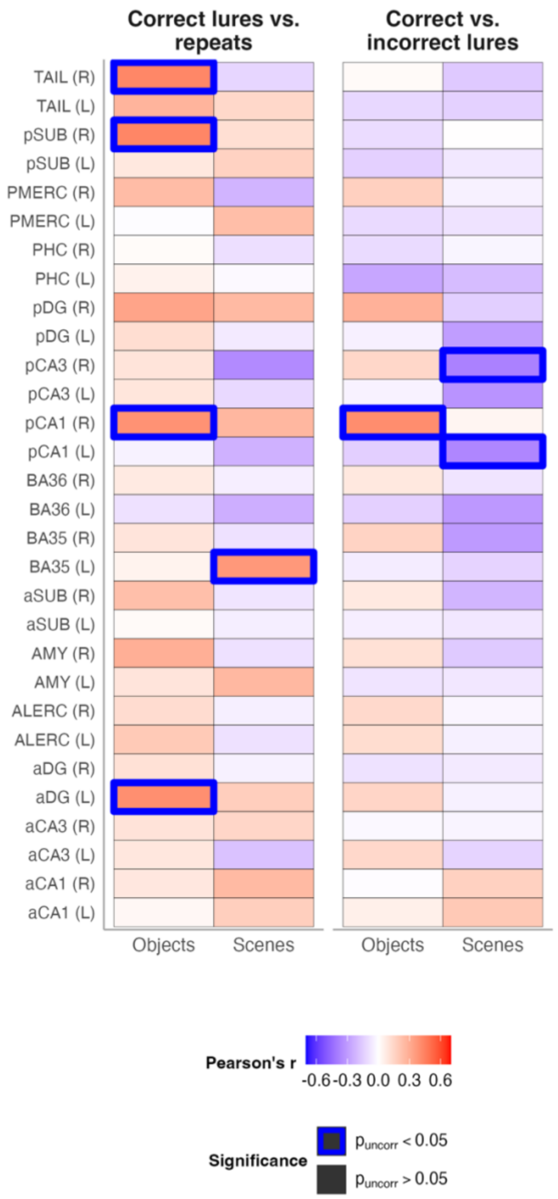


**Supplementary Figure 14. Overview of correlations between univariate and multivariate effects across all ROIs. *P*-values are FDR corrected at the level of contrasts and stimulus type. For univariate effects, positive values denote greater activity during correct lures vs. repeats or incorrect lures, respectively. For multivariate effects, positive effects refer to greater pattern dissimilarity between first targets and correct lures than between first targets and repeats or incorrect lures, respectively. Therefore, warm colours here point to a positive correlation indicative of greater activity during correct lures being associated with more distinct neural patterns during correct lures. This association here suggests that higher univariate activity in the respective ROI is linked to potential pattern separation-like responses in the same region. In contrast, cold colours suggest that greater activity during correct lures (compared to incorrect lures and repeats) is associated with higher pattern similarity, i.e. lower pattern dissimilarity, between first targets and correct lures compared to first targets and incorrect lures or repeats. Such a negative association suggests that higher univariate activity in the respective region is linked to potential pattern completion processes that are reinstating the neural pattern of previously encoded stimuli. Notes. The prefix '*a*' denotes anterior hippocampus; '*p*', posterior hippocampus. ALERC: anterolateral entorhinal cortex; AMY: amygdala; B: bilateral; BA: Brodmann area; CA: cornu ammonis; DG: dentate gyrus; L corr: correct rejection of the lure; L incorr: false alarm to lure; PHC: parahippocampal cortex; PMERC: posterior-medial entorhinal cortex; Rep: repetition of the target item; SUB: subiculum.**

**Supplementary Table 9. Statistics for correlations between univariate measures of response magnitude and multivariate measures of pattern similarity for contrasts and regions of interest. Univariate effects were contrast beta values from first-level analyses. Multivariate effects were differences in neural pattern similarity. Neural pattern similarity for repeats, correct lures, and incorrect lures here refers to the similarity between the baseline (first target presentations) and a given trial type. The correct lures < repeats contrast thus refers to lower pattern similarity between correct lures and firsts compared to pattern similarity between first and repeated targets.**

| Region | Stimulus | Contrast | Pearson's *r* | DF | Uncorrected *p*-value | Corrected *p*-value |
|---|---|---|---|---|---|---|
| ALERC (L) | Objects | Correct lures vs. repeats | 0.195 | 31 | 0.276 | 0.828 |
| ALERC (L) | Scenes | Correct lures vs. repeats | -0.088 | 30 | 0.632 | 0.836 |
| ALERC (L) | Objects | Correct vs. incorrect lures | 0.126 | 30 | 0.491 | 0.920 |
| ALERC (L) | Scenes | Correct vs. incorrect lures | -0.046 | 30 | 0.804 | 0.907 |
| ALERC (R) | Objects | Correct lures vs. repeats | 0.128 | 31 | 0.476 | 0.883 |
| ALERC (R) | Scenes | Correct lures vs. repeats | -0.046 | 30 | 0.801 | 0.858 |
| ALERC (R) | Objects | Correct vs. incorrect lures | 0.138 | 31 | 0.445 | 0.920 |
| ALERC (R) | Scenes | Correct vs. incorrect lures | -0.029 | 30 | 0.877 | 0.907 |
| AMY (L) | Objects | Correct lures vs. repeats | 0.098 | 31 | 0.586 | 0.883 |
| AMY (L) | Scenes | Correct lures vs. repeats | 0.259 | 30 | 0.152 | 0.656 |
| AMY (L) | Objects | Correct vs. incorrect lures | -0.079 | 30 | 0.665 | 0.920 |
| AMY (L) | Scenes | Correct vs. incorrect lures | -0.070 | 30 | 0.704 | 0.907 |
| AMY (R) | Objects | Correct lures vs. repeats | 0.289 | 31 | 0.103 | 0.516 |
| AMY (R) | Scenes | Correct lures vs. repeats | -0.090 | 30 | 0.623 | 0.836 |
| AMY (R) | Objects | Correct vs. incorrect lures | 0.110 | 30 | 0.549 | 0.920 |
| AMY (R) | Scenes | Correct vs. incorrect lures | -0.160 | 29 | 0.391 | 0.907 |
| BA35 (L) | Objects | Correct lures vs. repeats | 0.045 | 28 | 0.815 | 0.956 |
| BA35 (L) | Scenes | Correct lures vs. repeats | 0.370 | 27 | 0.048 | 0.656 |
| BA35 (L) | Objects | Correct vs. incorrect lures | -0.056 | 26 | 0.779 | 0.933 |
| BA35 (L) | Scenes | Correct vs. incorrect lures | -0.130 | 28 | 0.494 | 0.907 |
| BA35 (R) | Objects | Correct lures vs. repeats | 0.098 | 29 | 0.601 | 0.883 |
| BA35 (R) | Scenes | Correct lures vs. repeats | -0.089 | 28 | 0.641 | 0.836 |
| BA35 (R) | Objects | Correct vs. incorrect lures | 0.161 | 29 | 0.387 | 0.920 |
| BA35 (R) | Scenes | Correct vs. incorrect lures | -0.306 | 29 | 0.094 | 0.561 |
| BA36 (L) | Objects | Correct lures vs. repeats | -0.088 | 29 | 0.637 | 0.883 |
| BA36 (L) | Scenes | Correct lures vs. repeats | -0.244 | 28 | 0.194 | 0.656 |
| BA36 (L) | Objects | Correct vs. incorrect lures | -0.139 | 28 | 0.462 | 0.920 |
| BA36 (L) | Scenes | Correct vs. incorrect lures | -0.310 | 28 | 0.096 | 0.561 |

| Region | Stimulus | Contrast | Pearson's *r* | DF | Uncorrected *p*-value | Corrected *p*-value |
|---|---|---|---|---|---|---|
| BA36 (R) | Objects | Correct lures vs. repeats | 0.078 | 29 | 0.677 | 0.883 |
| BA36 (R) | Scenes | Correct lures vs. repeats | -0.049 | 28 | 0.797 | 0.858 |
| BA36 (R) | Objects | Correct vs. incorrect lures | 0.086 | 28 | 0.652 | 0.920 |
| BA36 (R) | Scenes | Correct vs. incorrect lures | -0.080 | 29 | 0.670 | 0.907 |
| PHC (L) | Objects | Correct lures vs. repeats | 0.049 | 29 | 0.795 | 0.956 |
| PHC (L) | Scenes | Correct lures vs. repeats | -0.016 | 28 | 0.931 | 0.931 |
| PHC (L) | Objects | Correct vs. incorrect lures | -0.268 | 28 | 0.152 | 0.920 |
| PHC (L) | Scenes | Correct vs. incorrect lures | -0.201 | 28 | 0.288 | 0.907 |
| PHC (R) | Objects | Correct lures vs. repeats | 0.016 | 29 | 0.933 | 0.956 |
| PHC (R) | Scenes | Correct lures vs. repeats | -0.094 | 28 | 0.622 | 0.836 |
| PHC (R) | Objects | Correct vs. incorrect lures | -0.108 | 28 | 0.571 | 0.920 |
| PHC (R) | Scenes | Correct vs. incorrect lures | -0.030 | 29 | 0.873 | 0.907 |
| PMERC (L) | Objects | Correct lures vs. repeats | -0.010 | 31 | 0.956 | 0.956 |
| PMERC (L) | Scenes | Correct lures vs. repeats | 0.238 | 30 | 0.189 | 0.656 |
| PMERC (L) | Objects | Correct vs. incorrect lures | -0.110 | 31 | 0.544 | 0.920 |
| PMERC (L) | Scenes | Correct vs. incorrect lures | -0.084 | 31 | 0.642 | 0.907 |
| PMERC (R) | Objects | Correct lures vs. repeats | 0.245 | 31 | 0.170 | 0.636 |
| PMERC (R) | Scenes | Correct lures vs. repeats | -0.226 | 30 | 0.214 | 0.656 |
| PMERC (R) | Objects | Correct vs. incorrect lures | 0.173 | 31 | 0.337 | 0.920 |
| PMERC (R) | Scenes | Correct vs. incorrect lures | -0.041 | 30 | 0.824 | 0.907 |
| TAIL (L) | Objects | Correct lures vs. repeats | 0.273 | 28 | 0.144 | 0.616 |
| TAIL (L) | Scenes | Correct lures vs. repeats | 0.142 | 28 | 0.453 | 0.836 |
| TAIL (L) | Objects | Correct vs. incorrect lures | -0.117 | 28 | 0.539 | 0.920 |
| TAIL (L) | Scenes | Correct vs. incorrect lures | -0.137 | 28 | 0.471 | 0.907 |
| TAIL (R) | Objects | Correct lures vs. repeats | 0.427 | 29 | 0.017 | 0.252 |
| TAIL (R) | Scenes | Correct lures vs. repeats | -0.122 | 28 | 0.519 | 0.836 |
| TAIL (R) | Objects | Correct vs. incorrect lures | 0.018 | 28 | 0.923 | 0.955 |
| TAIL (R) | Scenes | Correct vs. incorrect lures | -0.161 | 29 | 0.388 | 0.907 |
| aCA1 (L) | Objects | Correct lures vs. repeats | 0.028 | 28 | 0.883 | 0.956 |
| aCA1 (L) | Scenes | Correct lures vs. repeats | 0.176 | 27 | 0.361 | 0.833 |
| aCA1 (L) | Objects | Correct vs. incorrect lures | 0.056 | 28 | 0.767 | 0.933 |
| aCA1 (L) | Scenes | Correct vs. incorrect lures | 0.199 | 27 | 0.301 | 0.907 |
| aCA1 (R) | Objects | Correct lures vs. repeats | 0.086 | 28 | 0.650 | 0.883 |

| Region | Stimulus | Contrast | Pearson's *r* | DF | Uncorrected *p*-value | Corrected *p*-value |
|---|---|---|---|---|---|---|
| aCA1 (R) | Scenes | Correct lures vs. repeats | 0.253 | 27 | 0.186 | 0.656 |
| aCA1 (R) | Objects | Correct vs. incorrect lures | -0.007 | 27 | 0.969 | 0.969 |
| aCA1 (R) | Scenes | Correct vs. incorrect lures | 0.168 | 28 | 0.374 | 0.907 |
| aCA3 (L) | Objects | Correct lures vs. repeats | 0.089 | 28 | 0.641 | 0.883 |
| aCA3 (L) | Scenes | Correct lures vs. repeats | -0.183 | 27 | 0.341 | 0.833 |
| aCA3 (L) | Objects | Correct vs. incorrect lures | 0.141 | 28 | 0.457 | 0.920 |
| aCA3 (L) | Scenes | Correct vs. incorrect lures | -0.129 | 28 | 0.497 | 0.907 |
| aCA3 (R) | Objects | Correct lures vs. repeats | 0.104 | 28 | 0.586 | 0.883 |
| aCA3 (R) | Scenes | Correct lures vs. repeats | 0.150 | 28 | 0.430 | 0.836 |
| aCA3 (R) | Objects | Correct vs. incorrect lures | -0.021 | 28 | 0.912 | 0.955 |
| aCA3 (R) | Scenes | Correct vs. incorrect lures | -0.033 | 28 | 0.862 | 0.907 |
| aDG (L) | Objects | Correct lures vs. repeats | 0.401 | 28 | 0.028 | 0.252 |
| aDG (L) | Scenes | Correct lures vs. repeats | 0.179 | 27 | 0.353 | 0.833 |
| aDG (L) | Objects | Correct vs. incorrect lures | 0.152 | 27 | 0.430 | 0.920 |
| aDG (L) | Scenes | Correct vs. incorrect lures | -0.041 | 27 | 0.834 | 0.907 |
| aDG (R) | Objects | Correct lures vs. repeats | 0.111 | 27 | 0.566 | 0.883 |
| aDG (R) | Scenes | Correct lures vs. repeats | -0.041 | 26 | 0.836 | 0.864 |
| aDG (R) | Objects | Correct vs. incorrect lures | -0.086 | 27 | 0.658 | 0.920 |
| aDG (R) | Scenes | Correct vs. incorrect lures | -0.064 | 26 | 0.748 | 0.907 |
| aSUB (L) | Objects | Correct lures vs. repeats | 0.018 | 28 | 0.923 | 0.956 |
| aSUB (L) | Scenes | Correct lures vs. repeats | -0.051 | 27 | 0.793 | 0.858 |
| aSUB (L) | Objects | Correct vs. incorrect lures | -0.049 | 27 | 0.799 | 0.933 |
| aSUB (L) | Scenes | Correct vs. incorrect lures | -0.070 | 28 | 0.712 | 0.907 |
| aSUB (R) | Objects | Correct lures vs. repeats | 0.233 | 28 | 0.215 | 0.716 |
| aSUB (R) | Scenes | Correct lures vs. repeats | -0.077 | 27 | 0.690 | 0.858 |
| aSUB (R) | Objects | Correct vs. incorrect lures | 0.080 | 28 | 0.675 | 0.920 |
| aSUB (R) | Scenes | Correct vs. incorrect lures | -0.223 | 28 | 0.236 | 0.907 |
| pCA1 (L) | Objects | Correct lures vs. repeats | -0.040 | 28 | 0.835 | 0.956 |
| pCA1 (L) | Scenes | Correct lures vs. repeats | -0.236 | 27 | 0.219 | 0.656 |
| pCA1 (L) | Objects | Correct vs. incorrect lures | -0.144 | 27 | 0.456 | 0.920 |
| pCA1 (L) | Scenes | Correct vs. incorrect lures | -0.366 | 28 | 0.047 | 0.561 |
| pCA1 (R) | Objects | Correct lures vs. repeats | 0.389 | 28 | 0.034 | 0.252 |
| pCA1 (R) | Scenes | Correct lures vs. repeats | 0.262 | 27 | 0.170 | 0.656 |

| Region | Stimulus | Contrast | Pearson's *r* | DF | Uncorrected *p*-value | Corrected *p*-value |
|---|---|---|---|---|---|---|
| pCA1 (R) | Objects | Correct vs. incorrect lures | 0.406 | 28 | 0.026 | 0.775 |
| pCA1 (R) | Scenes | Correct vs. incorrect lures | 0.035 | 28 | 0.854 | 0.907 |
| pCA3 (L) | Objects | Correct lures vs. repeats | 0.093 | 28 | 0.625 | 0.883 |
| pCA3 (L) | Scenes | Correct lures vs. repeats | -0.112 | 27 | 0.562 | 0.836 |
| pCA3 (L) | Objects | Correct vs. incorrect lures | -0.039 | 28 | 0.840 | 0.933 |
| pCA3 (L) | Scenes | Correct vs. incorrect lures | -0.325 | 28 | 0.080 | 0.561 |
| pCA3 (R) | Objects | Correct lures vs. repeats | 0.100 | 28 | 0.598 | 0.883 |
| pCA3 (R) | Scenes | Correct lures vs. repeats | -0.355 | 27 | 0.059 | 0.656 |
| pCA3 (R) | Objects | Correct vs. incorrect lures | 0.145 | 28 | 0.443 | 0.920 |
| pCA3 (R) | Scenes | Correct vs. incorrect lures | -0.381 | 28 | 0.038 | 0.561 |
| pDG (L) | Objects | Correct lures vs. repeats | 0.122 | 28 | 0.520 | 0.883 |
| pDG (L) | Scenes | Correct lures vs. repeats | -0.062 | 27 | 0.751 | 0.858 |
| pDG (L) | Objects | Correct vs. incorrect lures | -0.044 | 27 | 0.822 | 0.933 |
| pDG (L) | Scenes | Correct vs. incorrect lures | -0.296 | 28 | 0.112 | 0.561 |
| pDG (R) | Objects | Correct lures vs. repeats | 0.328 | 28 | 0.077 | 0.460 |
| pDG (R) | Scenes | Correct lures vs. repeats | 0.252 | 27 | 0.187 | 0.656 |
| pDG (R) | Objects | Correct vs. incorrect lures | 0.282 | 28 | 0.131 | 0.920 |
| pDG (R) | Scenes | Correct vs. incorrect lures | -0.144 | 27 | 0.455 | 0.907 |
| pSUB (L) | Objects | Correct lures vs. repeats | 0.085 | 28 | 0.655 | 0.883 |
| pSUB (L) | Scenes | Correct lures vs. repeats | 0.166 | 27 | 0.389 | 0.833 |
| pSUB (L) | Objects | Correct vs. incorrect lures | -0.139 | 27 | 0.473 | 0.920 |
| pSUB (L) | Scenes | Correct vs. incorrect lures | -0.067 | 28 | 0.725 | 0.907 |
| pSUB (R) | Objects | Correct lures vs. repeats | 0.430 | 28 | 0.018 | 0.252 |
| pSUB (R) | Scenes | Correct lures vs. repeats | 0.119 | 27 | 0.540 | 0.836 |
| pSUB (R) | Objects | Correct vs. incorrect lures | -0.105 | 27 | 0.590 | 0.920 |
| pSUB (R) | Scenes | Correct vs. incorrect lures | 0.004 | 28 | 0.984 | 0.984 |

**Notes. '*a*' denotes anterior, '*p*' posterior hippocampus. ALERC: anterolateral entorhinal cortex; AMY: amygdala; B: bilateral; BA: Brodmann area; CA: cornu ammonis; DG: dentate gyrus; L: left; PHC: parahippocampal cortex; PMERC: posterior-medial ERC; R: right; SUB: subiculum.**
